# A draft Arab pangenome reference

**DOI:** 10.1101/2024.07.09.602638

**Authors:** Nasna Nassir, Mohamed A. Almarri, Muhammad Kumail, Nesrin Mohamed, Bipin Balan, Shehzad Hanif, Maryam AlObathani, Bassam Jamalalail, Hanan Elsokary, Dasuki Kondaramage, Suhana Shiyas, Noor Kosaji, Dharana Satsangi, Madiha Hamdi Saif Abdelmotagali, Ahmad Abou Tayoun, Olfat Zuhair Salem Ahmed, Douaa Fathi Youssef, Hanan Al Suwaidi, Ammar Albanna, Stefan Du Plessis, Hamda Hassan Khansaheb, Alawi Alsheikh-Ali, Mohammed Uddin

**Affiliations:** Center for Applied and Translational Genomics (CATG), Mohammed Bin Rashid University of Medicine and Health Sciences, Dubai Health, Dubai, UAE; College of Medicine, Mohammed Bin Rashid University of Medicine and Health Sciences, Dubai Health, Dubai, UAE; Genome Center, Department of Forensic Science and Criminology, Dubai Police GHQ, Dubai, UAE; Manipal Centre for Biotherapeutics Research, Manipal Academy of Higher Education, Manipal, Karnataka, India; Primary Health Care Services Sector, Dubai Health, Dubai, UAE; Al Jalila Genomics Center of Excellence, Al Jalila Children’s Specialty Hospital, Dubai Health, Dubai, UAE; Center for Genomic Discovery, Mohammed Bin Rashid University of Medicine and Health Sciences, Dubai Health, Dubai, UAE; AlAmal Psychiatric Hospital, UAE; Dubai Health, Dubai, UAE; GenomeArc Inc., Mississauga, ON, Canada

## Abstract

Pangenomes represent a significant shift from relying on a single reference sequence to a robust set of assemblies, Arab populations remain significantly underrepresented; hence, we present the first Arab Pangenome Reference (APR) utilizing 53 individuals of diverse Arab ethnicities. We assembled nuclear and mitochondrial pangenomes using 35.27X high-fidelity long reads, 54.22X ultralong reads and 65.46X Hi-C reads yielded contiguous haplotype-phased *de novo* assemblies of exceptional quality, with an average N50 of 124.28 Mb. We discovered 111.96 million base pairs of novel euchromatic sequences absent from existing human pangenomes, the T2T-CHM13, GRCh38 reference human genomes, and other public datasets. We identified 8.94 million population-specific small variants and 235,195 structural variants within the Arab pangenome. We detected 883 gene duplications including 15.06% associated with recessive diseases and 1,436 bp of novel mitochondrial pangenome sequence. Our study provides a valuable resource for future genomic medicine initiatives in Arab population and other global populations.

## Introduction

The vast complexity and diversity of the human genome has opened new avenues for research in genomics, leading to significant advances in our understanding of human biology and disease. However, the application of these genomic advances is limited by the quality, completeness, and representativeness of the reference genome used.^1^ In the pursuit of understanding the intricate tapestry of human genome variation, large-scale sequencing projects have provided invaluable insights.^2,3^ Recently, due to significant advancements in long-read genome technologies, the first complete telomere-to-telomere (T2T) sequence of the haploid human genome CHM13^4^ and the complete sequence of the Y chromosome were obtained.^5^ This finding is a remarkable achievement that fills 8% of the gaps that exist in the current reference genome GRCh38^4^ Although this sequence represents the first complete genome, it does not reflect human sequence diversity, and a population-wide approach is necessary to detect population-specific variants, sequences and regulatory elements. The Human Pangenome Reference Consortium (HPRC) has made significant strides in genomic research by constructing a pangenome from 47 ethnically diverse samples, adding 119 million base pairs and identifying 1,115 gene duplications absent in the GRCh38 reference genome.^6^ This finding led to a 104% increase in structural variant detection per haplotype.^6^ A recent study in China of 58 samples from 36 ethnic minorities identified 5.9 million small variants and 34,223 structural variants not found in the HPRC pangenome.^7^ These findings underscore the value of incorporating genetically diverse individuals for a comprehensive understanding of the human genome landscape.

Arabs constitute culturally diverse communities primarily from Middle East and North Africa (MENA) regions with a combined population of nearly 500 million, comprising approximately 6% of the global population. Unfortunately, Arab populations lack adequate representation in large-scale sequencing projects (i.e., the gnomAD database); neither the HPRC pangenome nor the 1000 Genomes Project include samples from this demographic.^8^ The ancestral complexities between different Arab ethnic backgrounds have not yet been fully understood through large-scale genomic initiatives.^9,10^ Moreover, the Arab population, which experiences a higher incidence of consanguineous marriages, is predisposed to an increased rate of rare recessive disorders.^11–13^ The incidence of diabetes, heart disease and cancer in Arab populations is increasing.^14,15^ The lack of reference genomes for Arab populations has limited the investigation of genetic diversity and the genetic underpinning of numerous diseases. Population-specific reference pangenomes will enable the identification of variants associated with diseases and sequences that are unique or prevalent in Arab populations.

In this study, we sought to generate a comprehensive Arab Pangenome Reference (APR) using multiple sequencing technologies and *de novo* assembly construction approaches. Here, we present the preliminary construction of the APR from 53 human genomes with high-quality *de novo* diploid phased assemblies and their comparison with reference genomes and genomic datasets. Our analysis also included the application of the APR in functional annotation, short and long-read whole-genome mapping for variant detection. We anticipate that our efforts will produce a pangenome that will allow for population-specific genome interpretation, facilitating more accurate genome interpretation with fewer biases and gaps, which will enhance the precision of detecting both structural and small variants.

### Healthy Arab sample cohort

We constructed a pangenome reference for healthy Arab individuals from eight geographical locations (the United Arab Emirates (UAE), Saudi Arabia, Oman, Jordan, Egypt, Morocco, Syria and Yemen) across the MENA region (Fig. 1a). 53 samples, including 50 unrelated adults (18 years or older) and a trio of family members with no known rare or common chronic diseases (i.e., no history of hypertension, diabetes mellitus, cancer, or lung or heart disease), were enrolled (Supplementary Fig. 1, Supplementary Table 1).

**Fig. 1:**
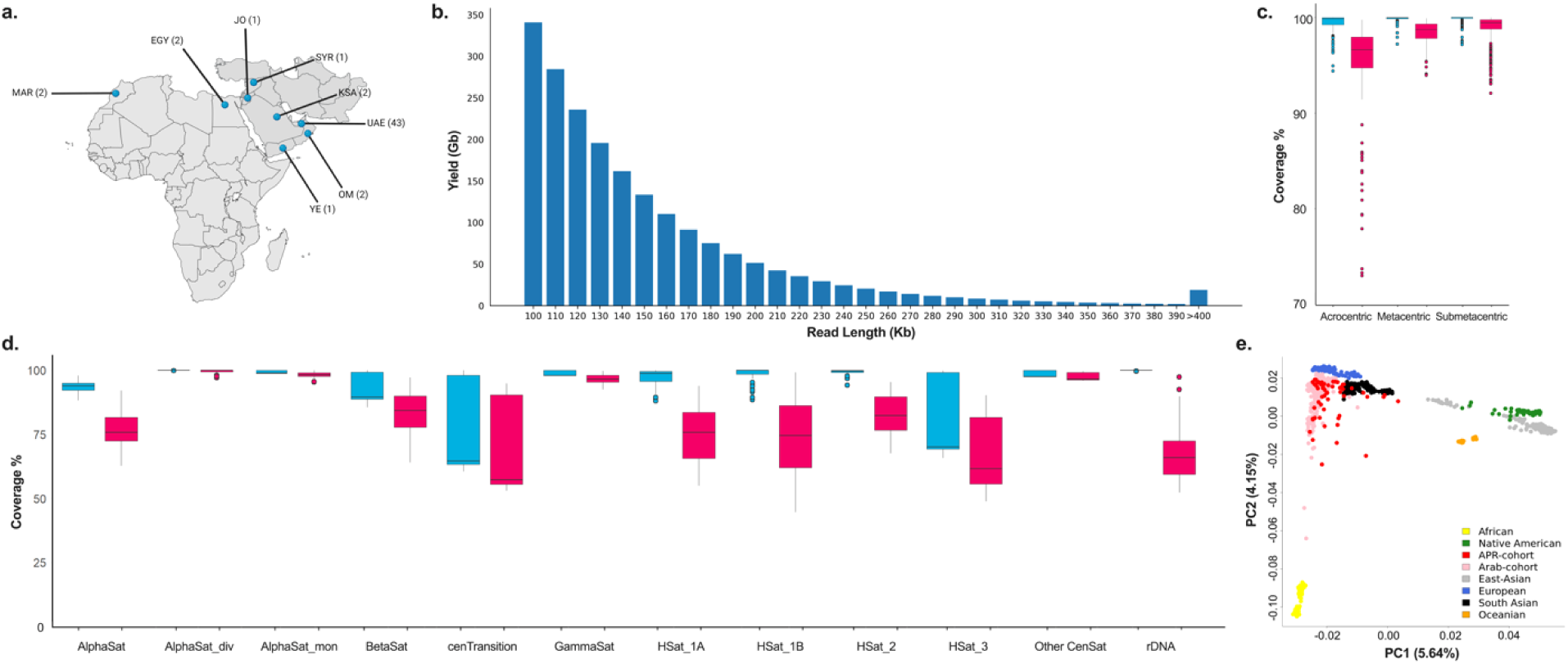
Cohort characterization and sequencing quality. **a,** Geographic diversity of sample collection. A map highlighting the distribution of sample collection sites. Each point marks the geographical location of cohorts involved in the study, illustrating the wide range recruitment strategy employed to capture genetic diversity. **b,** Ultra-long read yield from ONT sequencing. Histogram depicting the yield of ultra-long reads (>100 kb) produced by ONT sequencing. The x-axis classifies the reads based on length intervals, while the y-axis displays the yield for each category. **c,** Chromosome mapping distribution of sequencing reads. Boxplot presenting the distribution of ONT and PacBio reads that align to acrocentric, metacentric, and submetacentric chromosomes. Neon pink and teal blue bars represent PacBio and ONT data respectively. **d,** Boxplot illustrating the coverage of subtelomeric and pericentric regions by both ONT and PacBio reads. **e,** Population ethnicity variance using PCA. Two-dimensional scatter plot derived from Principal Component Analysis (PCA), visualizing the genetic variance among different ethnicities. Samples from Human Genome Diversity Project (HGDP) and Human Origins database are color coded by ethnicity, with APR highlighted in red. The two axes indicate the variance captured by first (PCA1) and second (PCA2) principal component, respectively.

### Assessment of sequencing quality and variant statistics

The genomes were sequenced using Pacific Biosciences (PacBio) high-fidelity (HiFi), Oxford Nanopore Technologies (ONT) ultralong read sequencing (ULK) and high-coverage (Hi-C) Illumina short-read sequencing methods. The average coverage for each sample was 35.27X (26.60X–69.39X) using PacBio HiFi sequencing, 54.22X (33.33X–90.87X) using ONT ultralong read sequencing and 65.46X (50.89X–91.96X) using Hi-C sequencing (Supplementary Fig. 2).

The average median Q score, indicating sequencing quality, was 33.19 (28-36) for PacBio HiFi sequencing, 17.35 (16-18.2) for ONT sequencing and 37 for Hi-C. The average N50 values were 16.69 kb (14.23 kb–19.06 kb) and 57.67 (43.64 kb–87.02 kb) for PacBio HiFi and ONT ultralong read sequencing, respectively (Supplementary Fig. 2 and Supplementary Table 2). Notably, ultralong reads exceeding 100 kb contributed to a substantial yield of 2011.14 Gb, translating to 670.38X coverage (an average of 12.53X per sample) (Fig. 1b, Supplementary Table 3). When mapped against CHM13 v2.0, both PacBio HiFi and ONT reads provided valuable insights into the sequencing coverage across acrocentric and metacentric chromosomes. Due to the use of ultralong protocols, the ONT reads exhibited better mapping across all chromosomes than did the PacBio reads, particularly for the acrocentric chromosomes, where 99.49% (94.95%-100%) coverage was achieved by ONT in comparison to 95.60% (91.61%- 99.75%) coverage achieved by PacBio (Fig. 1c, Supplementary Fig. 3, Supplementary Table 4a). We utilized both read types to analyze coverage across diverse regions of the human genome, including satellite DNA, centromeric transitions, and ribosomal DNA. Both platforms exhibited excellent coverage, with regions such as AlphaSat_div, AlphaSat_mon, GammaSat, and Other CenSat exhibiting average coverage above 96% (96.67%-100.00%). However, the ONT platform demonstrated a marked increase in rDNA coverage, with 99.93% (99.57%-100%) and 67.44% (52.25%-97.25%) coverage for the ONT and PacBio HiFi reads, respectively (Fig. 1d, Supplementary Table 4b).

Our joint calling analysis of PacBio HiFi data incorporating the DeepVariant pipeline (GRCh38) revealed an average of 4.21 million (M) (ranged from 4.02 M–4.48 M) single-nucleotide variants (SNVs), 913,786 (800,438-971,731) insertion/deletions (indels) and 53,618 (51,732-56,573) structural variants (SVs) per sample (Supplementary Table 5, Supplementary Fig. 4). We identified population-specific variants, including 1.40 M SNVs and 308,803 M indels, that were not previously reported in the 1000 Genomes Project,^2^ dbSNP,^16^ gnomAD^17^ or GME.^18^ We identified 158,696 population-specific structural variants not previously reported in the 1000 Genomes Project or Database of Genomic Variants (DGV).^19^

### Population structure of the APR samples

We subsequently analyzed the genetic structure of 53 APR genomes from published global and regional populations.^9,20–22^ Principal component analysis (PCA) demonstrated distinct clustering patterns of the APR samples and Arab groups (Fig. 1e), reflecting their unique population history and diversity in contrast to other global populations (Supplementary Fig. 5a, b and Supplementary Table 6a). Using the relatedness analysis using plink, we confirmed all APR samples are unrelated except for the trio (Supplementary Table 6b). Haplotype sharing information obtained using the fineSTRUCTURE algorithm^23^ demonstrated the substructures within our dataset, reflecting the different Arab subpopulations and the unrelated status of all 50 samples (Supplementary Fig. 5c). Model-based clustering using the ADMIXTURE algorithm, which integrated the genomic data from different Arab groups, indicated that the ancestry of the APR cohort generally reflects that of various Arab subpopulations (Supplementary Fig. 6).

Our analysis revealed that the APR cohort exhibited mitochondrial and Y chromosome haplogroup patterns closely resembling those associated with Arab populations. Specifically, the hierarchical clustering heatmaps demonstrated a significant alignment in haplogroup frequencies between our cohort and reference populations from Arab regions (Supplementary Fig. 7, Supplementary Table 7).

### Assembly construction and evaluation of diverse Arab genomes

Our comparative analysis of the APR trio revealed that both the Hifiasm^24^ and Verkko^25^ assemblers yielded comparable contig lengths (Supplementary Table 8). In samples lacking parental genome data, Hifiasm exhibited better performance in terms of both contig length and overall contiguity. Consequently, we performed *de novo* assembly using Hifiasm v0.19.5-r603 for our entire cohort and used those assemblies for downstream analysis. Sequence quality control was performed using Kraken,^26^ which filtered out non-human eukaryotic pathogen genomes and retained reads classified as human. We excluded an additional 109 contigs that mapped to ChrM with at least 95% identity (Supplementary Table 9). Additionally, we identified and cataloged 185 interchromosomal joins in the assemblies based on their sequence identity using paftools^27^ (Supplementary Fig. 8, Supplementary Table 10), resulting in the exclusion of four contigs due to significantly large mis-joins in acrocentric chromosomes. Inspector assembly polishing^28^ was further carried out to reduce small-scale assembly errors.

The resulting assemblies yielded an average genome size of 3.01 Gb (2.86 Gb–3.11 Gb) (Supplementary Fig. 9, Supplementary Table 11), with more than 83.02% of assemblies surpassing the GRCh38 benchmark genome size of 2.94 Gb. On average, the assemblies comprised 103 contigs (ranged from 58 to 245), with a contig N50 length of 124.28 Mb (86.26 Mb–146.15 Mb) (Fig. 2a) and an average auN of 122.48 Mb (87.77 Mb–146.00 Mb) (Supplementary Fig. 10a-b). This value exceeds the GRCh38 contig N50 of 57.88 Mb, demonstrating enhanced contiguity across all our assemblies. All samples surpassed the 40 Mb N50 of the HPRC reference genome (Supplementary Fig. 10c). The average QV of our assemblies, computed using yak v0.1-r66, was 57.53 (52.85–62.59), indicating robust assembly quality (Fig. 2b). We compared our assemblies to the CHM13 and GRCh38 reference genomes and revealed an average coverage of 93.59% (92.79% for males, 94.24% for females) and 96.53% (95.49% for males, 97.39% for females), respectively (Fig. 2c). We identified an average of 49.86 Mb (29.31 Mb-73.14 Mb) and 76.81 Mb (54.74 Mb-108.31 Mb) of assembled contigs that did not align with CHM13 and GRCh38, respectively (Fig. 2d, Supplementary Fig. 11 a-c). This observation is consistent with the unmapped rates reported in diverse pangenome studies, reflecting the complexity of the genome assembly and alignment processes. The duplication ratio, indicative of read mapping multiplicity against reference genomes, averaged 1.03 (1.023- 1.047) and 1.012 (1.005-1.022) for GRCh38 and CHM13, respectively (Supplementary Fig. 11d), highlighting the precision of our assembly with minimal duplicated regions.

**Fig. 2:**
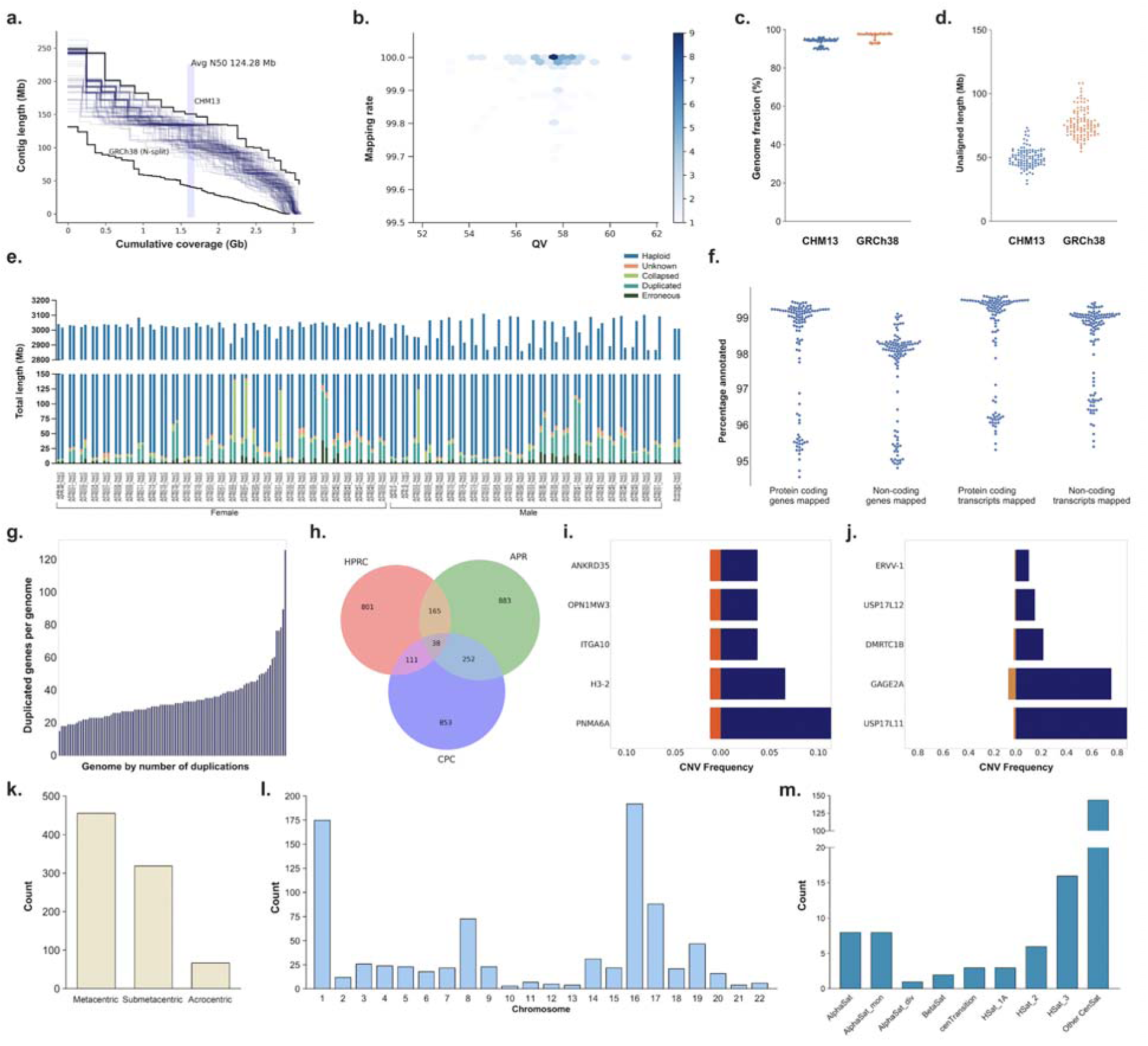
Quality assessment of 53 phased diploid assemblies and gene duplication analysis. **a,** Assembly contiguity. A line graph showing contig length plotted against cumulative assembly coverage, with reference contiguities for both CHM13 and GRCh38 genomes included for comparison. **b,** Assembly accuracy and completeness. A plot illustrating the mapping rate versus consensus accuracy (QV), offering insights into the completeness and accuracy of the assemblies. **c,** Genome fraction. Scatter plot showing the fraction of the genome covered by the assemblies compared to benchmark references CHM13 and GRCh38. **d,** Unaligned length. A scatter plot comparing the unaligned length of the assemblies relative to the CHM13 and GRCh38 references. **e,** Flagger analysis. Bar chart illustrating the reliability of 53 APR assemblies using read mapping. The plot differentiates between paternal and maternal haplotypes, with regions flagged as reliable (blue) representing the majority of each assembly. The y-axis is broken to emphasize the dominant reliable haploid component and the stratification of the unreliable blocks. **f,** Gene and transcript annotation. Scatter plot showing the percentages of protein-coding and noncoding genes, as well as transcripts annotated from the reference set in each of the assemblies. **g,** Gene duplication per assembly. Histogram presenting the number of unique duplicated gene families in each phased assembly in comparison to the number of duplicated genes annotated in GRCh38. **h,** Comparative duplicated gene analysis. Venn diagram visualizing the overlap and unique counts of duplicated genes across APR, HPRC, and CPC assemblies. **i,** Arab-HPRC duplicated gene overlap. Bar graph showcasing five overlapped duplicated genes with a higher frequency (≥5%) in Arab assemblies (blue) compared to HPRC (orange). **j,** Arab-CPC duplicated gene overlap. Bar chart illustrating five overlapped duplicated genes with a significantly higher frequency (≥5%) in Arab assemblies (blue) in contrast to CPC (yellow). **k,** Bar graphs indicating the count of APR unique duplicated genes across chromosome types: acrocentric, metacentric, and submetacentric. **l,** Bar graph showing the count of APR unique duplicated genes dispersed across all individual chromosomes, highlighting regions of enrichment. **m,** Gene duplication in microsatellite region. Bar graph depicting the count of APR unique duplicated genes located in microsatellite regions.

To evaluate the reliability of our assembled genomic sequences, we employed the Flagger tool,^29^ which is designed to identify misassemblies within a phased diploid assembly. Our analysis using Flagger revealed that a minimal proportion (1.28% or 37.91 Mb) of the total assembly was deemed unreliable (Fig. 2e, Supplementary Table 12), indicating the high accuracy of the APR assemblies.

The phasing accuracy of the trio samples was evaluated using yak trioeval, revealing a switch error rate and Hamming error of 0.18% and 0.23%, respectively. For the samples with Hi-C data, the phasing accuracy was computed using pstools,^30^ with an average switch error rate and Hamming error of 0.64% (0.21%-1.07%) and 1.96% (0.76%-3.21%), respectively (Supplementary Table 13a). To evaluate the mapping accuracy of our assembled contigs within the most complex centromeric regions, we employed UniAligner^31^ to calculate the mapping percentages at 100 bp intervals across these regions. Our analysis revealed the average mapping rate of 26.97% for APR (Supplementary Fig. 12), compared to 20.42% for HPRC, highlighting the accuracy and reliability of APR assemblies in representing complex genomic regions such as centromeres. Scaffolds were categorized at the chromosome level using strict criteria, requiring that each scaffold predominantly map to a single chromosome with a maximum of three gaps and cover a significant portion of the chromosome’s length, excluding regions composed of microsatellites. Applying these criteria, we identified 417 chromosome-scale scaffolds in the APR genome assemblies (Supplementary Table 13b). These comprehensive quality control assessments verified the high contiguity and accuracy of our *de novo* assembled genomes, providing a solid foundation for downstream genomic and clinical applications.

### Gene duplications in APR assemblies

To investigate gene duplication events within APR diploid assemblies, we employed the Liftoff v1.6.3^32^ tool and annotated the genes using GENCODE GRCh38.p14 (GENCODE release 38). The identified gene duplications revealed novel and diverse genomic features of the Arab population. Remarkably, a median of 99.04% (94.52%-99.47%) of protein-coding genes and 99.28% (95.29%-99.63%) of protein-coding transcripts were identified across all the APR assemblies (Fig. 2f). Each genome had an average of 35 genes with a gain in copy number relative to GRCh38 (Fig. 2g, Supplementary Table 14). We observed that 37.83% of duplicated genes were present as singletons, whereas 8.17% were common (>5%) among APR assemblies.

We performed a comparative assessment of gene duplications within the APR, HPRC, and CPC assemblies to identify genes that were specific to the Arab population. We identified 1135 duplicated genes unique to the APR assemblies that were absent from the HPRC assemblies and a further 883 duplicated genes that were absent from both the HPRC and CPC assemblies (Fig. 2h). In addition, 38 duplicated genes from the APR assemblies were also observed in the HPRC and CPC assemblies. Among these overlapping genes, some genes were present at a greater frequency (≥5%) in the APR assemblies than in the HPRC and CPC assemblies (Fig. 2i, 2j), such as the *USP17L* genes. The *USP17L* genes encode ubiquitin-specific proteases, which are involved in various cellular processes, such as protein degradation, DNA repair, and cell cycle regulation.^33,34^ Duplicated genes absent in APR but present in both the HPRC and CPC assemblies were involved in the olfactory signaling (p<3.39x10^-20^) and G alpha signaling (p<1.82x10^-11^) pathways (Supplementary Table 15).

Among the unique APR duplicated genes, the *TAF11L5* gene found to be duplicated across all the APR assemblies but was absent in the HPRC and CPC assemblies. The *TAF11L5* gene encodes a TATA-box binding protein (TBP)-associated factor, which is predicted to be involved in the assembly of the RNA polymerase II preinitiation complex, a large complex of proteins that initiates the transcription of protein-coding genes.^35^ The unique APR duplicated genes were significantly enriched in the oxidative phosphorylation (p<9.41x10^-09^) and ribonucleotide metabolism (p<9.41x10^-09^) pathways (Supplementary Table 16). These unique gene duplications were predominantly located on submetacentric chromosomes (Fig. 2k), primarily chromosomes 16 and 1 (Fig. 2l). Conversely, the acrocentric chromosomes had a low number of duplication events. This observation may be attributed to the inherent challenges in resolving these regions. Notably, unique gene duplications were present in microsatellite regions (191 out of 883), primarily within centromeric satellites (Fig. 2m, Supplementary Table 17). Furthermore, 15.06% of unique APR duplicated genes were implicated in recessive conditions, compared to 14.61% of unique HPRC genes and 11.25% of unique CPC genes (Supplementary Table 17). Using the Horizon platform, 1.70% of unique duplicated genes were annotated as pan-ethnic disease genes associated with severe phenotypes^36^ (Supplementary Table 18).

Recurrent gene duplications specific to APR assemblies further highlight the unique genetic landscape, with 255 and 123 duplicated genes not found in the HPRC and CPC assemblies, respectively (Supplementary Fig. 13). In addition, 13 duplicated genes from the APR assemblies were also observed in the HPRC and CPC assemblies. These unique recurrent duplicated genes in the APR were enriched in pathways related to odorant binding (p<9.54x10^-17^), defensin production (p<5.60x10^-07^) and antimicrobial peptide activity (p<5.95x10^-05^) (Supplementary Table 19).

### Pangenome graph construction and Arab genome-specific variants

We constructed a pangenome graph incorporating 53 Arab genomes using Minigraph-Cactus (v2.7.2),^37^ integrating 106 long-read assemblies into a graph structure. The graph was seeded with the CHM13 (as the backbone) and GRCh38 reference genomes and expanded using Minigraph to incorporate the new assemblies. The resulting pangenome encompassed a total length of 3,330,235,835 bp, with 80,141,431 nodes and 110,455,287 edges, whereas the HPRC pangenome has a total length of 3,328,787,872 bp (Supplementary Table 20, Supplementary Fig. 14).

We conducted a comparative analysis of the Arab pangenome graph with the combined HPRC and CPC pangenome graph, which represents a diverse set of human genomes from different populations. Each genome in our cohort contained an average of 5.45 M (5.11 M-5.99 M) small variants (Fig. 3a, Supplementary Table 21). Our analysis revealed an average of 899,585 (782,486-1,077,439) unique small variants per sample (Fig. 3b) and 13.70 M small variants that were unique to APR (Fig. 3c) compared with HPRC-CPC, CHM13 and GRCh38. Furthermore, we identified 8.94 M population-specific small variants absent in the HPRC-CPC pangenomes, CHM13, GRCh38 and variant databases, including dbSNP,^16^ gnomAD,^17^ 1000G,^2^ and GME.^18^ SV analysis revealed that each sample contained an average of 33,383 (30,811-37,314) structural variants (Fig. 3d) and 15,475 (14,157-17,444) unique SVs (Fig. 3e) in comparison with HPRC- CPC, CHM13 and GRCh38. Moreover, we identified 235,195 population-specific SVs that were absent in the HPRC-CPC pangenomes, CHM13, GRCh38 and other databases, including DGV^19^ and 1000G^2^ (Fig. 3f).

**Fig. 3:**
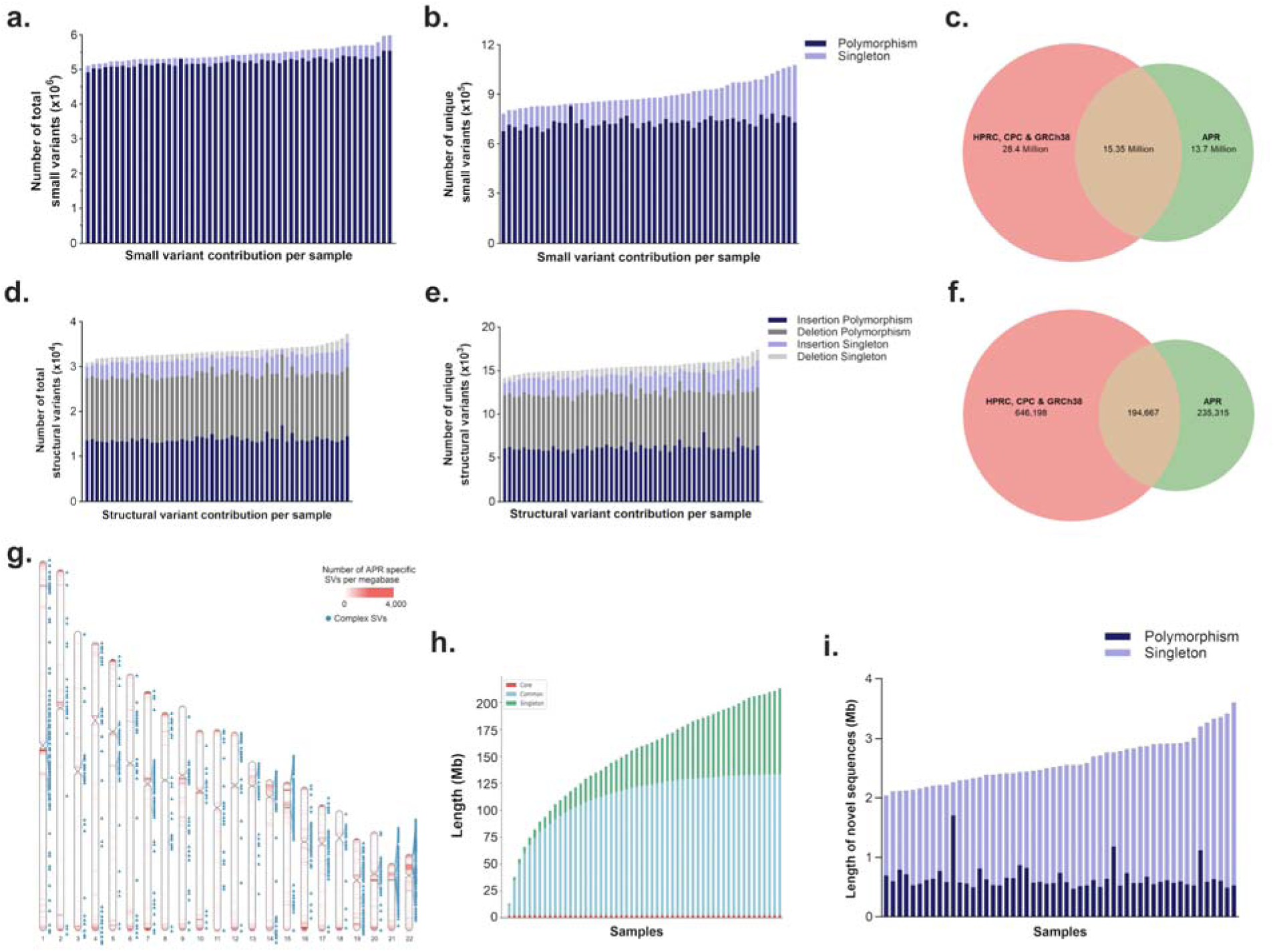
Arab genome specific sequences. **a,** Bar graph demonstrating the total number of small variants for each sample, distinguishing between singleton (light blue) and polymorphic (dark blue) variants. **b,** Bar graph showcasing the small variants specific to APR per sample, further differentiating between singleton and polymorphic variants. **c,** Venn diagram showcasing the small variants from the Arab pangenome in relation to HPRC, CPC, CHM13, GRCh38, dbSNP, gnomAD, 1000G, and GME datasets. **d,** Stacked bar graph detailing the total structural variants (SVs) per sample, categorizing between singleton and polymorphic variants for both insertions and deletions. **e,** Stacked bar graph illustrating the SVs that are APR-specific for each sample, for both insertions and deletions. **f,** Venn diagram visualizing the overlap and differences in SVs from the Arab pangenome with HPRC and CPC datasets, CHM13, GRCh38, 1000G and DGV. **g,** Visualization of Arab-specific SVs from the pangenome graph across autosomes. Sites of complex SVs are marked with blue. **h,** Pangenome growth curve for APR pangenome graph. Core represents (≥95%), common (≥5%), and singleton (only one haplotype). **i,** Bar graph displaying the length distribution of newly identified sequences for each sample, offering insights into the diversity of novel sequence lengths.

The average SV length was 3.08 kb (1.17kb–974kb), and the median was 154 bp (Supplementary Fig. 15). These findings validated the assembly integrity by showing the Alu and LINE-1 repeat content along the chromosomes. We further analyzed the Arab-specific SVs in the autosomes (Fig. 3g, Supplementary Table 22a), involving 8995 unique genes (Supplementary Table 22b).

These regions may harbor important genomic features that are specific to the Arab population. Genes affected by novel SVs in the APR were enriched in the synaptic component (p<5.42x10^-19^), regulation of cell projection organization (p<5.42x10^-19^) and small GTPase binding (p<6.35x10^-16^) pathways (Supplementary Table 23).

Pangenome growth was assessed using Panacus^38^ and we identified a total of 212.94 Mb of nonreference sequences that were added from the 53 diploid genomes. Of these, 35.30 Mb were classified as singletons, representing unique genetic variations specific to individual genomes. Furthermore, 2.60 Mb was present in ≥95% of all analyzed haplotypes, which can be considered the core genome of our sampled populations. Additionally, 133.57 Mb of nonreference sequences were identified in ≥5% of the haplotypes, representing a common genome among the Arab populations (Fig. 3h).

### Novel euchromatic sequences in the APR

We measured the number of novel sequences that are not present in GRCh38, T2T-CHM13, HPRC, CPC and DGV revealing that each Arab diploid genome harbored an average of 4,616 (4,049 - 5,787) novel sequences (Supplementary Fig. 16 a-b, Extended Data Fig. 4a). The pangenome graph added 111.96 Mb of unique non-reference sequences from the 106 diploid Arab genomes, including 104.04 Mb of singletons. The average non-reference sequence length per individual was 2.61 Mb (range 2.04-3.60 Mb) (Fig. 3i, Supplementary Table 24). A small percentage of the total novel sequences were present in centromeric and telomeric regions (1.62% and 5.53%, respectively) (Supplementary Fig. 16c). Nearly one-fifth (22.80%) of the novel sequences were in microsatellite regions (Supplementary Fig. 17a), especially in the pericentromeric satellite, HSat3 and other centromeric satellites. To characterize the context of these novel sequences located within 84,311 loci, we performed repeat annotation using RepeatMasker and found that LINE, SINE, LTR and satellite repeats constituted 8.22%, 8.38%, 4.99% and 26% of the novel sequences, respectively (Supplementary Fig. 17b; Supplementary Table 25). These novel sequences may represent novel functional elements or structural variations that are missed by conventional methods. These results demonstrate the power and utility of using long-read sequencing and pangenome graph construction to capture the genomic diversity and complexity of human populations.

### Complex structural variation in the APR pangenome graph

We used a pangenome graph to visualize and analyze complex structural variation (CSV) in the APR samples. A complex structural variation site was defined as an SV site with at least one 10 kb SV with a minimum of 5 haplotypes. We found that 1.40% (733 out of 52,465) of these SV sites were complex multiallelic bubbles (Supplementary Table 26a) and 406 sites were APR specific (Supplementary Table 26b). The CSVs were dispersed across all chromosomes, with a significant concentration on acrocentric chromosomes (average 38.3 CSVs), particularly chromosome 22, which had the highest count (60 CSVs). APR CSVs were predominantly located in pericentromeric satellite, HSat3 and alpha satellite DNA (Supplementary Fig. 18). Our analysis revealed that most CSVs were approximately 10 kb in length, which was the established lower bound for CSV inclusion in our analysis (Supplementary Fig. 19). Utilizing rigorous criteria, we delineated complex structural variation regions as areas within a 100 kb window that consisted of two or more multiallelic SV sites, with each site containing at least one 10 kb SV within the haplotypes. This approach led to the discovery of 659 CSV regions. The genes (Supplementary Table 27) nested within this 100 kb window of complex regions were enriched in the antigen processing and presentation (p<6.73x10^-08^) and immunoregulatory interaction (p<1.69x10^-05^) pathways (Supplementary Table 28). We further investigated the complex variant regions (Supplementary Table 29a) by overlapping them with APR-specific SVs (Supplementary Table 29b).

We compared the allelic variations in the APR to those in the combined HPRC+CPC pangenomes in regions of clinical relevance: the *PRAMEF* (Fig. 4a, Supplementary Table 30) and *POLR2J3* - *SPDYE2* (Fig. 4b) gene regions. In the *PRAMEF* region, we observed a common allele present in HPRC+CPC pangenomes that was also present in 86.80% of the APR samples, while others were unique haplotypes to the APR pangenome. We observed 2 novel haplotypes that were exclusively found in 13.20% of the Arab cohort. Preferentially expressed antigen in melanoma (PRAME) belongs to a group of cancer/testis antigens that are mainly expressed in the testis and an array of tumors and play crucial roles in immunity and reproduction.^39^ In the *POLR2J3* - *SPDYE2* region, we observed that seven alleles present in the HPRC+CPC pangenomes were absent in the APR pangenome. We observed three novel allele exclusively present in APR (absent in HPRC+CPC, CHM13 and GRCh38). The *POLR2J* gene encodes a subunit of RNA polymerase II, which is essential for synthesizing messenger RNA in eukaryotes.^40^ The *SPDYE2* gene encodes a member of the Speedy/RINGO cell cycle regulator family and is involved in protein kinase binding activity.^41^ Furthermore, allelic variations were observed in the HLA-DRB region that were absent in the references (Supplementary Fig. 20).

**Fig. 4:**
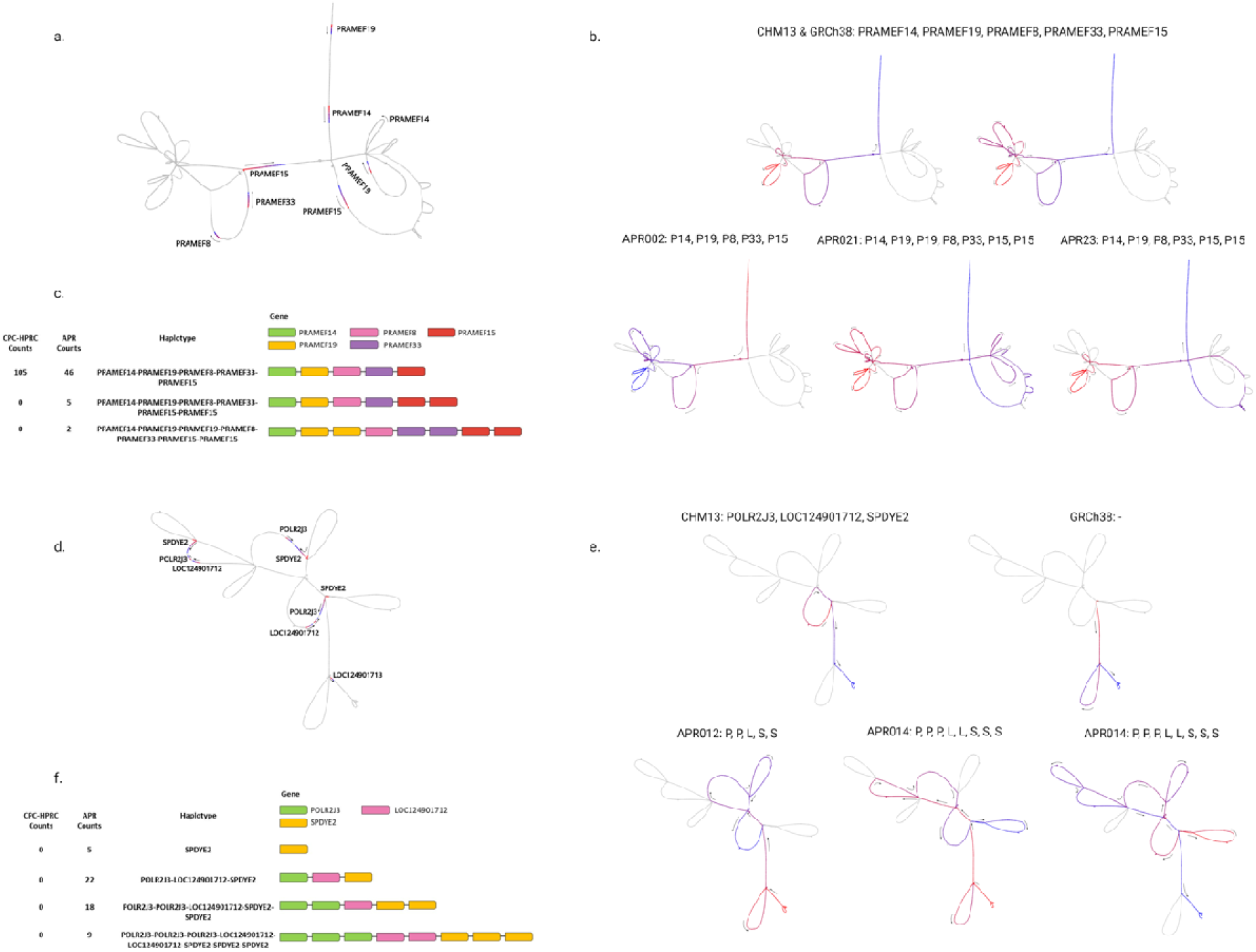
Visualizing complex structural variation region. **a,** PRAMEF region subgraph. Diagram showcasing the specific location of the PRAMEF genes. **b,** Sample haplotypes in PRAMEF Region. Distinct paths taken by different samples through the PRAMEF region. **c,** PRAMEF region haplotype count. Linear structural diagrams representing the frequency and structural visualization of haplotypes identified by the graph across 106 haploid assemblies, compared against the HPRC-CPC graph. **d,** *POLR2J3* - *SPDYE2* region subgraph. Diagram highlighting the specific location of the *POLR2J3* - *SPDYE2* region. **e,** Sample haplotypes in *POLR2J3* - *SPDYE2* region. Unique paths traversed by different samples through the *POLR2J3* - *SPDYE2* region. **f,** *POLR2J3* - *SPDYE2* region haplotype count. Linear structural diagrams depicting the frequency and structural visualization of haplotypes as determined by the graph among 106 haploid assemblies, compared with the HPRC-CPC graph for a comprehensive comparison. Variation among haplotype walks that did not involve genes was visualized using color coded lines, from red to blue to indicate directions.

Other CSV regions containing *CYP2D6* exhibited variable haplotype presences within the APR that were different from those of the CHM13 or GRCh38 references (Supplementary Fig. 21).

### Mitochondrial pangenome characterization

The mitochondrial Arab pangenome (mtAPR) was constructed using PacBio HiFi reads from 53 individuals (Supplementary Fig. 22a). The reads that mapped to ChrM with at least 90% similarity and a length greater than 15 kb were used. The average lengths of the resulting mtAPR and mtHPRC reads were 16.29 kb and 16.39 kb, respectively. The mtAPR graph encapsulating the mitochondrial heteroplasmy diversity consisted of 34,143 nodes and 61,187 edges (Fig. 5a). In contrast, the mtHPRC pangenome graph, derived from a comparable number of reads using CHM13 as a reference, consisted of 32,078 nodes and 57,494 edges. The graph shows a complex and diverse structure that reflects the genetic diversity of human mitochondrial DNA. We annotated the graph using the ChrM annotations from GENCODE (v38),^42^ facilitating the identification of mitochondrial genes.

**Fig. 5:**
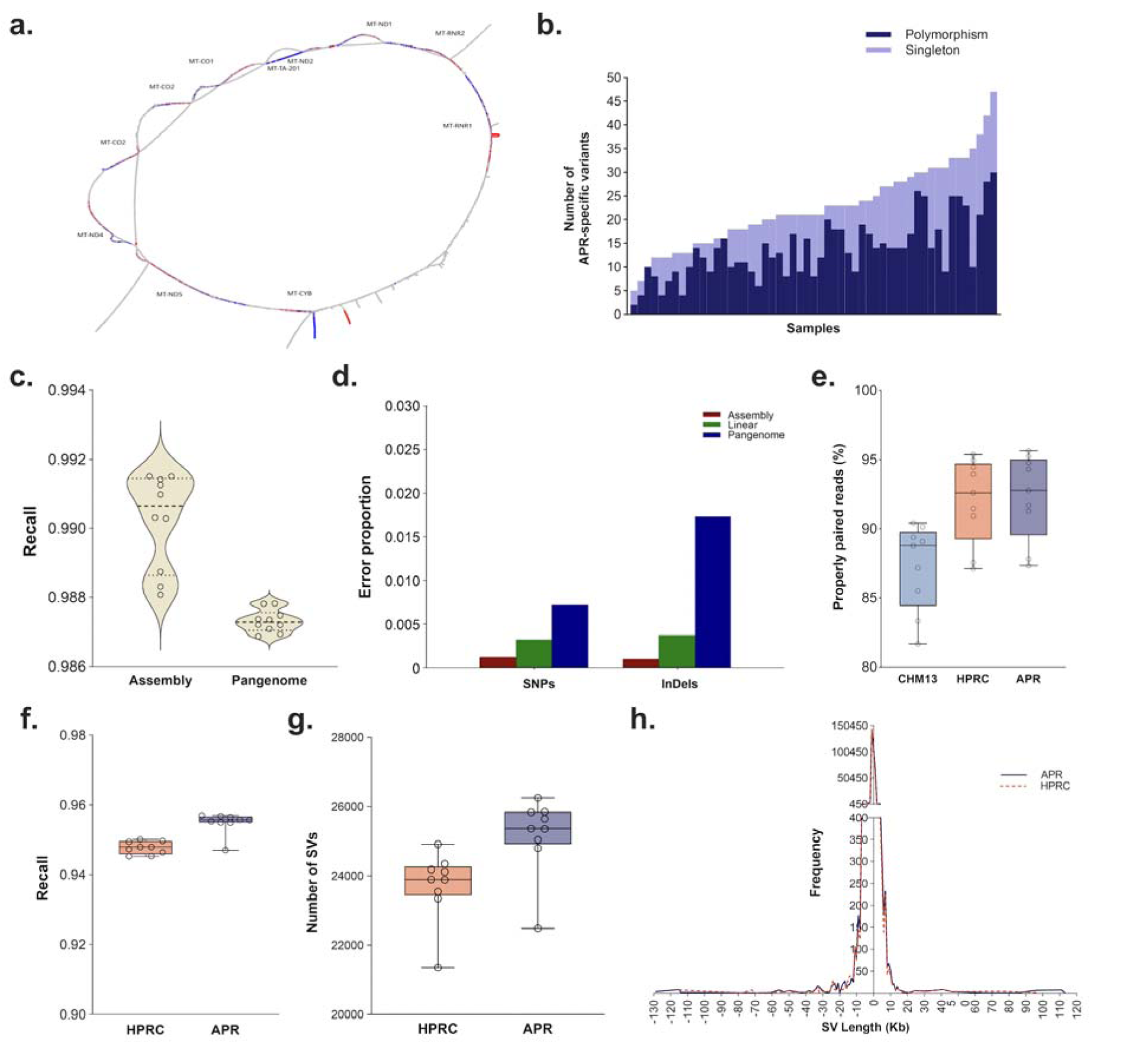
Mitochondrial pangenome analysis and nuclear pangenome performance gain. **a,** A circular representation of the mitochondrial pangenome, detailing the position and nomenclature of annotated mitochondrial genes within the pangenome. Each bubble or loop represents a haplotype. **b,** mtAPR variant distribution. A bar chart showcasing the number of APR-specific small variants observed across different samples in comparison to HPRC, differentiated between polymorphism (dark blue) and singleton (light blue). **c**, Comparative analysis of variant calling performance using linear, assembly and pangenome methods. Violin plot displaying the recall of linear variant calls using assembly-based and pangenome-based methods. **d,** Bar graph illustrating the proportion of errors in SNP and Indel variant calls using three different methods: assembly (red), linear (green), and pangenome (blue). **e,** Mapping accuracy assessment. Box plot illustrating the percentage of properly paired reads in alignments of 9 whole genome sequenced Arab samples (from UAE, Saudi, Syria, and Oman) to the APR and HPRC genomic graphs, compared to the CHM13 reference. **f,** Genotyping recall for SNPs. Box plot depicting the recall rates for genotyping of polymorphic variants in easy genomic region based on CHM13 variant calls. Easy genomic regions are defined as parts of the genome excluding segmental duplications, centromeric/satellite sequences, composite repeats, satellites, chrXY sequence classes, telomeres, and palindromes/inverted repeats. **g,** Structural variants across samples in easy genomic regions. Line graph comparing the count of structural variants identified across Arab samples mapped to the APR and HPRC graphs. **h,** Line graph depicting the frequency of SV lengths across Arab samples mapped to APR and HPRC graphs.

Comprehensive variant analysis within the mtAPR graph revealed 843 variant sites with a total of 3,291 variants, of which 902 were unique (Supplementary Fig. 22b, Supplementary Table 31a). Compared with the mtHPRC, the mtAPR revealed 623 unique small variants specific to the Arab pangenome, with 457 singletons and 166 polymorphic variants. These variants were distributed across 600 distinct sites (Fig. 5b, Supplementary Table 31b) cataloging the APR heteroplasmy heterogeneity. A novel structural variant spanning 652 bp encapsulating duplicated mitochondrial genes (*MT-TF* and *MT-RNR1*) was also detected. Insertions greater than 10 bp in length derived from the mitochondrial pangenome variant calling data cumulatively measured 1,738 bp (Supplementary Table 32a). Clustering of these sequences using the cd-hit-est program with a 90% similarity threshold (-c 0.9) revealed unique novel insertions that were not accounted for in the reference genomes, which included up to 1,436 bp of mitochondrial DNA (Supplementary Table 32b).

### Performance gains for pangenome-aided analysis

To determine the benefits of utilizing a pangenome-based approach in genomic analysis, we conducted a comprehensive evaluation comparing pangenome-based analyses with traditional methods. We assessed the performance of assembly based variant caller, PAV^43^ and graph-based variant calls by comparing them to a consensus set derived from HiFi read-based variant calls using DeepVariant and Longshot.^44,45^ The results demonstrated high overall variant recall rates, with an average of 0.990 (0.988-0.991) for PAV calls and 0.987 (0.987-0.988) for graph-based calls (Fig. 5c). This indicates that pangenome-based analyses can achieve comparable accuracy to read-based methods, emphasizing their utility in capturing diverse genetic variations efficiently. We further assessed the Mendelian concordance of the trio samples by comparing read-based, assembly-based and pangenome-based variant calls (Fig. 5d). The assembly-based calls had the lowest proportion of variants inconsistent with Mendelian inheritance, with error rates of 0.0011 for SNPs and 0.0009 for indels. Further, the APR pangenome variant calls exhibited lower errors in Mendelian concordance (with error rates of 0.0071 for SNPs and 0.0172 for indels), significantly lower than those observed by HPRC (with error rates of 0.0504 for SNPs and 0.0749 for indels).

We assessed the mapping accuracy of short reads from 9 whole genome sequenced Arab samples^9^ (UAE, Saudi, Syria and Oman). These samples were aligned to both APR and HPRC graphs using the Giraffe mapper^46^ and for comparison, reads were aligned to CHM13 using BWA-MEM. Our analysis showed that the average mapping rate for alignment to the pangenome graph increased to 92.32% (87.34-95.64) with APR and 92.04% (87.12-95.37) with HPRC, compared to 87.28% (81.68-90.42) when aligned to CHM13 (Fig. 5e). Additionally, we found that, on average, 5.04% fewer reads mapped to CHM13 compared to pangenome graph alignments, highlighting that these reads are better represented by the pangenome graph. Further analysis on the genotyping of polymorphic variants revealed high recall rates for easy genomic region (excluding satellite, repeat and segmental duplication region), with APR achieving a 95.49% (94.70-95.70) average genotyping recall and HPRC 94.78% (94.52-95.01), as determined against CHM13 variant calls (Fig. 5f). Recall rate for APR and HPRC for all region was 91.74% (91.21-92.04) and 90.71% (90.21-91.10) respectively (Supplementary Fig. 23a).

Our comparative analysis shows that the APR identified an average of 6.10% more structural variants in easy genomic regions compared to HPRC (Fig. 5g), with notable increases in both small and large structural variants (Fig. 5h). We conducted comprehensive genome sequencing on 4 Arab trios, each including a proband diagnosed with autism spectrum disorders (ASD), utilizing both PacBio HiFi long-read whole-genome sequencing (average coverage 21.85) and short-read whole-exome sequencing (average coverage 428) to evaluate the performance of the APR reference. The mapping accuracy of the exome sequencing data revealed an average mapping rate of 99.67% when aligned to the CHM13 reference genome and 99.62% to the APR graph, indicating a comparably high level of precision in genomic alignment for both references (Supplementary Fig. 23b). Our genotyping analysis revealed that when mapping whole-exome sequencing data to the Arab Pangenome Reference (APR), 93.29% and 90% of missense variants identified with the APR were also recalled using the short-read exome and long-read whole genome mapping to CHM13 reference, respectively (Supplementary Table 33). Overall, these findings demonstrate the APR’s robust performance in variant detection and recall, particularly for clinically relevant genetic alterations, underscoring its utility in genetic research and diagnostics.

## Discussion

We constructed an openly accessible pangenome reference specific to the Arab population from 53 deeply phenotyped apparently healthy adults by processing 106 high-quality assemblies.

These samples were subjected to careful phenotyping to identify manifestations of the onset and progression of chronic complex diseases. Using 35.27X PacBio HiFi reads and 54.22X ultralong reads, we achieved average contig N50 of 124.28 Mb, which is 3.11-fold longer than that recently reported by HPRC. *De novo* assembly algorithms used 99.96% of available long-read sequences to construct the contigs. Our average assembly quality score of QV 57.53 suggested a high quality of the assemblies used to construct the pangenome. The average haplotype of the APR samples showed 93.59% and 96.53% coverage of CHM13 and GRCh38, respectively, which is consistent with the HPRC and CPC assemblies. The coverage was lowest for the Y chromosome due to its complex Yq12 heterochromatin repeat complex region. The construction of a high-resolution reference pangenome for the Arab population provided novel insight into the unique genome organization within these populations. While PacBio HiFi data provided a base of highly accurate long reads, ONT ultralong reads contributed significantly to filling the gap in scaffolding and covering complex regions of APR genomes. The ULK data (>100 kb) exhibited an average coverage of 12.53X for the APR samples, contributing significantly to the extended N50 length that APR achieved.

The CHM13 reference is the most complete human genome to date, and GRCh38 has the most complete annotations of functional elements. The Arab pangenome was constructed based on CHM13 by integrating GRCh38 and the APR assemblies. We observed a significant overlap between APR and HPRC+CPC variants. We anticipate that the application of the APR in clinical settings will markedly increase the diagnostic yield for numerous single-gene disorders. The identified population-specific variants illustrate the separation time of Middle Eastern populations from other continental groups and the subsequent effects of isolation, admixture and drift^9^. Unfortunately, the lack of representative data from the Middle Eastern populations in large genomic initiatives (i.e., the 1000 Genomes Project, gnomAD) prevented the generation of impactful genomic resources for this region. To assess the clinical significance of thousands of novel APR-specific variants, further follow up clinical cohort-based analyses are needed. One of the unresolved limitations that we observed across all pangenomes, is the mapping rate within the complex satellite regions which still suffers in accuracy. Significant care should be taken to study these regions where mutations rate is known to be multiple-fold higher.^47^

APR-specific structural variants revealed 111.96 Mb sequences (range, 2.04 to 3.60 Mb of new sequences per haplotype) that were not present within the CHM13 and GRCh38 reference genomes or the HPRC and CPC pangenomes and DGV. These novel sequences include highly repeated satellites that are inaccessible by short-read technologies. A linear reference genome is limited by its static sequences, which hinder the ability to decipher the polymorphic nature of SVs. The clinical application of APR pangenome-based SV detection will increase the clinical yield due to the large number of uncharacterized SVs. Moreover, the APR will enable the detection of base resolution phased haplotype complexities to infer their association with diseases. The SVs have a profound effect on how the APR differs from the HPRC or CPC graphs. The APR included 235,195 unique SVs (with an average of 15,472 unique SVs per sample) that create different bubbles or haplotype walks (involving genes) within these genomic regions compared to other pangenomes or human genome references. Our findings suggest that using a larger sample size could produce a more accurate and comprehensive pangenome, capturing additional rare gene duplications and novel sequences. These rare genetic variants have the potential to enhance the diagnostic yield for rare genetic diseases and cancer. By employing a pangenome graph approach, we were able to identify novel complex haplotypes that might contribute to various disorders and to track the origin and distribution of these haplotypes among different cohorts. In the analysis of short-read whole-genome data from Arab individuals, we observed improved accuracy in mapping paired reads using APR compared to CHM13. Additionally, the transcriptome annotation is highly complete, particularly for protein-coding transcripts, for which the mapping rate was approximately 99%. However, short- or long-read sequence alignment, variant calling and phasing workflows using the pangenome remain too time-consuming for clinical laboratories. Through our analysis, we observed mapping and variant genotyping rate from short- or long-read improve when using APR, its practical application is expected to significantly improve over time.

The mapping and genotyping accuracy demonstrated a significant advantage for APR over linear reference CHM13. This is largely attributed to the unique APR-specific sequences and the utilization of a graph genome that integrates multiple individuals as a reference. Novel methodologies are needed for rapid and precise graph pangenome mapping of long-read whole genome to fully leverage the benefits of global graph pangenomes. Our short-read mapping to APR and recalling rate on long read data shows over 90% recall rate of missense variants. The additional variants detected uniquely in the APR mapping but not in CHM13 could likely be ascribed to differences stemming from variant detection sensitivity within the complex repetitive regions, APR assemblies with novel sequences, differences in the algorithms used for read mapping, variant calling and assembly building, difference in short and long read sequencing technologies and potential assembly inaccuracies. Overall, our findings underscore the APR’s utility in capturing a broader spectrum of genomic variants, with the potential to enhance genetic diagnosis yield and additional genomic insights into ASD and other rare disorders within Arab populations.

Our APR-specific gene duplication analysis identified 883 unique genes involved in ribonucleotide metabolism and oxidative phosphorylation pathways. Of the duplicated genes identified, 15.06% have been previously associated with recessive conditions, which is particularly prominent in Arab populations.^48^ Duplicated recessive genes may increase the risk factor of manifesting rare diseases, we have not found any recessive pathogenic variants within these genes in our APR cohort. According to previous reports,^49^ gene duplications are highly active in regions with segmental duplications. *TAF11L5* is the most frequently duplicated gene in APR cohort, it encodes TATA-box binding protein (TBP), which is predicted to be involved in RNA polymerase II preinitiation complex assembly. Our results suggest that TBP is a functionally active element within Arab populations and follow up studies will be required to assess the role of *TAF11L5* gene duplication. The core conserved region of TBP that is also duplicated within the APR, is predominantly involved in double-stranded nucleic acid binding and enables transcription initiation.^50^

The mitochondrial APR contributed novel 1,436 bp sequence that was absent from the HPRC mitochondrial pangenome. This addition significantly enriches the Arab mitochondrial reference with a diverse array of haplotypes and variants. Total mtAPR unique variants impact 600 sites of the CHM13 mtDNA bases, effectively cataloging the mitochondrial heteroplasmy haplotypes found in healthy Arab populations. As the APR cohort was clinically assessed as healthy, this Arab mitochondrial pangenome serves not only as a comprehensive reference but also as a crucial tool for mitochondrial disease diagnostics, offering valuable heteroplasmy frequency data specific to the Arab population.

The understanding of human genomic diversity and complexity depends on the collective effort to produce multiple pangenomes from different ethnicities. Currently, technological limitations hinder our ability to precisely decipher heterochromatic regions, specifically those near the centromeres. We have observed, despite using various alignment algorithms, mapping reads into heterochromatic region still is a challenging problem. These regions harbor highly complex repeats and mobile elements. However, advancements in sequencing technologies and algorithms are expected to gradually overcome the difficulties in decoding regions with complex repeats.

While we acknowledge that more samples are needed to adequately represent the genetic diversity within Arab populations, the APR represents a crucial foundation for genetic research. Arab populations are notably underrepresented in large genomic databases, and the APR will address this gap by offering essential resources for clinical genomic laboratories to enhance the precision of variant interpretation.

## Methods

### Sample collection and phenotyping

We collected 8-10 ml blood samples from 53 healthy individuals (one trio and 50 unrelated individuals) of Arab descent from the United Arab Emirates (UAE), Saudi Arabia, Oman, Jordan, Egypt, Morocco, Syria and Yemen. Informed written consent was obtained from each participant regarding their participation in our research study and sharing the data for public access. In addition, we included a family trio (mother (APR-M), father (APR-F), and child (APR-S)) for in-depth analysis. The inclusion criteria were self-identified Arabs who were 18 years or older, willing to participate voluntarily and presumptively healthy (free from diseases such as diabetes, cardiovascular disease, hypertension, chronic kidney disease, lung disease and liver disease). To map long-read genome and exome sequencing data to the APR, we recruited 4 trios including children with autism spectrum disorders (ASDs) to conduct whole exome short read and whole genome long read sequencing. Furthermore, we processed short-read Illumina whole-genome sequencing data from 9 published Arab samples.^9^ This study was approved by the Institutional Review Boards of Dubai Scientific Research Ethics Committee (DSREC- 12/2022_01, DAHC/MBRU-IRB/2023-15) and Mohammed Bin Rashid University of Medicine and Health Sciences (MBRU-IRB-2017-004).

### DNA isolation and sequencing

#### Pacific Bioscience High-Fidelity sequencing

For PacBio HiFi long-read sequencing of APR cohort samples and ASD trios, high-molecular-weight DNA was isolated from 200 *µL* of flash-frozen blood using both GenFind V3 and Nanobind DNA extraction kits per the manufacturer’s instructions. The DNA concentration was measured using a Qubit 3 fluorometer with a dsDNA HS Assay kit (Thermo Fisher), and the samples were adjusted to a concentration of 30 ng/*µL* in a total volume of 130 µL for shearing. Shearing was performed on a Diagenode Megaruptor 3 hydropore-Syringe at a speed setting of 29-31, targeting 15-18 kb fragment lengths according to PacBio’s recommendations. The sheared sample fragment lengths were verified on a 4200 TapeStation system using genomic DNA screen tape. SMRTbell libraries were prepared using the SMRTbell Prep Kit 3.0, with libraries anchored to Sequel II primer 3.2 and Sequel II DNA polymerase 2.2. Sequencing was conducted on PacBio Sequel IIe equipment using SMRT cells 8M and the Sequel II Sequencing Kit 2.0.

The sequencing protocol was designed to enable adaptive loading, with 2 hours of preextension and 30 hours of movie capture. For each sample run on Sequel 11e, three SMRT cells were used. A total of 17 samples were processed on a Revio system using a Revio SMRT cell tray and a polymerase kit (Supplementary Methods Section 2).

#### Oxford Nanopore Technologies (ONT) ultralong DNA sequencing

Blood samples from the APR cohort were aliquoted into 1.8 ml cryovials, flash frozen and stored at -80 °C until further processing. For sequencing, frozen aliquots were thawed, and peripheral blood mononuclear cells (PBMCs) were isolated using red blood cell lysis buffer. The purified PBMCs were counted, and approximately 60 million cells per sample were subjected to ultralong DNA extraction using the NEB Monarch Tissue DNA Extraction Kit (#T3060L). Sequencing libraries were then prepared with the extracted DNA using the Oxford Nanopore Technologies Ultra-Long DNA Sequencing Kit (SQK-ULK114) following the manufacturer’s protocol with minor modifications. Briefly, the DNA was tagmented at room temperature for 10 min, followed by a 10 min incubation at 75 °C. Rapid sequencing adapters were added, and the samples were incubated for 30 min at room temperature. Cleanup of the adapted DNA was performed using the precipitation star and buffers provided in the kit. The final libraries were quantified and loaded onto a minimum of 3 PromethION flow cells at approximately 30 ng per flow cell (Supplementary Table 2). Sequencing was performed for 96 hours per flow cell using R10.4.1 kits, with a minimum read length setting of 1 kb.

#### Hi-C sequencing

The Hi-C libraries were prepared using the Qiagen EpiTect Hi-C Kit with significant modifications. PBMCs were isolated from approximately 1 ml of frozen blood, with cell counts ranging from 5x10^3^ to 2.5x10^6^ cells per sample. The cells were first pelleted using a 10× volume of RBC lysis buffer, followed by washing in PBS containing 2% FBS. The cell pellet was then fixed in 1% formaldehyde at room temperature, and crosslinking was quenched using 3 M Tris, followed by washing with PBS. The fixed cells were lysed using Hi-C lysis buffer, and the crosslinked chromatin was digested at GATC sites using a specific combination of endonuclease and buffer. This step was followed by the standard Hi-C protocol for end labeling, ligation, chromatin decrosslinking, and purification. The purified DNA was fragmented to 400-600 bp fragments using a Bioruptor Pico sonicator, followed by purification and enrichment. The DNA was then prepared for sequencing with dual indexing using Illumina barcode adapters and amplified using Illumina sequencing primers. The final step involved the purification of amplified libraries, which were quantified, pooled, and sequenced on an Illumina NovaSeq-6000 platform, achieving read lengths of 2 x 150 bp.

### Sequencing and variant QC

For ONT sequencing data, base calling was performed using Oxford Nanopore’s high-accuracy model utilizing Guppy basecaller v6.5.7 with modifications for detecting 5-methylcytosine (Supplementary Methods Section 3). We employed the high accuracy (HAC) model to maximize the precision of the derived base sequences. The PacBio sequencing data were processed using the circular consensus sequencing (CCS) algorithm. In the subsequent quality control steps, we employed NanoStat v1.6.0, which provides an in-depth statistical overview of the sequence data. Additionally, metrics obtained from the smrtlink software were incorporated into the analysis to provide a comprehensive assessment. To refine the PacBio reads, HiFiAdapterFilt v2.0.1 was used to remove any residual adapter sequences that might hinder the assembly process.

The PacBio human whole-genome sequencing (WGS) pipeline was utilized for read alignment and variant calling. Reads were aligned to the GRCh38 and CHM13 reference genomes using pbmm2 v1.10.0. The CHM13 reference used was chm13v2.0_maskedY_rCRS.fa, while the hg38 reference used was human_GRCh38_no_alt_analysis_set.fasta. Variant calling was then performed using DeepVariant v1.5.0. pbsv version 2.9 was utilized to jointly call SVs from the aligned long reads. To identify small variants, DeepVariant and GLnexus (v1.4.1) were applied for joint SNP and indel calling. Variant annotation was performed using sliVAR (v0.2.2) for small variants and svPACK for SVs, which also includes functional predictions and other annotations for the raw variant calls. All pass variants with GQ>= 10 were used for novel variant calculations. We used bedtools (version 2.31.0) to compute the coverage of individual chromosomes and pericentromeric regions across all samples. We utilized Longshot^45^ for detecting SNPs from sequencing reads using the following command:

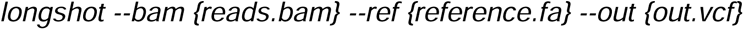

### Population ancestry inference

To discern the genetic diversity and ascertain the population structure of the 53 APR genomes, we created a merged dataset of published global and Middle Eastern populations with our samples. We performed variant calling using the command:

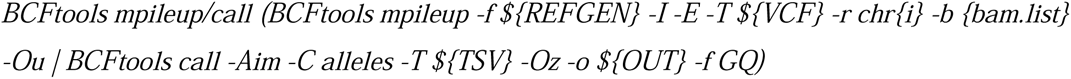

on a set of 596,417 variants found in the Human Origins array dataset downloaded from the Allen Ancient DNA Resource (V54.1.p1).^51^ The joint called APR VCF was merged with a set of 1040 samples from the Human Origins (HO) dataset. This dataset comprised 54 global populations from the HGDP dataset (which contains 4 Middle Eastern populations: Bedouins, Druze, Palestinians and Mozabites), in addition to other relevant Middle Eastern populations (Egyptians, Moroccans, Saudis, Yemenis, Iraqis, Syrians, and Jordanians) genotyped on the HO array.^20–22^ We excluded any sample that had an ‘outlier’ label. We subsequently added additional relevant Middle Eastern samples from Emiratis, Saudis, Omanis, Yemenis, Iraqis, Jordanians, and Syrians.^9^ The merged file was filtered for genotype quality (GQ) greater than 20, minor allele frequency (MAF) greater than 0.05, and linkage disequilibrium pruning using the following command:

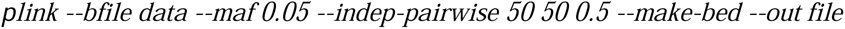

LD filtering revealed 182,322 variants. We then employed SNPRelate (v1.28.0)^52^ on a merged dataset with APR (HiFi variants), and Human Origins (1040 samples, including 304 Arab samples). A total of 182,322 SNPs per sample were utilized for principal component analysis (PCA) and ADMIXTURE. We applied the model-based clustering ADMIXTURE tool^53^ to model the genetic diversity of all 53 APR samples in the merged global dataset. We used the major continental labels from the HGDP dataset (South Asian, European, East Asian, American, African, and Oceanian), to which we added ‘Arab’ to our APR samples. We used *k* values ranging from 3 to 8 and found that k=8 had the lowest cross-validation score.

For the relatedness analysis, we ran the following command on the pruned dataset:

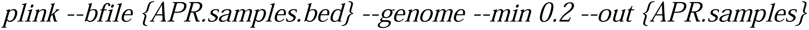

This command performs a genome-wide relatedness check, printing out pairs of samples that share more than 20% identity-by-descent, effectively flagging any related individuals. Further, to confirm the unrelated status of all 50 APR samples, we constructed a heatmap using haplotype sharing variance obtained from fineSTRUCTURE^54^ runs. In brief, variants called using DeepVariant were filtered for a GQ of ≥20. We then reduced the dataset to one variant per 10 kb window, jointly phased it with Eagle, and processed the data using the fineSTRUCTURE pipeline.

#### Mitochondrial and Y haplogroup classification

We calculated the mitochondrial haplogroups of the APR samples using the Haplogrep^55^ tool (v2.4.0). To contextualize our findings, we extracted haplogroup frequency information for various ethnicities, including Arab regions, from published literature.^56,57^ The nonrecombining region of the Y chromosome (NRY) haplogroups in the APR samples was classified using the Y-LineageTracker^58^ tool (version 1.3.0). We conducted an extensive literature review to compile Y-haplogroup frequency data across ethnicities.^59–61^ Both mitochondrial and Y chromosome haplogroup distributions were visualized using hierarchical clustered heatmaps generated with the Python Seaborn library to illustrate the comparative haplotype frequencies among different populations.

### *De novo* genome assembly

Sequence reads were screened for contaminant sequences and polished to remove artifacts prior to analysis. Kraken^26^ was used to taxonomically classify the assembled contigs, retaining only those annotated as human. Kraken was run using the following command:

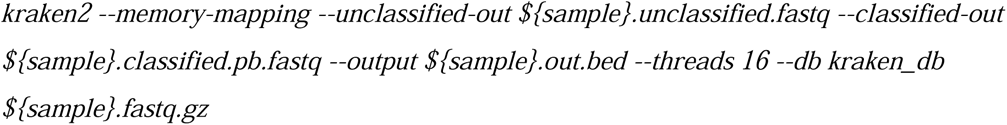

High-quality *de novo* assemblies were generated for each sample using a hybrid assembly approach, combining PacBio HiFi, ONT ultralong reads and Hi-C reads. A total of 53 samples were sequenced on 1-6 SMRT cells and 3-7 ONT flow cells (Supplementary Table 4). We combined PacBio HiFi reads, ultralong (ULK) reads and Hi-C reads from multiple runs of each sample and used Hifiasm^24^ (v0.19.5-r603) to carry out both primary and diploid *de novo* assembly for 53 samples (Supplementary Methods Section 4). The following commands were used to generate the assemblies:

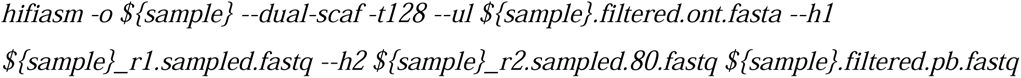

To evaluate and compare the effectiveness of genome assembly approaches, we utilized both Verkko (v1.3.1) and Hifiasm for constructing genome assemblies of both trio and individual samples within our dataset. Our initial step involved the generation of Merqury hapmer databases, which are crucial for facilitating high-quality, k-mer-based evaluation of haplotype assemblies. We generated hapmer databases for our trio samples using Merqury with the following command:

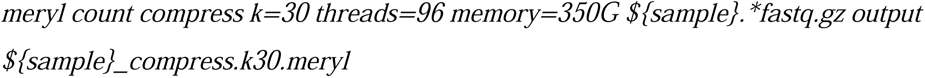

Following the creation of the hapmer databases, we proceeded with the assembly process using Verkko, which was configured specifically to leverage the advantages of trio-based phased assembly. Verkko was run with the following parameters:

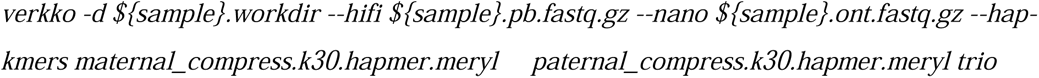

#### Genome assembly polishing

We employed Inspector (v1.2) to evaluate and improve the quality of the 106 genome assemblies. Initially, we assessed assembly errors using Inspector, followed by an error correction step executed using the following command:

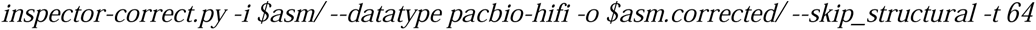

This step was specifically designed to refine the assemblies by addressing identified inaccuracies while excluding structural errors.

Mitochondrial read pairs were identified by mapping to ChrM using minimap2 (v2.26) with the option *-L -eqx,* and mapped reads were filtered using samtools (v1.6) view -F 4, and any corresponding mitochondrial contigs were removed. Potential interchromosomal joins in the assembly were identified using minimap software, followed by paftools.^27^ The specific commands initiated were *-cxasm chm13v2.0.fa APR043.bp.hap1.p_ctg.fa* and *paftools misjoin-e APR043.1.misjoins.paf,* respectively (Supplementary Methods Section 6).

We employed PAV^43^ for identifying variants from genome assemblies. The assemblies were sourced from ‘assemblies.tsv’, and were analyzed against a reference genome file ‘config.json’. The command executed for this analysis was:

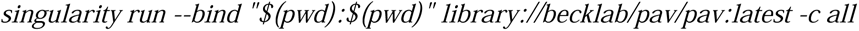

#### Assembly quality assessment

To evaluate the quality and structural integrity of 53 primary assemblies and 106 haplotype assemblies, we used QUAST (v5.2.0),^62^ which provides metrics such as completeness, N50, and number of contigs. QUAST was run with the following extensive parameter set, as detailed:

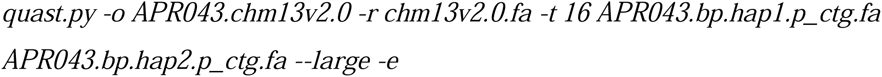

The yak suite (v0.1-r66),^24^ which includes yak count, yak qv and yak trioeval, was employed for genome quality validation.

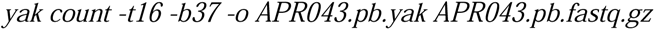

was first used to build kmer database and yak qv was used to calculate the QV score with the following parameters:

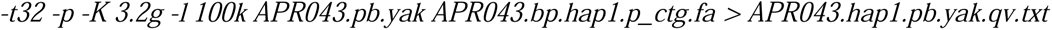

The mapping rate was assessed using minimap2 (v2.26).

We utilized Flagger (v0.3.3) to identify small-scale errors within the diploid assemblies. The HiFi reads were mapped to the concatenated assembly of both haplotypes using the following minimap2 command:

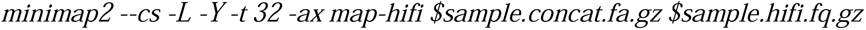

The mapped reads were then sorted using SAMtools sort. Each haplotype assembly was further mapped to chm13v2.0 using the following command:

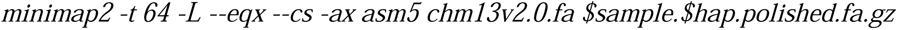

The Flagger WDL pipeline^29^ was subsequently executed using the sorted bam files as input in no-variant-calling mode. The resulting bed files were analyzed to differentiate between the haplotypes within the assemblies.

All assembled contigs were aligned to the CHM13v2 reference genome using Minimap2 with *-L --eqx* option. After successful mapping, the chromosome to which each contig was predominantly aligned was identified for further analysis. The centromeric region corresponding to this chromosome was then extracted from the reference genome using *SAMtools faidx*. To assess the alignment quality of the assembly contig to the extracted centromeric region, we employed UniAligner,^31^ specifically using the tandem_aligner command:

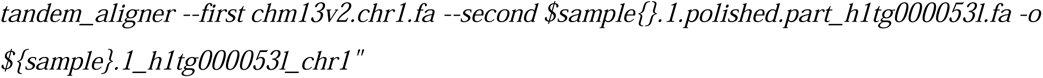

The output of this alignment is a CIGAR string and each contig’s alignment to the centromere of its respective chromosome was further analyzed. A sliding window approach was applied to each CIGAR string, using a window size of 100 base pairs, to calculate the alignment percentage within each window. Finally, the alignment percentages from all windows were plotted to visualize the distribution of alignment quality across different sections of the contigs.

To calculate the Hamming rate and switch errors in assemblies constructed with Hi-C data, we used pstools^30^ (v0.1). The phasing errors were assessed employing the following command:

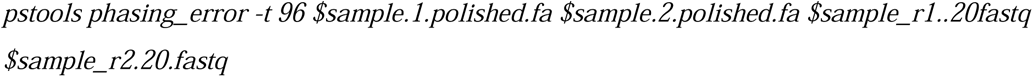

#### Gene duplication

We used Liftoff^32^ v1.6.3Click or tap here to enter text., a tool that accurately maps gene annotations between genome assemblies, to identify gene duplications in our dataset. Liftoff aligns gene sequences from a reference (GENCODE v38) to a target genome and finds the mapping that maximizes sequence identity while preserving the gene structure. An identity threshold of 90% (-sc 0.9) was used, and the following command was used:

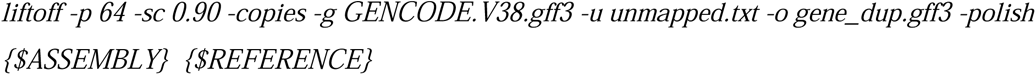

To remove partial matches from the analysis, we used the -exclude_partial option in Liftoff. We then quantitatively assessed the frequency of copy number variations (CNVs) for each gene in the target genome by comparing the number of gene copies to that in the reference (GENCODE GRCh38.p14 (GENCODE v38)). APR-specific gene duplications were compared against HPRC and CPC gene duplication matrices, as reported in studies utilizing similar methodologies and tool versions. We have constructed a downstream analytical tool PanScan (https://github.com/muddinmbru/panscan) for analyses of complex structural variant, gene duplication analysis and novel sequence estimation from pangenome graph vcf file.

#### Pangenome graph construction

We constructed Arab pangenome variation graphs using the Minigraph-Cactus pipeline (v2.7.2)^37^, which combines structural variants and single-nucleotide variants (SNVs) into a single graph representation. We used two reference genomes, GRCh38 and CHM13, with CHM13 as the backbone for graph construction following the steps described in the Minigraph-Cactus documentation. Initially, an SV graph (>50 bp) was constructed using Minigraph^63^ by sequentially aligning the 106 assemblies to the reference genome. Centromeric and telomeric regions were masked using dna-brnn,^64^ and the assemblies were remapped to exclude highly repetitive sequences. Contigs were then split and assigned to chromosomes based on their alignment coordinates. Base-level alignment was performed using Cactus, and the HAL output^65^ was converted to vg format (hal2vg). Paths >10 kb that were not aligned to the graph were removed, and the graph was normalized with GFAffix. The chromosome graphs were combined into a whole-genome graph, indexed with vg v1.50.1, and exported to VCF format (vg view).

Subsequently, SNP variants were incorporated into the SV graph using the same pipeline. The assemblies were remapped to the SV graph, and quality control was applied, excluding softmasked sequences >100 kb and alignments with MAPQ < 5. Cactus was executed on the chromosome graphs to introduce base-level variants, and the outputs were converted to vg format. The same filtering, normalization, combining, and indexing steps were applied to produce the final SNP+SV Arab pangenome graph. All analyses were conducted using the CATG high-performance computing cluster Flamingo. The command line used in our end-to-end pangenome construction pipeline is as follows:

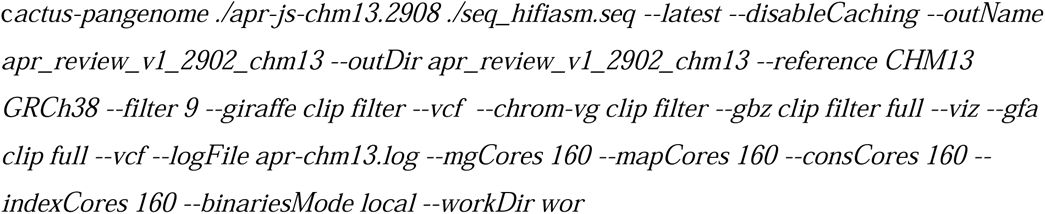

### Pangenome growth curve

Panacus^38^ was used to calculate the APR pangenome growth. The percentages of samples included were >5% (common), >95% (core) of the total samples and 1 haplotype (singleton). The command used was:

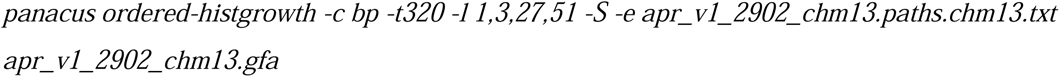

### Identification and visualization of population-specific variants and novel sequences

The multiallelic sites in the VCF file were split into biallelic records using the "bcftools norm - m-" command, and then the VCF file was separated into SNPs and other complex variants using custom Perl script. The complex variants were decomposed into SNPs, indels (<=50 bp) and SVs (>50 bp) using the command:

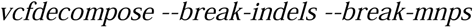

from RTG tools (v.3.12.1), and the genotype information from identical variants across multiple samples was merged to create a single record using custom Perl script. The SNPs obtained from "vcfdecompose" were also included in the downstream analysis. The same methodology was used for the analysis of the CPC-HPRC VCF file.

Unique SVs (<80% reciprocal overlap with HPRC and CPC, 1000 Genomes, and DGV) found in the Arab assemblies were classified as population specific. The unique SVs in the APR were identified by comparison with the HPRC-CPC SVs using the Truvari command:

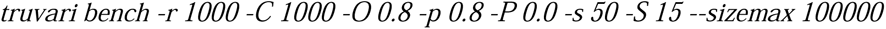

A reciprocal overlap of 80% was the criterion used to classify an SV as unique. This process collapses multiple SVs into one unique SV within our cohort. The resulting SVs were compared against the DGV Gold Standard Variants and the 1000 Genomes Project data (phase1_v3.20101123) with 80% reciprocal overlap using custom perl script. To identify novel sequences within the APR unique SV insertions, we clustered them using cd-hit-est (4.8.1) with the default sequence identity threshold of 0.9.

We visualized SV distributions and enrichment significance on chromosomes with RIdeogram^66^ (v0.2.2). Subgraphs surrounding SVs were extracted with gfabase v0.6.0 and visualized in Bandage NG version^67^ (v2022.09) after aligning gene sequences with GraphAligner^68^ (v1.0.17) using the following command:

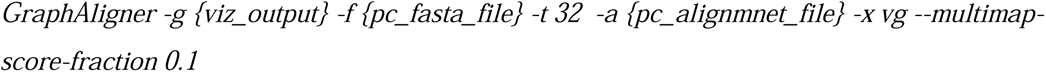

The gfatools package v0.5-r287 was subsequently used to derive in-depth statistical data from the pangenome graphs.

To identify the repeat elements within novel sequences, we screened fasta files with RepeatMasker (4.1.2-p1) using command:

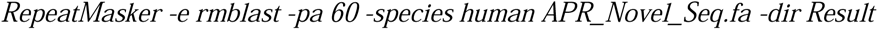

### Assessment of Mendelian concordance in trio samples

To evaluate the consistency of our trio samples’ variant calls with Mendelian concordance, we employed the PLINK command:

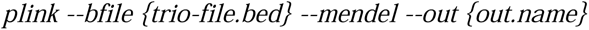

### Complex SV region and Bandage plotting

A site was considered complex if at least one variant larger than 10 kb was present, as well as at least five different alleles. The complex sites were identified by analyzing the snarls VCF files. The unique SVs were then overlapped with the complex sites to identify unique complex sites (see unique variants subsection) in the Arab population. The precise locations of the genes were determined by mapping the gene sequences to the graph using a graph aligner with the following parameter: --multimap-score 0.1. If multiple genes mapped to the same location (in the case of isoforms), only the gene mapping with the highest accuracy was retained. Complex structural variation (CSV) regions were defined as areas within a 100-kilobase (kb) window that consisted of two or more multiallelic complex SV sites, with each site containing at least one 10-kilobase (kb) SV within the haplotypes. The complex sites were plotted using Bandage. The 50 Kbp flanking region for each complex site was extracted using the Gfabase sub. To determine whether a haplotype consisted of a gene, the path was followed to determine whether it traversed through the gene region. Variation among haplotype walks that did not involve genes was visualized using color coded lines, from red to blue to indicate directions.

### Mitochondrial pangenome construction

To construct a mitochondrial Arab pangenome (mtAPR) that captures the diversity of Arab mitochondrial DNA, we used high-quality long reads from 53 individuals. We first mapped the reads to ChrM of the CHM13v2 reference genome using minimap2^69^ (v2.26) with 90% similarity and retained only reads that were longer than 15 kb; this threshold was set to substantially reduce the chances of inadvertent nuclear DNA contamination. This resulted in a total of 20,520 reads (19,251 ONT reads and 1,385 HiFi reads). We used PGGB^70^ (v0.5.4) to construct a mitochondrial pangenome graph from the mitochondrial contigs of HiFi reads of all individuals, and each read of >15 kb was concatenated in one fasta file along with the CHM13v2 mitochondrial chromosome. PGGB was then run on these samples with the following command:

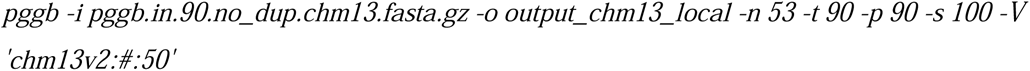

We visualized the mtAPR graph using Bandage NG version^67^ (v2022.09) and displayed the nodes, edges and variants in different colors and shapes.

### Mitochondrial pangenome variant identification

To process the VCF files of the mtAPR pangenome, we employed a multistage approach to ensure data accuracy and integrity. The initial step involved segregation of multiallelic variant sites into biallelic records, which was accomplished using the BCFtools normalization function (BCFtools norm -m). Next, we implemented the RTG tool vcfdecompose (version 3.12.1) with the parameters --break-mnps and --break-indels to further resolve complex variants into their constituent single-nucleotide polymorphisms (SNPs), indels and SVs. To merge genotype information, identical variants identified across the dataset were consolidated into singular records. This step was carried out using a custom Perl script. Parallel to the APR data, the HPRC mitochondrial pangenome VCF file was subjected to the same rigorous processing methodology to maintain consistency across datasets. The small variants (<10 bp) were further filtered to obtain those that were concordant with DeepVariant calls of the mitochondrial reads. To identify novel variants unique to the APR mitochondrial pangenome, we systematically removed variants that were shared with the HPRC mitochondrial pangenome. We then extracted insertions of 10 base pairs or more that were subsequently clustered using the cd-hit-est program, which applied a stringent 90% similarity threshold (-c 0.9) to discern and characterize novel insertions.

### Evaluation of short read whole genome and whole exome mapping

We used 9 WGS data from Arab descent^9^ to quantify the mappability accuracy of short read sequences into APR and HPRC pangenome and CHM13 reference. The quality metrics of each samples were obtained using FASTQC tool (https://www.bioinformatics.babraham.ac.uk/projects/fastqc/) and the low quality bases were removed using fastp (version 0.23.4).

To calculate the mapping rate of short reads to CHM13, we first aligned the paired end reads to CHM13v2 using bwa-mem. The following commands were used:

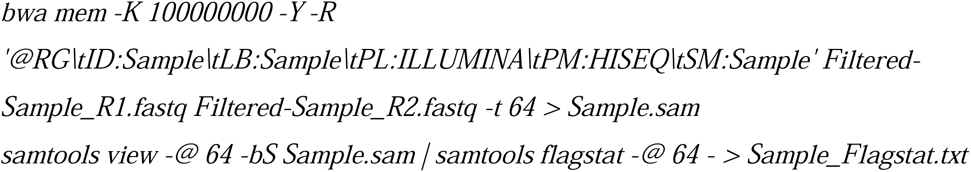

The variant calling was performed using DeepVariant (v1.5.0) by providing --model_type=WGS for the whole genome samples and --model_type=WES for the whole exome samples respectively.

To determine the mapping rate of short reads to the APR and HPRC graphs and to conduct variant calling, filtered samples were mapped to pangenomes using vg Giraffe (v1.55.0). The "vg stats" command was used to produce stats for the properly paired reads for graph references and "vg call" command was used to call the variants:

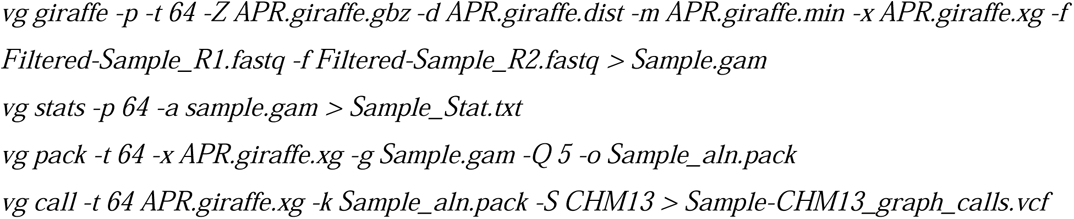

### Pathway enrichment analysis

To discern the biological pathways affected by gene sets, we conducted gene set enrichment analysis (GSEA) leveraging the KEGG and GO pathway databases, a well-curated repository featuring pathways that include molecular interactions, reactions, and relation networks. Gene names were converted to Entrez gene identifiers using the bioMart tool. Our analytical framework included determining the degree of overlap between the gene sets and the pathways in the KEGG-GO database, utilizing GeneOverlap to conduct one-sided Fisher’s exact tests (FETs). We excluded pathway gene sets that numbered fewer than 50 or exceeded 1000. To account for multiple hypothesis testing and control the false discovery rate (FDR), we applied the Benjamini Hochberg procedure to adjust the *p* values obtained from Fisher’s exact test.

## Data availability

The first draft of the APR data from this work can be found at https://www.mbru.ac.ae/the-arab-pangenome-reference/. We hope that the downloadable pangenome will impact the research and clinical genetics communities. We have submitted our sequencing data (including PacBio, ONT) for all the samples to the global Sequence Read Archive (SRA) repository that can be openly accessed and downloaded under the accession no PRJNA1108179.

## Code availability

The code used to reproduce the pangenome from this work can be found at GitHub (https://github.com/muddinmbru/arab_pangenome_reference) and PanScan is available at (https://github.com/muddinmbru/panscan). The relevant commands used in other analyses can be found in the Methods or Supplementary Information.

## Acknowledgements

We would like to thank all the study participants for giving us this opportunity of building a reference pangenome. Yehia Zakaria Kotp provided constant support for the CATG Flamingo computation cluster. Avinash Krishnan, and Freeda Pinto provided constant help with logistics at CATG sequencing lab. HPRC team for commenting on our assembly data quality. Dubai Academic Healthcare Corporation (DAHC) funded this work and supported the procurement of reagents, chemicals, and computing hardwares. The funders had no role in the design of the study, data collection, analysis, publish or preparation of the manuscript. We thank Nature Proofediting Services for its linguistic assistance, which greatly aided the preparation of this manuscript.

## Author contributions

AAA, MU, SDP and MA conceived and designed the study. AAA, MU were responsible for ethical, legal and social implications. AAA, MU, NN coordinated and supervised the project. BJ, NN, BA, MAO, MHSA, AA, OZSA, DFY, HAS, HHK, were responsible for recruitment and collection of blood samples and clinical data from medical health records. AAA, MU, NN, SH, MA, HHK, BJ were responsible for sample selection and population genetic analysis. MAO, MU, NN, NM, HE, DK, DS, were responsible for DNA and RNA sequencing. MK, BB, SH, NN, MU were responsible for variant annotation. MK was responsible for assembly creation and MK,MU, NN were responsible for assembly quality control and assembly reliability analysis. MU, MK, BB, NN, SH were responsible for variant detection from assembly, pangenome empirical analysis and quality control. BB, SH, MU, NN were responsible for pangenome applications of SVs, short and long-read mapping. MK, NN, MU, NK, SS were responsible for pangenome visualization and complex loci analysis and pangenome graph creation. MU, MK, BB, NN, SH statistical analysis. AAA, MU, NN, SDP, MA, AAT were responsible for manuscript writing. MU, AAA, NN were responsible for manuscript editing. AAA, MU, and NN were responsible for data coordination and management.

## Ethics declarations

### Supplementary information

Supplementary document: Includes methods of APR cohort and sample phenotype information, Pacific Bioscience High Fidelity (HiFi) sequencing, Nanopore ultra-long sequencing, Hi-C sequencing, exome sequencing, variant identification and filtering, and *de novo* genome assembly. Includes Supplementary Figures S1–S23.

Supplementary Tables: Table 1 – 34.

## References

1. Wang, T., Antonacci-Fulton, L., Howe, K., Lawson, H.A., Lucas, J.K., Phillippy, A.M., Popejoy, A.B., Asri, M., Carson, C., Chaisson, M.J.P., et al. (2022). The Human Pangenome Project: a global resource to map genomic diversity. Nature 604, 437–446. 10.1038/s41586-022-04601-8.

2. 1000 Genomes Project Consortium, Auton, A., Brooks, L.D., Durbin, R.M., Garrison, E.P., Kang, H.M., Korbel, J.O., Marchini, J.L., McCarthy, S., McVean, G.A., et al. (2015). A global reference for human genetic variation. Nature 526, 68–74. 10.1038/nature15393.

3. Bergström, A., McCarthy, S.A., Hui, R., Almarri, M.A., Ayub, Q., Danecek, P., Chen, Y., Felkel, S., Hallast, P., Kamm, J., et al. (2020). Insights into human genetic variation and population history from 929 diverse genomes. Science 367. 10.1126/science.aay5012.

4. Nurk, S., Koren, S., Rhie, A., Rautiainen, M., Bzikadze, A. V, Mikheenko, A., Vollger, M.R., Altemose, N., Uralsky, L., Gershman, A., et al. (2022). The complete sequence of a human genome. Science 376, 44–53. 10.1126/science.abj6987.

5. Rhie, A., Nurk, S., Cechova, M., Hoyt, S.J., Taylor, D.J., Altemose, N., Hook, P.W., Koren, S., Rautiainen, M., Alexandrov, I.A., et al. (2023). The complete sequence of a human Y chromosome. Nature 621, 344–354. 10.1038/s41586-023-06457-y.

6. Liao, W.-W., Asri, M., Ebler, J., Doerr, D., Haukness, M., Hickey, G., Lu, S., Lucas, J.K., Monlong, J., Abel, H.J., et al. (2023). A draft human pangenome reference. Nature 617, 312–324. 10.1038/s41586-023-05896-x.

7. Gao, Y., Yang, X., Chen, H., Tan, X., Yang, Z., Deng, L., Wang, B., Kong, S., Li, S., Cui, Y., et al. (2023). A pangenome reference of 36 Chinese populations. Nature 619, 112–121. 10.1038/s41586-023-06173-7.

8. Popejoy, A.B., and Fullerton, S.M. (2016). Genomics is failing on diversity. Nature 538, 161–164. 10.1038/538161a.

9. Almarri, M.A., Haber, M., Lootah, R.A., Hallast, P., Al Turki, S., Martin, H.C., Xue, Y., and Tyler-Smith, C. (2021). The genomic history of the Middle East. Cell 184, 4612–4625.e14. 10.1016/j.cell.2021.07.013.

10. Mbarek, H., Devadoss Gandhi, G., Selvaraj, S., Al-Muftah, W., Badji, R., Al-Sarraj, Y., Saad, C., Darwish, D., Alvi, M., Fadl, T., et al. (2022). Qatar genome: Insights on genomics from the Middle East. Hum Mutat 43, 499–510. 10.1002/humu.24336.

11. Tadmouri, G.O., Nair, P., Obeid, T., Al Ali, M.T., Al Khaja, N., and Hamamy, H.A. (2009). Consanguinity and reproductive health among Arabs. Reprod Health 6, 17. 10.1186/1742-4755-6-17.

12. Teebi, A.S. (1994). Autosomal recessive disorders among Arabs: an overview from Kuwait. J Med Genet 31, 224–233. 10.1136/jmg.31.3.224.

13. Al-Gazali, L., Hamamy, H., and Al-Arrayad, S. (2006). Genetic disorders in the Arab world. BMJ 333, 831–834. 10.1136/bmj.38982.704931.AE.

14. Rahim, H.F.A., Sibai, A., Khader, Y., Hwalla, N., Fadhil, I., Alsiyabi, H., Mataria, A., Mendis, S., Mokdad, A.H., and Husseini, A. (2014). Non-communicable diseases in the Arab world. Lancet 383, 356–367. 10.1016/S0140-6736(13)62383-1.

15. El-Kebbi, I.M., Bidikian, N.H., Hneiny, L., and Nasrallah, M.P. (2021). Epidemiology of type 2 diabetes in the Middle East and North Africa: Challenges and call for action. World J Diabetes 12, 1401–1425. 10.4239/wjd.v12.i9.1401.

16. Sherry, S.T., Ward, M.H., Kholodov, M., Baker, J., Phan, L., Smigielski, E.M., and Sirotkin, K. (2001). dbSNP: the NCBI database of genetic variation. Nucleic Acids Res 29, 308–311. 10.1093/nar/29.1.308.

17. Karczewski, K.J., Francioli, L.C., Tiao, G., Cummings, B.B., Alföldi, J., Wang, Q., Collins, R.L., Laricchia, K.M., Ganna, A., Birnbaum, D.P., et al. (2020). The mutational constraint spectrum quantified from variation in 141,456 humans. Nature 581, 434–443. 10.1038/s41586-020-2308-7.

18. Scott, E.M., Halees, A., Itan, Y., Spencer, E.G., He, Y., Azab, M.A., Gabriel, S.B., Belkadi, A., Boisson, B., Abel, L., et al. (2016). Characterization of Greater Middle Eastern genetic variation for enhanced disease gene discovery. Nat Genet 48, 1071– 1076. 10.1038/ng.3592.

19. MacDonald, J.R., Ziman, R., Yuen, R.K.C., Feuk, L., and Scherer, S.W. (2014). The Database of Genomic Variants: a curated collection of structural variation in the human genome. Nucleic Acids Res 42, D986–92. 10.1093/nar/gkt958.

20. Lazaridis, I., Nadel, D., Rollefson, G., Merrett, D.C., Rohland, N., Mallick, S., Fernandes, D., Novak, M., Gamarra, B., Sirak, K., et al. (2016). Genomic insights into the origin of farming in the ancient Near East. Nature 2016 536:7617 *536*, 419–424. 10.1038/nature19310.

21. Lazaridis, I., Patterson, N., Mittnik, A., Renaud, G., Mallick, S., Kirsanow, K., Sudmant, P.H., Schraiber, J.G., Castellano, S., Lipson, M., et al. (2014). Ancient human genomes suggest three ancestral populations for present-day Europeans. Nature 2014 513:7518 *513*, 409–413. 10.1038/nature13673.

22. Patterson, N., Moorjani, P., Luo, Y., Mallick, S., Rohland, N., Zhan, Y., Genschoreck, T., Webster, T., and Reich, D. (2012). Ancient admixture in human history. Genetics 192, 1065–1093. 10.1534/GENETICS.112.145037/-/DC1/GENETICS.112.145037-1.PDF.

23. Lawson, D.J., Hellenthal, G., Myers, S., and Falush, D. Inference of population structure using dense haplotype data.

24. Cheng, H., Concepcion, G.T., Feng, X., Zhang, H., and Li, H. (2021). Haplotype-resolved de novo assembly using phased assembly graphs with hifiasm. Nat Methods 18, 170–175. 10.1038/s41592-020-01056-5.

25. Rautiainen, M., Nurk, S., Walenz, B.P., Logsdon, G.A., Porubsky, D., Rhie, A., Eichler, E.E., Phillippy, A.M., and Koren, S. (2023). Telomere-to-telomere assembly of diploid chromosomes with Verkko. Nat Biotechnol 41, 1474–1482. 10.1038/s41587-023-01662-6.

26. Wood, D.E., and Salzberg, S.L. (2014). Kraken: ultrafast metagenomic sequence classification using exact alignments. Genome Biol 15, R46. 10.1186/gb-2014-15-3-r46.

27. Li, H. (2021). New strategies to improve minimap2 alignment accuracy. Bioinformatics 37, 4572–4574. 10.1093/BIOINFORMATICS/BTAB705.

28. Chen, Y., Zhang, Y., Wang, A.Y., Gao, M., and Chong, Z. (2021). Accurate long-read de novo assembly evaluation with Inspector. Genome Biol 22, 312. 10.1186/s13059-021-02527-4.

29. GitHub - mobinasri/flagger: Evaluating genome assemblies https://github.com/mobinasri/flagger.

30. GitHub - shilpagarg/pstools https://github.com/shilpagarg/pstools.

31. Bzikadze, A. V, and Pevzner, P.A. (2023). UniAligner: a parameter-free framework for fast sequence alignment. Nat Methods 20, 1346–1354. 10.1038/s41592-023-01970-4.

32. Shumate, A., and Salzberg, S.L. (2021). Liftoff: accurate mapping of gene annotations. Bioinformatics 37, 1639–1643. 10.1093/bioinformatics/btaa1016.

33. McFarlane, C., Kelvin, A.A., de la Vega, M., Govender, U., Scott, C.J., Burrows, J.F., and Johnston, J.A. (2010). The deubiquitinating enzyme USP17 is highly expressed in tumor biopsies, is cell cycle regulated, and is required for G1-S progression. Cancer Res 70, 3329–3339. 10.1158/0008-5472.CAN-09-4152.

34. Komander, D., Clague, M.J., and Urbé, S. (2009). Breaking the chains: structure and function of the deubiquitinases. Nat Rev Mol Cell Biol 10, 550–563. 10.1038/nrm2731.

35. Luse, D.S. (2014). The RNA polymerase II preinitiation complex. Through what pathway is the complex assembled? Transcription 5, e27050. 10.4161/trns.27050.

36. Chen, C.-L., Lee, N.-C., Chien, Y.-H., Hwu, W.-L., Hung, M.-Z., Lin, Y.-L., Lin, S.-Y., and Lee, C.-N. (2023). Ethnically unique disease burden and limitations of current expanded carrier screening panels. Int J Gynaecol Obstet. 10.1002/ijgo.15072.

37. Hickey, G., Monlong, J., Ebler, J., Novak, A.M., Eizenga, J.M., Gao, Y., Human Pangenome Reference Consortium, Marschall, T., Li, H., and Paten, B. (2023). Pangenome graph construction from genome alignments with Minigraph-Cactus. Nat Biotechnol. 10.1038/s41587-023-01793-w.

38. GitHub - marschall-lab/panacus: Panacus is a tool for computing statistics for GFA-formatted pangenome graphs https://github.com/marschall-lab/panacus.

39. Kern, C.H., Yang, M., and Liu, W.-S. (2021). The PRAME family of cancer testis antigens is essential for germline development and gametogenesis†. Biol Reprod 105, 290–304. 10.1093/biolre/ioab074.

40. Proshkin, S.A., Shematorova, E.K., and Shpakovski, G. V (2019). The Human Isoform of RNA Polymerase II Subunit hRPB11bα Specifically Interacts with Transcription Factor ATF4. Int J Mol Sci 21. 10.3390/ijms21010135.

41. Chauhan, S., Zheng, X., Tan, Y.Y., Tay, B.H., Lim, S., Venkatesh, B., and Kaldis, P. (2012). Evolution of the Cdk-activator Speedy/RINGO in vertebrates. Cell Mol Life Sci 69, 3835–3850. 10.1007/S00018-012-1050-1.

42. Frankish, A., Diekhans, M., Jungreis, I., Lagarde, J., Loveland, J.E., Mudge, J.M., Sisu, C., Wright, J.C., Armstrong, J., Barnes, I., et al. (2021). GENCODE 2021. Nucleic Acids Res 49, D916–D923. 10.1093/nar/gkaa1087.

43. Ebert, P., Audano, P.A., Zhu, Q., Rodriguez-Martin, B., Porubsky, D., Bonder, M.J., Sulovari, A., Ebler, J., Zhou, W., Serra Mari, R., et al. (2021). Haplotype-resolved diverse human genomes and integrated analysis of structural variation. Science 372. 10.1126/science.abf7117.

44. Poplin, R., Chang, P.-C., Alexander, D., Schwartz, S., Colthurst, T., Ku, A., Newburger, D., Dijamco, J., Nguyen, N., Afshar, P.T., et al. (2018). A universal SNP and small-indel variant caller using deep neural networks. Nat Biotechnol 36, 983–987. 10.1038/nbt.4235.

45. Edge, P., and Bansal, V. (2019). Longshot enables accurate variant calling in diploid genomes from single-molecule long read sequencing. Nature Communications 2019 10:1 *10*, 1–10. 10.1038/s41467-019-12493-y.

46. Sirén, J., Monlong, J., Chang, X., Novak, A.M., Eizenga, J.M., Markello, C., Sibbesen, J.A., Hickey, G., Chang, P.-C., Carroll, A., et al. (2021). Pangenomics enables genotyping of known structural variants in 5202 diverse genomes. Science 374, abg8871. 10.1126/science.abg8871.

47. Logsdon, G.A., Vollger, M.R., Hsieh, P., Mao, Y., Liskovykh, M.A., Koren, S., Nurk, S., Mercuri, L., Dishuck, P.C., Rhie, A., et al. (2021). The structure, function and evolution of a complete human chromosome 8. Nature 593, 101–107. 10.1038/s41586-021-03420-7.

48. Aamer, W., Al-Maraghi, A., Syed, N., Gandhi, G.D., Aliyev, E., Al-Kurbi, A.A., Al-Saei, O., Kohailan, M., Krishnamoorthy, N., Palaniswamy, S., et al. (2024). Burden of Mendelian disorders in a large Middle Eastern biobank. Genome Med 16, 46. 10.1186/s13073-024-01307-6.

49. Vollger, M.R., Dishuck, P.C., Harvey, W.T., DeWitt, W.S., Guitart, X., Goldberg, M.E., Rozanski, A.N., Lucas, J., Asri, M., Human Pangenome Reference Consortium, et al. (2023). Increased mutation and gene conversion within human segmental duplications. Nature 617, 325–334. 10.1038/s41586-023-05895-y.

50. Ravarani, C.N.J., Flock, T., Chavali, S., Anandapadamanaban, M., Babu, M.M., and Balaji, S. (2020). Molecular determinants underlying functional innovations of TBP and their impact on transcription initiation. Nat Commun 11, 2384. 10.1038/s41467-020-16182-z.

51. Mallick, S., Micco, A., Mah, M., Ringbauer, H., Lazaridis, I., Olalde, I., Patterson, N., and Reich, D. (2024). The Allen Ancient DNA Resource (AADR) a curated compendium of ancient human genomes. Scientific Data 2024 11:1 *11*, 1–10. 10.1038/s41597-024-03031-7.

52. Zheng, X., Levine, D., Shen, J., Gogarten, S.M., Laurie, C., and Weir, B.S. (2012). A high-performance computing toolset for relatedness and principal component analysis of SNP data. Bioinformatics 28, 3326–3328. 10.1093/bioinformatics/bts606.

53. Alexander, D.H., and Lange, K. (2011). Enhancements to the ADMIXTURE algorithm for individual ancestry estimation. BMC Bioinformatics 12, 246. 10.1186/1471-2105-12-246.

54. Lawson, D.J., Hellenthal, G., Myers, S., and Falush, D. (2012). Inference of population structure using dense haplotype data. PLoS Genet 8, e1002453. 10.1371/journal.pgen.1002453.

55. Weissensteiner, H., Pacher, D., Kloss-Brandstätter, A., Forer, L., Specht, G., Bandelt, H.- J., Kronenberg, F., Salas, A., and Schönherr, S. (2016). HaploGrep 2: mitochondrial haplogroup classification in the era of high-throughput sequencing. Nucleic Acids Res 44, W58–63. 10.1093/nar/gkw233.

56. Fähnrich, A., Stephan, I., Hirose, M., Haarich, F., Awadelkareem, M.A., Ibrahim, S., Busch, H., and Wohlers, I. (2023). North and East African mitochondrial genetic variation needs further characterization towards precision medicine. J Adv Res 54, 59–76. 10.1016/j.jare.2023.01.021.

57. Aljasmi, F.A., Vijayan, R., Sudalaimuthuasari, N., Souid, A.-K., Karuvantevida, N., Almaskari, R., Mohammed Abdul Kader, H., Kundu, B., Michel Hazzouri, K., and Amiri, K.M.A. (2020). Genomic Landscape of the Mitochondrial Genome in the United Arab Emirates Native Population. Genes (Basel) 11. 10.3390/genes11080876.

58. Chen, H., Lu, Y., Lu, D., and Xu, S. (2021). Y-LineageTracker: a high-throughput analysis framework for Y-chromosomal next-generation sequencing data. BMC Bioinformatics 22, 114. 10.1186/s12859-021-04057-z.

59. Hallast, P., Agdzhoyan, A., Balanovsky, O., Xue, Y., and Tyler-Smith, C. (2021). A Southeast Asian origin for present-day non-African human Y chromosomes. Hum Genet 140, 299–307. 10.1007/s00439-020-02204-9.

60. Elliott, K.S., Haber, M., Daggag, H., Busby, G.B., Sarwar, R., Kennet, D., Petraglia, M., Petherbridge, L.J., Yavari, P., Heard-Bey, F.U., et al. (2022). Fine-Scale Genetic Structure in the United Arab Emirates Reflects Endogamous and Consanguineous Culture, Population History, and Geography. Mol Biol Evol 39. 10.1093/molbev/msac039.

61. Abu-Amero, K.K., Hellani, A., González, A.M., Larruga, J.M., Cabrera, V.M., and Underhill, P.A. (2009). Saudi Arabian Y-Chromosome diversity and its relationship with nearby regions. BMC Genet 10, 59. 10.1186/1471-2156-10-59.

62. Mikheenko, A., Prjibelski, A., Saveliev, V., Antipov, D., and Gurevich, A. (2018). Versatile genome assembly evaluation with QUAST-LG. Bioinformatics 34, i142–i150. 10.1093/bioinformatics/bty266.

63. Li, H., Feng, X., and Chu, C. (2020). The design and construction of reference pangenome graphs with minigraph. Genome Biol 21, 265. 10.1186/s13059-020-02168-z.

64. Li, H. (2019). Identifying centromeric satellites with dna-brnn. Bioinformatics 35, 4408–4410. 10.1093/bioinformatics/btz264.

65. Hickey, G., Paten, B., Earl, D., Zerbino, D., and Haussler, D. (2013). HAL: a hierarchical format for storing and analyzing multiple genome alignments. Bioinformatics 29, 1341–1342. 10.1093/bioinformatics/btt128.

66. Hao, Z., Lv, D., Ge, Y., Shi, J., Weijers, D., Yu, G., and Chen, J. (2020). RIdeogram: drawing SVG graphics to visualize and map genome-wide data on the idiograms. PeerJ Comput Sci 6, e251. 10.7717/peerj-cs.251.

67. Wick, R.R., Schultz, M.B., Zobel, J., and Holt, K.E. (2015). Bandage: interactive visualization of de novo genome assemblies. Bioinformatics 31, 3350–3352. 10.1093/bioinformatics/btv383.

68. Rautiainen, M., and Marschall, T. (2020). GraphAligner: rapid and versatile sequence-to-graph alignment. Genome Biol 21, 253. 10.1186/s13059-020-02157-2.

69. Li, H. (2018). Minimap2: pairwise alignment for nucleotide sequences. Bioinformatics 34, 3094–3100. 10.1093/bioinformatics/bty191.

70. Garrison, E., Guarracino, A., Heumos, S., Villani, F., Bao, Z., Tattini, L., Hagmann, J., Vorbrugg, S., Marco-Sola, S., Kubica, C., et al. (2023). Building pangenome graphs. bioRxiv. 10.1101/2023.04.05.535718.

