## Supplemental methods and figures for "A draft Arab pangenome reference"

### Supplementary Methods

#### A. APR cohort and sample phenotype information

We recruited samples from the Dubai Academic Healthcare Corporation (DAHC), including a trio from Al Jalila Children’s Hospital and 50 unrelated healthy individuals from various DAHC facilities. The cohort comprised of samples from eight Arab populations (Saudi Arabia, Egypt, Jordan, Syria, Oman Morocco, Yemen and the UAE) from the Middle East and North Africa (MENA) region. These individuals underwent extensive clinical assessments (Supplementary Table 1, Supplementary Figure 1) by family physicians and were confirmed to be free of common chronic diseases such as diabetes, hypertension, and cancer. This assessment included reviewing multi-year health records for adult participants and conducting interviews by a genetic counselor. At the time of recruitment, all participants were deemed healthy, with normal laboratory test results, although some exhibited elevated serum cholesterol levels (Supplementary Figure 1). All participants underwent extensive phenotyping, including clinical, biochemical, and anthropometric assessments as well as review of their electronic health records. Through verbal interviews, 30% individuals from UAE confirmed migratory history of their recent ancestors from other Arab or Middle Eastern countries (for details see Supplementary Table 34).

#### B. Pacific Bioscience High Fidelity (HiFi) Sequencing

We constructed whole genome sequencing (WGS) libraries using the long-read PacBio SMRTbell prep kit 3.0 protocol (102-166-600) as detailed below.

##### a. DNA quality control

To evaluate and ensure suitable quality and size of DNA for the use of this protocol, we began by homogenizing the DNA in the solution by pulse vortexing and gently pipetting our samples. Quick spinning the samples, we took 1 µL aliquot from each sample and diluted them with 9 µL of elution buffer or water. Next, we measured the DNA concentration using the 1X dsDNA HS kit and the Qubit fluorometer. Each aliquot was diluted in Tape Station dilution buffer to a total concentration of 250 pg/µL, based on the Qubit readings. DNA size was measured using the Femto Pulse system and the gDNA 165Kb analysis kit.

##### b. DNA shearing and cleanup

Low TE buffer was added to bring the volume of DNA samples to 130 µL, with a target concentration of 30 ng/µL. The DNA was sheared on the Megaruptor 3 system as per the recommended settings. Sheared DNA was transferred into a tube, and 1 X v/v of resuspended, room-temperature SMRTbell cleanup beads were added. The beads were pipette mixed until evenly distributed. Tubes were then quick-spinned and incubated at room temperature for 10 min. After incubation, tubes were placed in a magnetic separation rack to allow the beads to separate fully, and the supernatant was discarded. The beads were slowly covered by adding 200 µL of fresh 80% ethanol. After 30 seconds, the ethanol was discarded, and the step was repeated.

To remove the residual ethanol, the tubes were removed from the magnetic rack, spun, placed again in the rack until the beads separated, and the remaining solution was aspirated. Immediately after removing the tubes from the magnetic rack, 47 µL of low TE buffer were added to the tubes and the beads were resuspended by pipetting 10 times. The tubes were quick-spinned and incubated at room temperature for 5 min for DNA elution. Next, the tubes were placed again in the magnetic rack, beads separated, and the supernatant transferred to a new tube. To verify sample quality per its concentration and size distribution, we aliquoted 1 µL from each sample and diluted with 9 µL elution buffer or water. Next, the Qubit fluorometer was used to measure DNA concentration using the 1X dsDNA HS kit. Each aliquot was diluted to 250 pg/µL in Tape Station dilution buffer and DNA size was measured using the Tape Station system.

##### c. Repair and A-tailing

Into a centrifuge tube, we created reaction mix 1 by adding repair buffer (35.2 µL for 4 libraries, 80 µL for 8 libraries), end repair mix (17.6 µL for 4 libraries, 40 µL for 8 libraries), and DNA repair mix (8.8 µL for 4 libraries, 30 µL for 8 libraries). After pipetting and spinning the reaction mix, we added 14 µL of reaction mix 1 to each of our samples, bringing the total reaction volume to 60 µL. Then we pipetted and spinned our sample reactions, followed by running the repair and A-tailing thermocycling program, 37°C for 30 min, 65°C for 5 min, and a 4°C hold.

##### d. Adapter ligation

Since we barcoded our samples, we added 4 µL of barcoded adapters from the SMRTbell barcoded adapter plate 3.0 to each sample. In a centrifuge tube, we created reaction mix 2 by adding ligation mix (132 µL for 4 libraries, 300 µL for 8 libraries), and ligation enhancer (4.4 µL for 4 libraries, 10 µL for 8 libraries). After pipetting and spinning the reaction mix, we added 31 µL of reaction mix 2 to each of our samples, bringing the total reaction volume to 95 µL. Then we pipetted and spinned our sample reactions, followed by running the adapter ligation thermocycling program, 20°C for 30 min and a 4°C hold.

##### e. Cleanup

95 µL of resuspended SMRTbell cleanup beads were added to each sample and pipetted for even distribution. The tubes were spun and incubated at room temperature for 10 min. Tubes were then placed in a magnetic separation rack, when the beads separated from the solution the supernatant was discarded. The beads were slowly covered by adding 200 µL of fresh 80% ethanol. After 30 seconds, the ethanol was discarded, and the step was repeated. To remove the residual ethanol, the tubes were removed from the magnetic rack, spun, placed again in the rack until the beads separated, and the remaining solution was aspirated. Immediately after removing the tubes from the magnetic rack, 40 µL of elution buffer were added to the tubes and the beads were resuspended. The tubes were quick-spinned and incubated at room temperature for 5 min for DNA elution. Next, the tubes were placed again in the magnetic rack, beads separated, and the supernatant transferred to a new tube.

##### f. Nuclease treatment

In a centrifuge tube, we created reaction mix 3 by adding nuclease buffer (22 µL for 4 libraries, 50 µL for 8 libraries), and nuclease mix (22 µL for 4 libraries, 50 µL for 8 libraries). After pipetting and spinning the reaction mix, we added 10 µL of reaction mix 3 to each of our samples, bringing the total reaction volume to 50 µL. Then we pipetted and spinned our sample reactions, followed by running the nuclease treatment thermocycling program, 37°C for 15 min and a 4°C hold.

##### g. AMPure PB bead size selection

By adding 1.75 mL of resuspended AMPure PB beads to 3.25 mL of elution buffer, we made a 35% v/v dilution. Then we added 155 µL (3.1 X v/v) of resuspended 35% AMPure PB beads to each of our samples.

##### h. Binding, washing, and eluting

The samples were pipette mixed and spun, then incubated at room temperature for 20 minutes. Tubes were then placed in a magnetic separation rack, when the beads separated from the solution the supernatant was discarded. The beads were slowly covered by adding 200 µL of fresh 80% ethanol. After 30 seconds, the ethanol was discarded, and the step was repeated. To remove the residual ethanol, the tubes were removed from the magnetic rack, spun, placed again in the rack until the beads separated, and the remaining solution was aspirated. Immediately after removing the tubes from the magnetic rack, 15 µL of elution buffer were added to the tubes and the beads were resuspended by pipetting 10 times. The tubes were quick-spinned and incubated at room temperature for 5 min for DNA elution. Next, the tubes were placed again in the magnetic rack, beads separated, and the supernatant transferred to a new tube. From each tube, a 1 µL aliquot is taken and diluted with 9 µL of elution buffer or water. DNA concentration was measured using a Qubit fluorometer and the 1X dsDNA HS kit.

##### i. ABC Steps

After completing library preparation, we performed ABC sequencing preparation as per the (102-739-700) kit protocol, detailed below.

###### i. Annealing Sequencing primer

Following the DNA quantification, we diluted our samples to a concentration of 30 ng/µL. In a new Lo-bind tube, we added 11 µL of sample, 5/5 µL of annealing buffer, and 5.5 µL of sequencing primer, bringing the total volume to 22 µL. After pipetting well, the tubes were incubated at room temperature for 15 min.

###### ii. Sequencing polymerase

We began by diluting 11.5 µL of sequencing polymerase in 563.5 µL of polymerase buffer in a new lo-bind tube, bringing the total volume to 575 µL diluted polymerase master mix to be used immediately. Next, for each of our samples, we added 22 µL annealed sample and 22 µL diluted polymerase, bringing the total volume to 44 µL. The tubes were then incubated at room temperature for 15 min.

###### iii. Purification of Polymerase bound SMRTbell complexes

SMRTbell cleanup beads and loading buffer were brought to room temperature. For each sample, 44 µL of Binding reaction and 56 µL of dilution buffer were added, bringing the total reaction to 100 µL. Next, 120 µL of SMRTbell cleanup beads were added to the samples, gently pipetted, and incubated at room temperature for 10 min. Tubes were then placed in a magnetic bead rack until the beads completely separated from the solution. The supernatant was discarded, and the beads were immediately resuspended in 50 µL loading buffer and gently pipetted. For elution of the polymerase-bound complexes, the samples were incubated at room temperature for at least 5 min. Tubes were placed again on the magnetic rack until the beads separated. Then the eluted solution was transferred to new Lo-Bind tubes and placed on ice, protected from the light.

###### iv. Sequencing control dilution

The control underwent three dilution steps, producing a master mix that is enough for all our samples. In a new Lo-bind tube, 19 µL dilution buffer and 1 µL sequencing control were mixed by flicking the tube and pulse spinning, then kept on ice. This process was repeated to achieve dilution 2. For the third and final dilution, 76 µL dilution buffer and 4 µL diluted sequencing control from the second dilution were mixed well by flicking and spinning and kept on ice.

###### v. Loading dilution

For each of the samples loaded on the Revio system, Revio Polymerase Kit was used (Serial Number: 102-739-700). We combined 50 µL of the prepared sample, 47 µL loading buffer, and 3 µL of the diluted sequencing control from the previous step, bringing the total volume to 100 µL. 95 µL of each sample were loaded per well.

#### C. Nanopore ultra-long sequencing

We used Oxford Nanopore Technologies ultra-long read sequencing kit (SQK-ULK114) protocol. Each sample was aliquoted at the time of collection into 1.8 ml Cryovials and stored at -80°C. 3 aliquots of each sample was used for preparing ultra-long libraries generating about 50× coverage of unsheared sequencing from 3 PromethION flow cells (R10.4.1) and a N50 value of around 60 kb. We have used additional flow cells (see Supplementary Table 2) for samples that did not yield adequate data.

Ultra-long libraries were prepared using the protocol outlined below:

##### a. PBMC isolation from frozen blood

1.6 ml frozen blood was thawed in 3X volume of cold NEB RBC lysis buffer in a 15 ml Falcon tube. Tube was gently inverted 10 times and kept at 4°C for 5 minutes. The tube was centrifuged at 2000 X g at 4°C for 2 min, after which the supernatant was discarded. The pellet was resuspended in 1.6 ml 1X PBS buffer by gently flicking and pipetting. The above steps were repeated for a total of 3 washes with NEB RBC lysis buffer. The final pellet of around 60 million cells was resuspended in 40 µl of 1X PBS and carried forward for ultra-long DNA extraction.

##### b. DNA extraction

Resuspended cells in 40ul of PBS were transferred to a fresh 5 ml Lo-Bind Eppendorf tube. In a separate tube, 1.8 ml NEB Monarch tissue lysis buffer and 40ul of NEB Monarch Proteinase K was mixed, then added to the resuspended PBMCs in the Lo-Bind tube. The reaction was mixed by slow pipetting five times using a 1 ml wide-bore pipette tip and incubated at 56°C on a thermomixer for 10 min. 15ul of NEB Monarch RNase A was added to the reaction and mixed by slow pipetting five times using a 1 ml wide-bore pipette tip. The reaction was incubated at 56°C on a thermomixer for 10 min at 650 rpm. Next, 900 µl of NEB Monarch protein separation solution was added to the reaction and the tube was placed in a Hula mixer for 10 mins at 3 rpm. The 5 ml tubes were then centrifuged for phase separation at 16,000 X g at 4°C for 10 min. Upper phase containing ultra-long DNA fragments was carefully collected into a fresh 5 ml Lo-Bind Eppendorf tube using a 1 ml wide-bore pipette tip. To this, 3 glass beads were added along with 2.5 ml isopropanol. The tube was placed in a Hula mixer for 20 mins at 3 rpm.

The ultra-long DNA fragments were precipitated around the 2 glass beads by the end of the incubation. The supernatant was discarded, and the glass beads were washed two times using 2ml NEB Monarch wash buffer per wash. The glass beads were then introduced to a Monarch bead retainer inserted into a Monarch collection tube and centrifuged at 1000 X g for 1 minute to remove any remaining wash buffer. The glass beads were immediately moved into a fresh 2 ml Lo-Bind Eppendorf tube containing 560 µl of ONT extraction elution buffer. The tube was incubated at 56°C for 10 min. The beads and the elution buffer were introduced into a clean bead retainer inserted into a Monarch collection tube and centrifuged at 1000 X g for 1 minute. The eluate was collected in the collection tube. The beads were discarded. To the collection tube, 200 µl of Oxford Nanopore Technologies extraction elution buffer (EEB) was added and the total volume of 760 µl of eluant was transferred to a fresh 1.5 ml Lo-Bind Eppendorf tube. The reaction was incubated at 56°C for 10 min. The final eluate of UHMW DNA was mixed by slow pipetting five times using a 1ml wide-bore pipette tip and stored overnight at room temperature.

##### c. DNA Tagmentation

In a 1.5 ml Lo-Bind Eppendorf tube, 5 µl of ONT ULK fragmentation mix (FRA) and 245 µl of FRA dilution buffer (FDB) were mixed by pipetting. 250 µl of the diluted fragmentation mix was added to 760 µl of UHMW DNA. The reaction was immediately mixed by slow pipetting ten times using a 1 ml wide-bore pipette tip. The reaction was incubated at room temperature for 10 min, followed by incubation on a thermomixer at 75°C for 10 min, and finally cooled on ice for 15 min.

##### j. Adapter attachment

5ul of rapid adapter (RA) was added to the DNA reaction and gently mixed by slow pipetting five times using a 1 ml wide-bore pipette tip. The reaction was incubated at room temperature for 30 min.

##### k. Clean-up

A metal precipitation star (PS) was added to the adapted DNA for clean-up. 500 µl of precipitation buffer (PTB) was added to the reaction and mixed by rotating on a Hula mixer for 20 min at 3 rpm. The adapted ultra-long DNA was precipitated around the precipitation star and the supernatant was discarded. The tube was briefly spun to remove any remaining supernatant. 300 µl of elution buffer (EB) was added to the tube containing the metal star and DNA. The tube was stored overnight at room temperature. Using a 1ml wide-bore pipette tip the ultra-long DNA library was removed and retained in a fresh 1.5 ml Lo-Bind Eppendorf tube. The tube containing the precipitation star was briefly spun and any remaining eluate was transferred to the tube containing the final ultra-long DNA library. The final library was gently mixed by slow pipetting five times using a 1 ml wide-bore pipette tip.

##### l. Flow cell loading and sequencing

ONT sequencing buffer (SQB) (100 µl) and ONT loading solution (10 µl) were added to 90 µl of the ultra-long DNA library from above. The mixture was gently mixed by slow pipetting five times using a wide-bore pipette tip. Libraries were then incubated at room temperature for 30 min. Next, the libraries were gently mixed by slow pipetting with a wide-bore tip to ensure homogeneity. Before loading the library, the flow cell was primed with flush buffer/flush tether mixture per ONT directions. The library was then added to the flow cell. The mixture was viscous and loaded in a drop-wise manner using a 1 ml wide-bore tip. The sequencing run had a pore scan time set for every 1.5h and a minimum read length set to 1000 bp. Live base calling using the high accuracy (HAC) basecalling model with modified base detection for 5mC was performed on MinKNOW.

#### D. Hi-C sequencing

##### a. Crosslinking Chromatin in Cultured Cells

We constructed Hi-C libraries using the Qiagen EpiTect Hi-C Kit protocol Kit with significant modifications as detailed below.

PBMCs were isolated from approximately 1 ml of frozen whole blood, with cell counts ranging from 5x10^3 to 2.5x10^6 cells per sample. The isolation process involved pelleting the cells from 1ml of thawed blood using a 10× volume of cold RBC lysis buffer. Tube was gently inverted 10 times to ensure thorough mixing and then incubated for 5 minutes at room temperature, with gentle inversion twice during the incubation period. Following this, centrifugation was carried out at 500 x g for 5 minutes at 4°C to pellet the white blood cells, and the supernatant was completely discarded. Next, 980 μl of 1X PBS and 20 μl of FBS (2%) were added, followed by centrifugation at 2500 × g for 5 minutes and subsequent discarding of the supernatant to remove any remaining debris. The pellet was then resuspended in 1960 μl of ice-cold PBS and 40 μl of FBS. 54 μl of 37% formaldehyde was added dropwise to ensure crosslinking of cells and preservation of cellular integrity. The mixture was then placed in a HulaMixer at 3 rpm for 10 minutes at room temperature. Afterwards, crosslinking was quenched using 1 ml of 3 M Tris pH 7.5 and the solution was mixed by inverting 5 times. After incubating for 5 minutes at room temperature and then on ice for at least 15 minutes, the cells were centrifuged for 10 minutes at 800 × g at 4°C, and the supernatant was removed. This was followed by an additional washing step with PBS to ensure removal of excess fixative. 1 ml of PBS was added and centrifuged at 800 g for 5 minutes. Finally, the cell pellet was resuspended in 50 μl of cold PBS.

##### b. Preparing Hi-C Library from fixed cells

###### i. Cell Lysis

Cell pellet was mixed with 150 μL of ice-cold RNase-Free Water, followed by the addition of 50 μL of ice-cold Buffer C1. The contents were mixed through gentle inversion back and forth 5 times, and then incubated on ice for 10 minutes. 250 μL of QIAseq Beads was added, mixed through inversion, and incubated at room temperature for 10 minutes, to capture the liberated nuclei. The tube was placed in a magnetic rack and incubated for 1 minute, after which the supernatant was carefully removed without disturbing the QIAseq Beads bound to the wall of the tube. The tube was then removed from the rack. Washing the beads was achieved by adding 500 μL of cold RNase-free water and gently inverting. The tube was again placed in a magnetic rack and incubated for 1 minute. After carefully removing the supernatant without disturbing the QIAseq Beads, proceeded immediately to the "Hi-C digestion" step.

###### ii. Hi-C digestion

The crosslinked chromatin was digested at GATC sites using a specific combination of buffers and enzymes in the Hi-C digestion solution. The Hi-C digestion solution for each sample was prepared by mixing Hi-C Digestion Buffer (4 μL), 1% SDS (4 μL), and RNase-Free Water (32 μL). Subsequently, 40 μL of this Hi-C Digestion Solution was added to the washed QIAseq Beads and mixed by gently pipetting 3-4 times to ensure complete resuspension. The tube was then incubated at 65°C for 10 minutes, followed by immediate placement on ice. After incubation, 4.4 μL of 10% Triton X-100 was added, and the mixture was mixed by gently pipetting up and down 3–4 times. Next, 4 μL of Hi-C Digestion Enzyme was added, and the mixture was again mixed through pipetting as before. To digest chromatin, the tube was incubated in a thermal mixer set at 37°C with 600 rpm shaking for 30 minutes. After the digestion step, the tube was incubated at 65°C for 20 minutes, followed by placement on ice to proceed to "Hi-C end labeling".

###### iii. Hi-C end labeling

While on ice, 6 μL of Hi-C End Labeling Mix and 1 μL of Hi-C End Labeling Enzyme were added to the tube containing the digested chromatin, from the last step. The contents were mixed by gently pipetting 3–4 times and then incubated at 37°C for 30 minutes to facilitate the labeling of digested chromatin with biotinylated nucleotides. Following incubation, the tube was placed on ice, ready to proceed to the next step, "Hi-C ligation".

###### iv. Hi-C ligation

The Hi-C Ligation Solution for each sample was prepared by combining Hi-C Ligation Buffer (200 μL), 10% Triton X-100 (40 μL), Ultralow Input Ligase (5 μL), and RNase-Free Water (105 μL). The full Hi-C Ligation Solution was then transferred to a tube with end-labeled chromatin. After transfer, the contents were thoroughly mixed by gently inverting the tube 5 times. The tube was incubated at 16°C for 30 minutes to facilitate the ligation, and then placed on ice, ready to proceed to "Chromatin de-crosslinking".

###### v. Chromatin de-crosslinking

To the ligated chromatin, 20 μL of Proteinase K solution was added, and the tube was gently inverted 5 times to mix thoroughly. The tube was incubated at 56°C for 30 minutes, followed by a subsequent incubation at 80°C for 90 minutes to ensure complete de-crosslinking. After the incubation, the tubes were briefly spun at low speed (4000 x g) to collect any condensation from the lid. Once cooled to room temperature, the reaction mixtures were ready for the next step, "DNA purification following de-crosslinking."

###### vi. DNA purification following de-crosslinking

After de-crosslinking, 40 μL of 3 M sodium acetate, pH 5.2, was added to the mixture, and the tube was briefly vortexed to ensure thorough mixing. Subsequently, 280 μL of 100% isopropanol was added to the tube, followed by another brief vortexing. A MinElute column, along with its provided collection tube, was placed in a rack. To bind the DNA, the entire mixture, including the QIAseq Beads, was applied to the MinElute column, which was then centrifuged for 1 minute at 17,900 × g. Flow-through was discarded, and the MinElute column was returned to the same tube. For washing, 0.75 mL of Buffer PE was added to the MinElute column then centrifuged for 1 minute at 17,900 × g. After discarding the flow-through and returning the MinElute column to the collection tube, an additional centrifugation step was performed for 1 minute at 17,900 × g. The MinElute column was then transferred to a new 1.5 mL microcentrifuge tube. For DNA elution, 35 μL of Buffer EB prewarmed to 65°C was added to the center of the membrane, followed by incubation for 1 minute at room temperature. Finally, the column was centrifuged for 1 minute at 17,900 × g to collect eluted DNA.

##### c. Processing Hi-C Library for NGS

###### i. Mechanical DNA fragmentation

The purified DNA underwent fragmentation in 100 μl of Buffer EB to achieve a median DNA fragment size of 400–600 bp, carried out through the Bioruptor® Pico sonication device, using 10 cycles with settings of 30 seconds on and 30 seconds off.

###### ii. DNA purification

Following fragmentation of the DNA sample, 4 volumes of Buffer SB1 were added to 1 volume of the fragmented DNA sample, and the mixture was briefly vortexed to ensure thorough mixing. A MinElute column, along with its provided collection tube, was then placed in a suitable rack. For DNA binding, the mixture was applied to the column, centrifuged for 1 minute at 17,900 x g, flow-through discarded, and the spin column returned to the tube. For washing, 700 μL of 80% ethanol was added to the MinElute column, followed by centrifugation for 1 minute at 17,900 x g. This washing step was repeated. After discarding flow-through and returning the MinElute spin column to the tube, it was centrifuged for an additional 1 minute at 17,900 x g. The MinElute spin column was then placed into a clean 1.5 mL microcentrifuge tube. Next, 50 μL of Buffer EB prewarmed to 65°C was added to the center of the membrane, and the column was allowed to stand for 1 minute before being centrifuged for 1 minute. Finally, the tubes were frozen overnight at −20°C for further processing.

###### iii. Streptavidin pulldown of Hi-C fragments

The Streptavidin Beads were thoroughly resuspended by vortexing. Subsequently, 25 μL of Streptavidin Beads was transferred into a new microcentrifuge tube, then placed in a magnetic rack and incubated for 1 minute. After removing supernatant, the tube was taken off the rack, 100 μL of Bead Wash Buffer 1 was added, and vortexed for 10 seconds. To capture the beads, the tube was placed in a magnetic rack and incubated for 1 minute before removing the supernatant. Beads were resuspended in 50 μL of Bead Resuspension Buffer. Subsequently, 50 μL of purified DNA sample was added to the beads, and the mixture was incubated at room temperature for 15 minutes in a thermal mixer set at 1000 rpm. Following incubation, the protocol proceeded directly to the next step.

###### iv. DNA end-repair/A-tailing/phosphorylation

The tube was placed in a magnetic rack and incubated for 1 minute before removing the supernatant and taking the tube off the rack. Next, 100 μL of Bead Wash Buffer 2 was added to the tube and vortexed for 10 seconds. The tube was placed in a magnetic rack to capture the beads, followed by incubation for 1 minute before removing supernatant. For each sample, ER/A-tailing solution was prepared using 5 μL of ER/A-Tailing Buffer, 10 μL of ER/A-Tailing Enzyme Mix, and 35 μL of RNase-Free Water. The beads were resuspended in 50 μL of the prepared ER/A-tailing solution. Subsequently, the tube was incubated for 15 minutes at 20°C, followed by an additional 15 minutes at 65°C.

###### v. Adapter ligation (Illumina)

The tube was placed in a magnetic rack and incubated for 1 minute before removing supernatant and taking the tube off the rack. Next, 100 μL of Bead Wash Buffer 2 was added to the tube and vortexed for 10 seconds. The tube was placed in a magnetic rack to capture the beads, followed by incubation for 1 minute before removing supernatant. Adapter ligation buffer dilution was prepared using 15 μL of Adapter ligation buffer and 135 μL of RNase-Free water for each sample. 95 μL of the prepared diluted Adapter Ligation Buffer was added to each tube then vortexed for 10 seconds. The tube was placed in a magnetic rack to capture the beads, incubated for 1 minute, and then supernatant was removed. The beads were resuspended in 50 μL of diluted Adapter Ligation Buffer. Subsequently, 5 μL of Illumina Adapter was transferred to the tube, and 2 μL of Ultralow Input Ligase was added to the sample. The mixture was carefully pipetted up and down 3–4 times and incubated for 45 minutes at room temperature.

###### vi. Amplification of Hi-C sequencing library

The tube was placed in a magnetic rack and incubated for 1 minute before removing supernatant and taking the tube off the rack. Next, 100 μL of Bead Wash Buffer 1 was added to the tube and vortexed for 10 seconds. The tube was then placed in a magnetic rack to capture the beads, incubated for 1 minute, and supernatant removed. Wash step was repeated. After removing the supernatant and taking the tube off the rack, 100 μL of Bead Wash Buffer 2 was added to the tube, and the process was repeated as before. Following this, the tube was placed in a magnetic rack, incubated for 1 minute, and the supernatant removed. After repeating the wash step once more, the tube was placed in a magnetic rack, incubated for 1 minute, and the supernatant was removed. Finally, 100 μL of RNase-Free Water was added to the tube and the tube was vortexed for 10 seconds. The tube was placed in a magnetic rack to capture the beads, incubated for 1 minute, and the supernatant was removed. For Hi-C sequencing library amplification, 75 μL of HiFi PCR Master Mix, 2x, 4.5 μL of Primer Mix Illumina Library Amp, and 70.5 μL of RNase-Free water were combined for each sample. Then, 150 μL of the prepared Hi-C sequencing library amplification mix was added to the beads, vortexed briefly for complete resuspension, and pipetted into a single PCR tube, a single well of an 8-well PCR strip, or 1 position of a 96-well PCR plate. The PCR tubes were transferred into a thermocycler according to the program described in the table below.

| Time | Temperature | Number of cycles |
| --- | --- | --- |
| 2 min | 98°C | 1 |
| 20 s | 98°C | 12 (>5x10^4 human cells/sample; >300ng DNA/sample)  16 (<5x10^4 human cells/sample; <300ng DNA/sample) |
| 30 s | 60°C |  |
| 30 s | 72°C |  |
| 1 min | 72°C | 1 |
| ∞ | 4°C | Hold |

PCR reaction was transferred to a fresh 1.5ml low bind tube and the libraries were placed in a magnetic rack and incubated for 1 minute to separate the libraries from the streptavidin beads. The clear supernatant was transferred into a new microcentrifuge tube.

###### vii. Purifying Hi-C sequencing library

The QIAseq Beads were briefly vortexed to thoroughly resuspend them before adding 150 μL to the clear supernatant (contain amplified NGS library). After brief vortexing, the mixture was incubated for 5 minutes at room temperature. Subsequently, the tube was placed in a magnetic rack and incubated for 1 minute, followed by removal of supernatant and taking the tube off the rack. Next, 500 μL of 80% ethanol was added to the tube and vortexed for 10 seconds. After placing the tube in a magnetic rack to capture the beads and incubating for 1 minute, the supernatant was removed. This wash step was repeated once more, to ensure maximum removal of ethanol. The tube was briefly centrifuged at 5000 x g, room temperature, transferred back to the magnetic rack, and incubated for 30 seconds. Maximum volume of ethanol was removed from bottom of the tube before incubation with the lid open in the magnetic rack for an additional 2–5 minutes or until the beads were dry. After removing the tube from the magnetic stand, the DNA library was eluted by adding 25 μL of Buffer EB to the beads. The mixture was pipetted up and down until the beads were completely resuspended and then incubated for 1 minute at room temperature. The tube was placed in a magnetic rack and incubated for 1 minute more, after which the supernatant containing the purified Hi-C sequencing library was transferred into a new DNA low-binding tube.

###### viii. Hi-C library quality control and quantification

Quality of the Hi-C sequencing libraries were determined using D5000 ScreenTape and Qubit™ 1X dsDNA High Sensitivity (HS) assay. The purified library was stored at –20°C in a DNA low-binding tube until ready to use for sequencing.

##### d. Sequencing on Novaseq 6000

Each library was normalized to 3nM in 35 μL. Samples with 6 different Illumina adapters were pooled into one pool, resulting in a total volume of 210 μL. The pool was thoroughly vortexed and spun down. Subsequently, 150 μL of this mix was transferred to a new 1.5 mL tube, and 37 μL of 0.2N HP3 was added. After vortexing and spinning, the mixture was incubated at room temperature for 8 minutes. Following this, 38 μL of 400 mM Tris-HCl, pH 8.0, was added, and the mixture was vortexed and spun down again. Finally, 225 μL of the mix was added to the library tube and loaded onto the Novaseq 6000 for sequencing.

#### E. Exome sequencing

Genome purification was performed using QIASymphony extraction Midi Kit (Qiagen). Ultrasonication (Covaris, USA) was used to fragment the DNA, followed by coding regions (exome) capturing using the Agilent Clinical Research Exome V2 (CREv2) capture probes (Agilent, USA). Library preparation was done using SureSelectXT protocol (Agilent, USA) followed by sequencing using the NovaSeq system (Illumina, USA); sequencing was performed at (2 x 150bp) with a minimum average depth of 100x*9*.

#### F. Variant identification and filtering

We reported the number of SNPs, indels, and SVs, the ratio of base substitutions, and the indel length distribution for each sample. For SNPs, variants detected in our samples were compared against dbSNP^1^ (v151), gnomAD^2^ (v2.2.1), the 1000 Genomes Project^3^ (phase1_v3.20101123), and GME^4^ (downloaded from ANNOVAR). For indel, variants were compared against dbSNP (v151), gnomAD (v2.2.1), the 1000 Genomes Project (Mills_and_1000G_gold_standard.indels.hg38), and GME (downloaded from ANNOVAR). Structural variants were compared against the DGV^5^ Gold Standard Variants and the 1000 Genomes Project data (phase1_v3.20101123) with 80% reciprocal overlap. We used custom perl script for the analysis.

For mitochondrial haplogroup prediction, the command employed was:

*haplogrep classify –in APR_MT.vcf –format vcf –out haplogroups.txt –lineage 2*

For Y haplogroup prediction, command used was:

*LineageTracker classify --vcf APR_Y.vcf -b 38 -o Y-Haplo_Result*

#### G. *de novo* genome assembly

High-quality *de novo* assemblies were generated for each sample using a hybrid assembly approach, combining PacBio HiFi and ONT ultralong reads. For each of the samples we ran HiFiAdapterFilt on the ubams to clean up any reads with adapters still attached. The resulting fastq files were combined into one file per sample using the zcat utility. We also combined all of the ultra-long DNA sequencing kit (ULK) reads (in fastq format) into one fastq file before using it along with the HiFi reads to build assemblies. Further, Hifiasm^6^ v0.19.5-r603 was used with the following arguments --ul APR029.ul.fastq and APR029.hifi.fastq.

For the trio, Verkko^7^ and Hifiasm was run to produce completely phased assemblies. To run Verkko, kmer databases were built using Meryl with the following command using the paternal HiFi reads

*meryl count compress k=30 threads=XX memory=YY maternal.*fastq.gz output maternal_compress.k30.meryl*

*meryl count compress k=30 threads=XX memory=YY paternal.*fastq.gz output paternal_compress.k30.meryl*

*$MERQURY/trio/hapmers.sh maternal_compress.k30.meryl paternal_compress.k30.meryl*

Verkko pipeline was run using the following command:

*verkko -d asm --hifi son.hifi.fastq --nano son.ul.fastq --hap-kmers maternal_compress.k30.hapmer.meryl paternal_compress.k30.hapmer.meryl trio*

To run Hifiasm, parental kmer databases were built using the following commands:

*yak count -k31 -b37 -t16 -o pat.yak paternal.fq.gz*

*yak count -k31 -b37 -t16 -o mat.yak maternal.fq.gz*

Hifiasm pipeline was run using the following sample command:

*hifiasm -o apr_son.asm -t 32 -1 pat.yak -2 mat.yak --ul son.ul.fastq son.hifi.fastq*

### Supplementary Figures

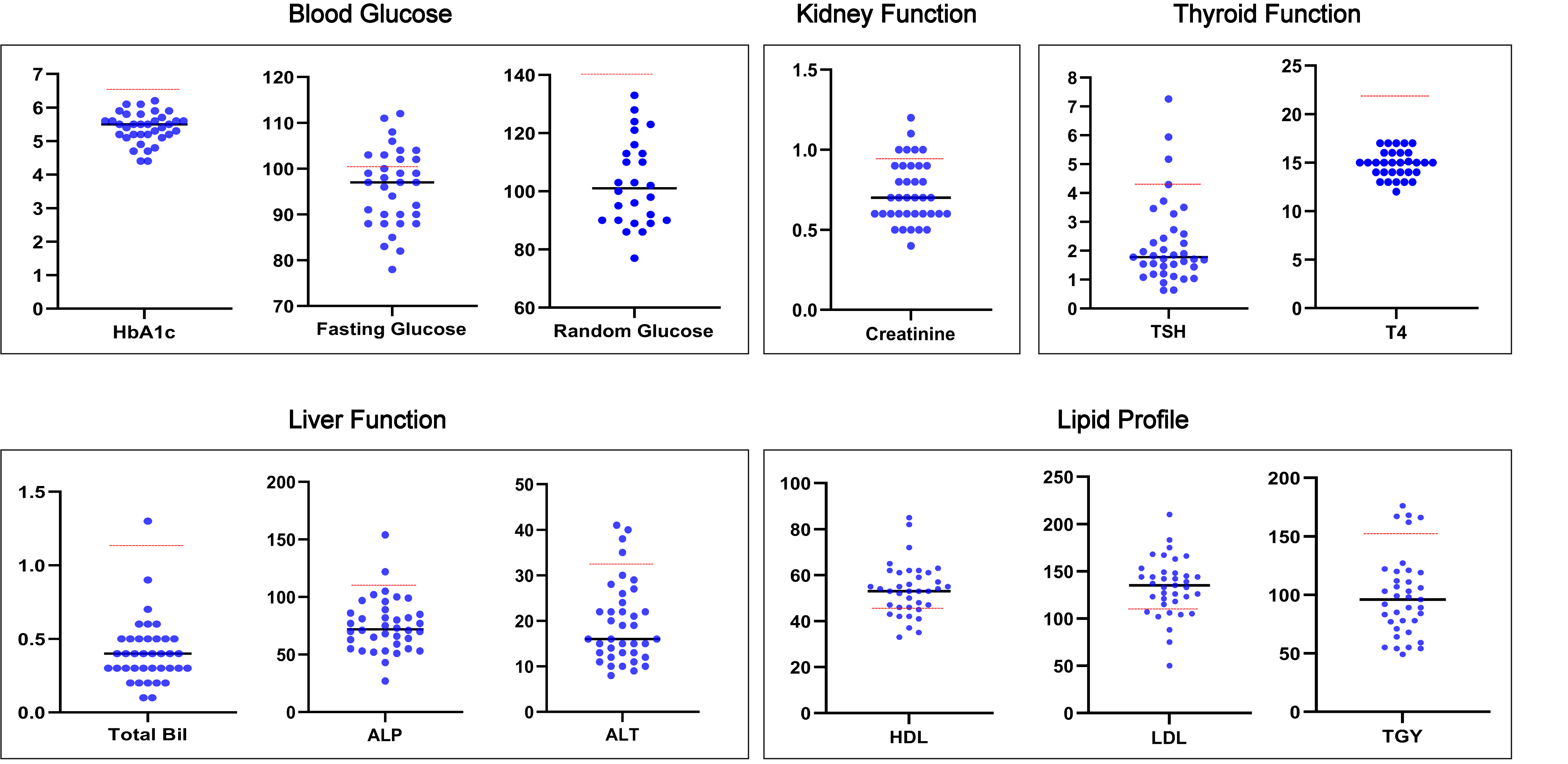

#### Supplementary Figure 1: Key serum measures kidney, thyroid, and liver function in addition to glucose and lipid values.

Each plot showcases a specific phenotypic attribute, with dots representing individual measurements and the red line representing the normal threshold.

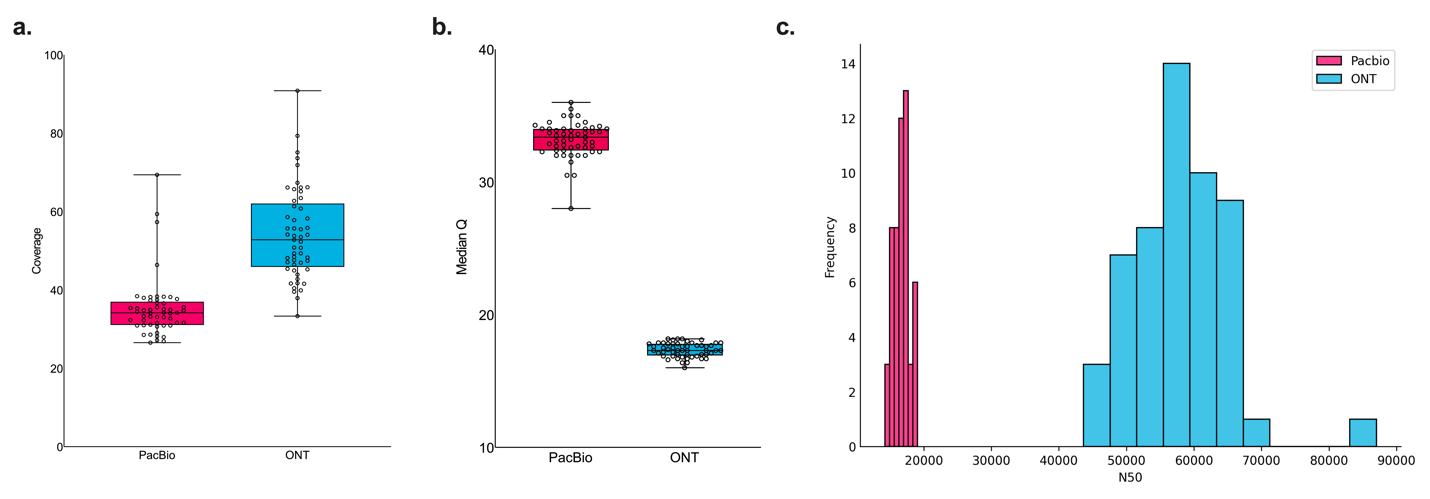

#### Supplementary Figure 2: Comparison of sequencing metrics for PacBio and ONT sequencing.

**a.** Box plot displaying genome coverage comparison between PacBio (in magenta) and ONT (in cyan). **b.** Box plot of median Q scores between the two sequencing methods. The y-axis represents Q score values, indicating the quality of sequencing reads. **c.** Histogram presenting the distribution of N50 read lengths.

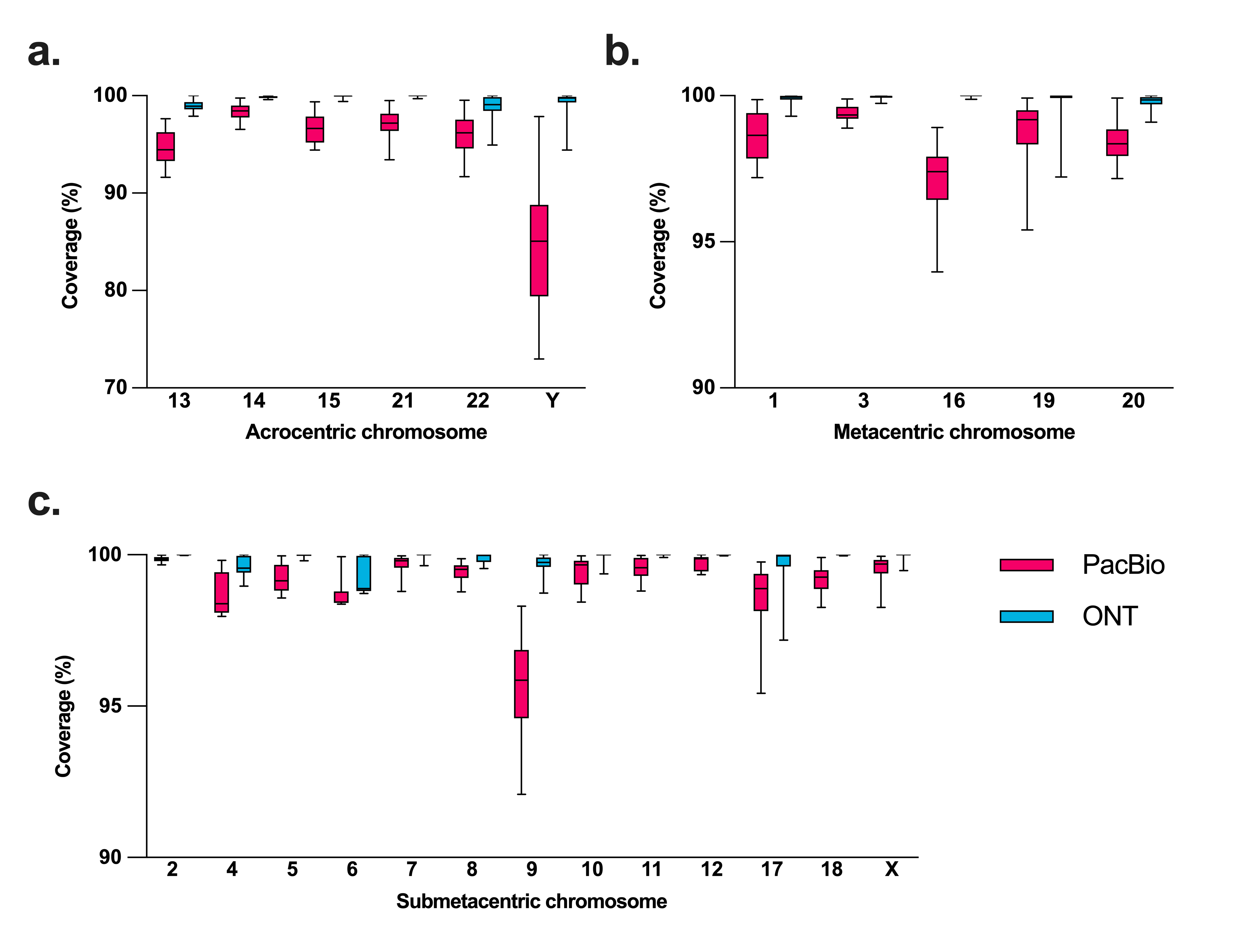

#### Supplementary Figure 3: Coverage of chromosomes from APR cohort.

Sample wise alignment to CHM13 across each chromosome comparing Hifi (magenta) and ONT (cyan) reads. The box plot represents the distribution of coverage from all samples for **a.** acrocentric, **b.** metacentric and **c.** submetacentric chromosomes.

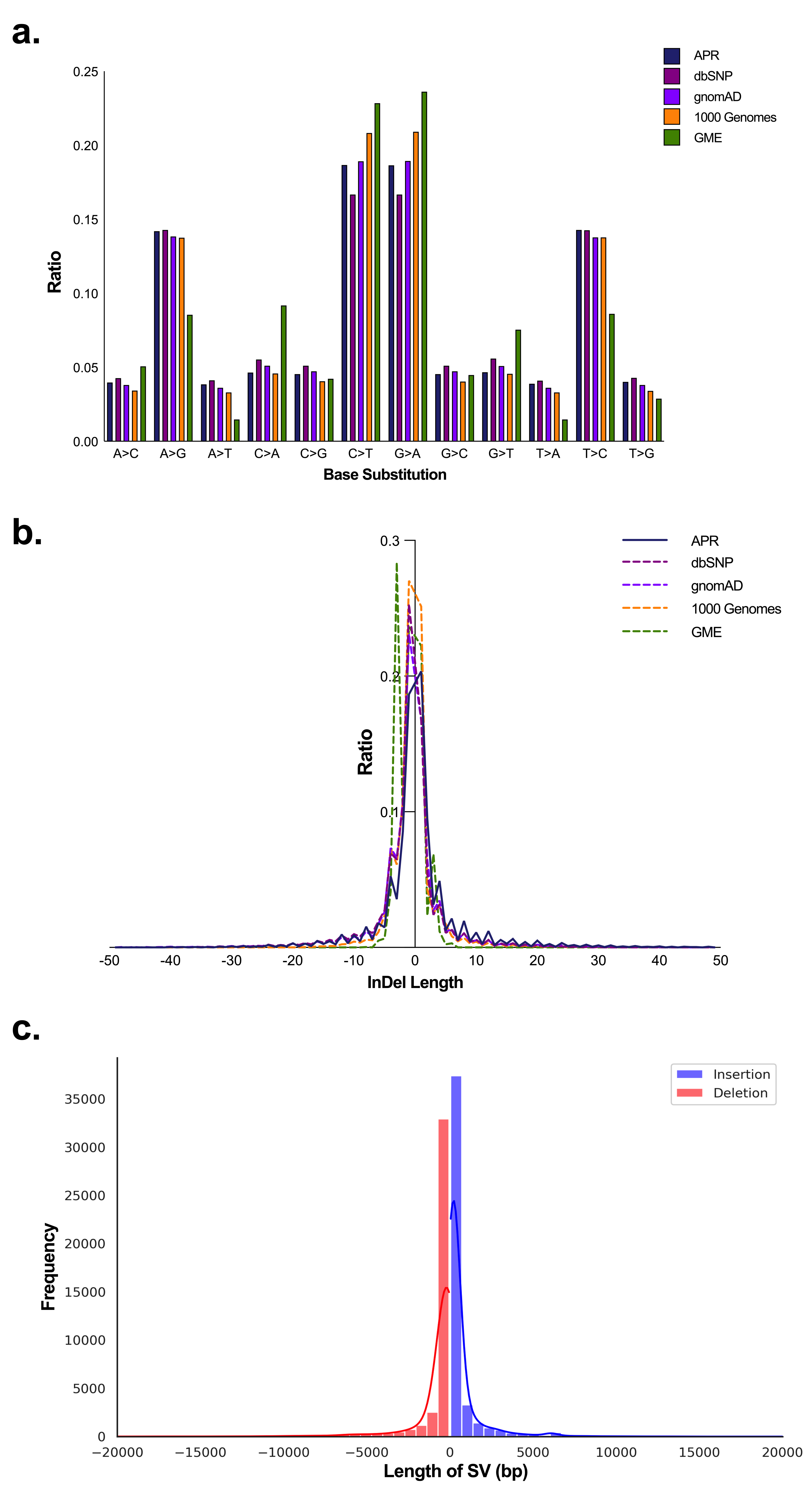

#### Supplementary Figure 4: Comparative variant analysis of databases used in identification of novel population-specific variants.

**a.** Bar chart comparing the ratio of different base substitutions across multiple datasets: APR, dbSNP, 1000 Genomes, and GME. The x-axis lists base substitution types, while the y-axis represents their ratios within each dataset. **b.** Line graph depicting the ratio of indel lengths across datasets (APR, dbSNP, 1000 Genomes, and GME). The x-axis shows indel length in base pairs (bps), while the y-axis shows the ratio of indels present at different lengths. **c.** Frequency distribution of the length of structural variations (SVs), classified as insertions (blue) or deletions (red). The x-axis denotes the length of SVs in base pairs, and the y-axis shows the frequency.

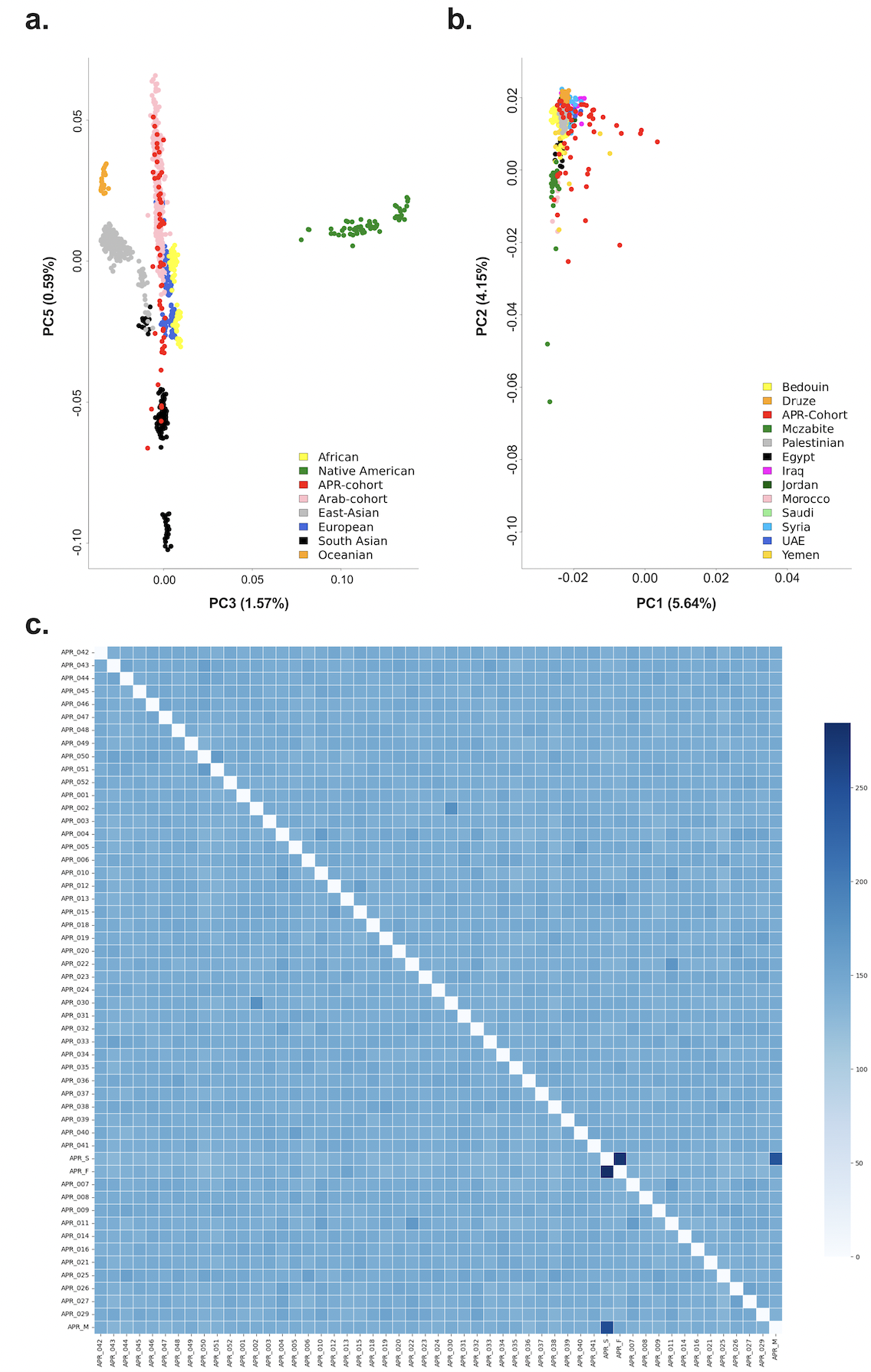

#### Supplementary Figure 5: Principal Component and fineSTRUCTURE analysis of APR cohort and Arab cohort.

**a.** PCA clustering of APR cohort in association with Arab ethnicities. Scatter plot displays the PCA clustering of APR cohort alongside Arab samples. The distribution is plotted using Principal Component 5 (PC5) on the y-axis against Principal Component 3 (PC3) on the x-axis, highlighting inherent clustering between these groups. The analysis includes samples from Human Genome Diversity Project (HGDP) and Human Origins database, with ach ethnicity color coded for distinction. APR cohort is highlighted in red. **b.** PCA scatter plot depicting population variance of APR samples across Arab ethnic subpopulation. The two axes indicate the variance captured by first (PCA1) and second (PCA2) principal component, respectively. **c.** The fineSTRUCTURE haplotype sharing information matrix shows relatedness between APR samples and the trio (dark blue square).

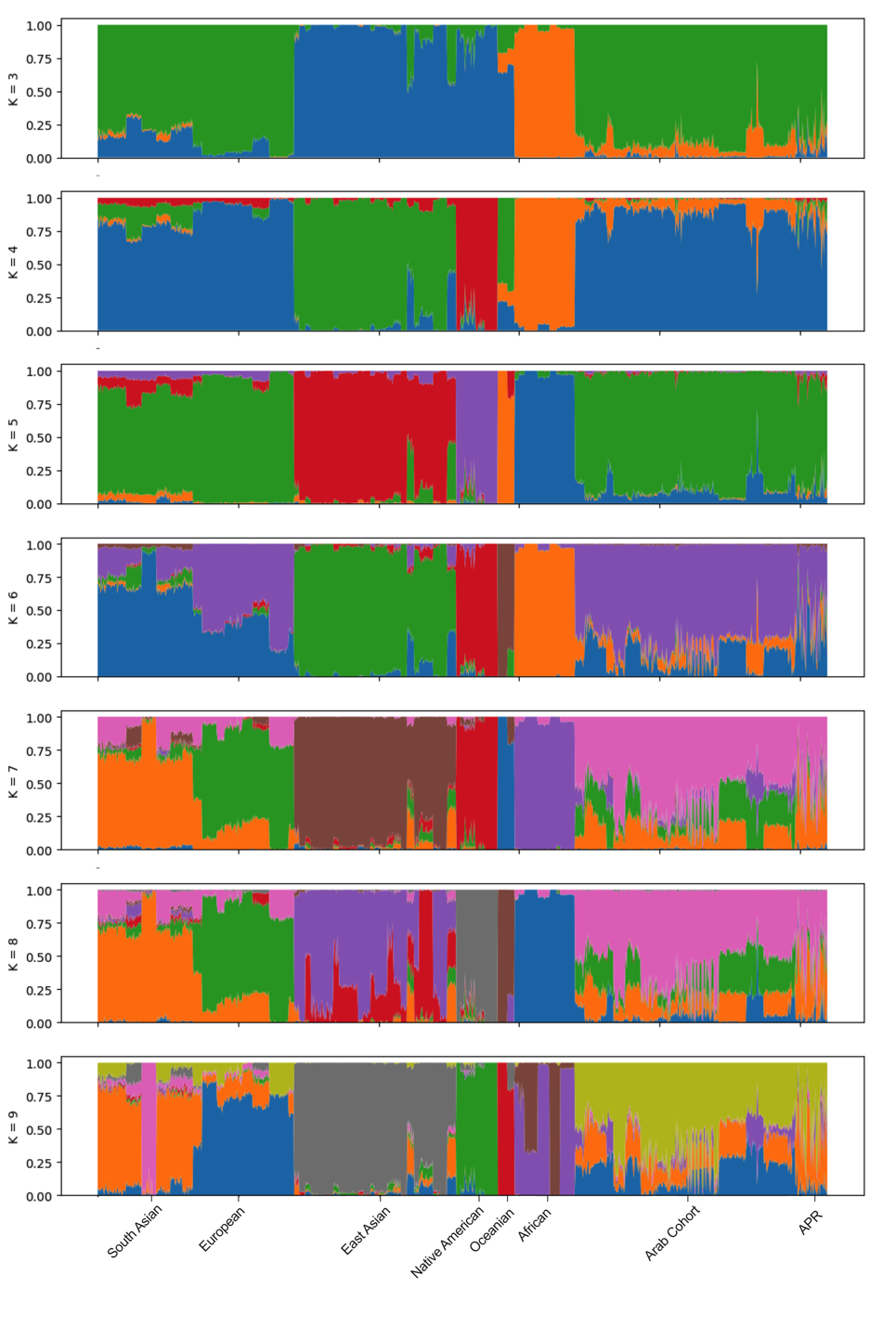

#### Supplementary Figure 6: Population genetic ancestry inference of the 53 APR samples using ADMIXTURE.

Assuming ancestry components K ranging from 3 to 9, each plot shows independent ancestry fraction with its own color-coding representation labeled with short vertical lines. Samples included are from South Asian, European, East Asian, Native American, Oceanian, African and Arab ethnicities.

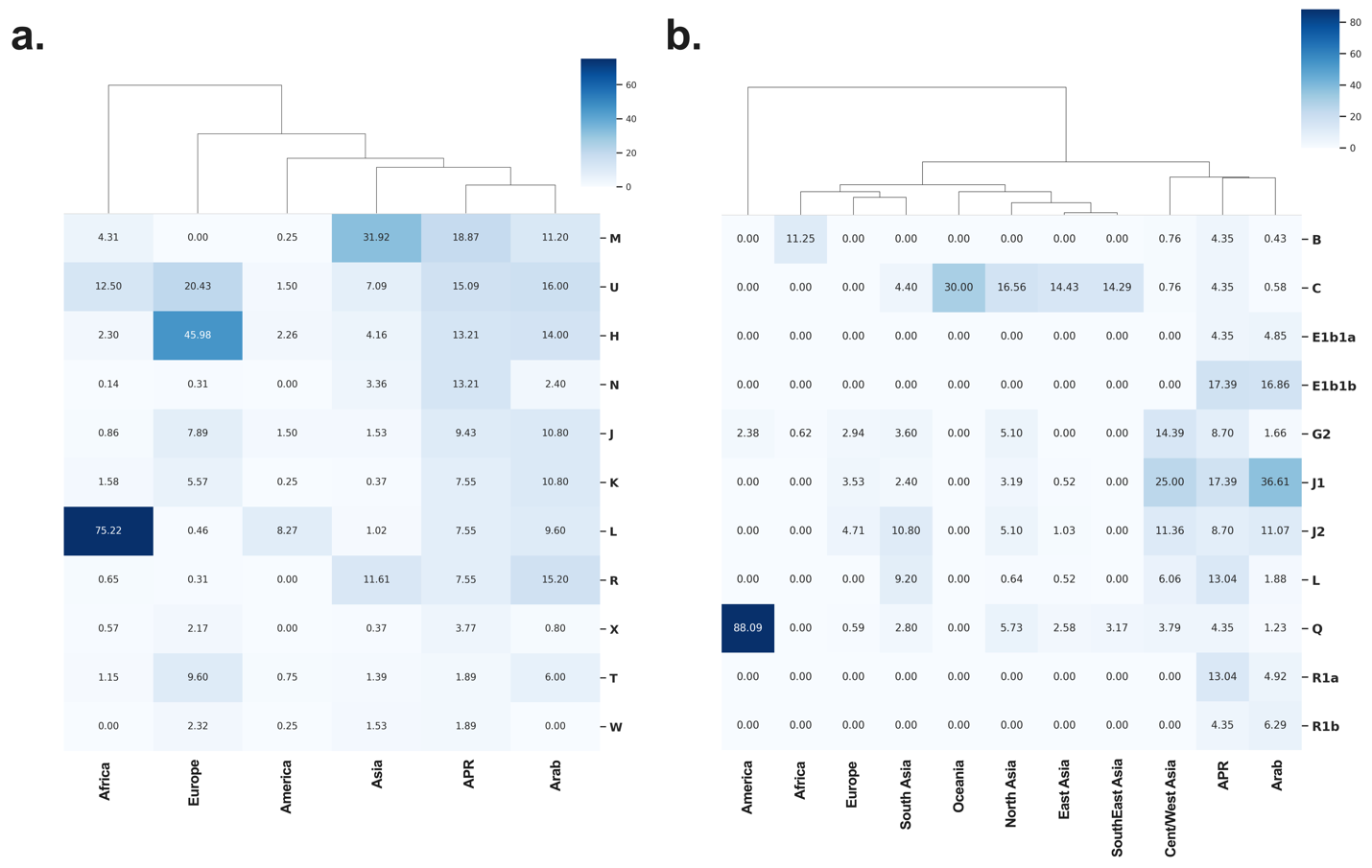

#### Supplementary Figure 7: Y and MT haplogroup distribution.

**a.** Hierarchical clustered heatmap displaying the distribution of mitochondrial haplogroup (M, U, H, N, J, K, L, R, X, T and W) frequencies across the APR cohort and reference populations (Africa, Europe, America, Asia and Arab). **b.** Hierarchical clustered heatmap showcasing the Y chromosome haplogroup (B, C, E1b1a, E1b1b, G2, J1, J2, L, Q, R1a and R1b) frequencies among the APR cohort compared to reference populations (America, Africa, Europe, South Asia, Oceania, North Asia, East Asia, Southeast Asia, Central Asia and Arab).

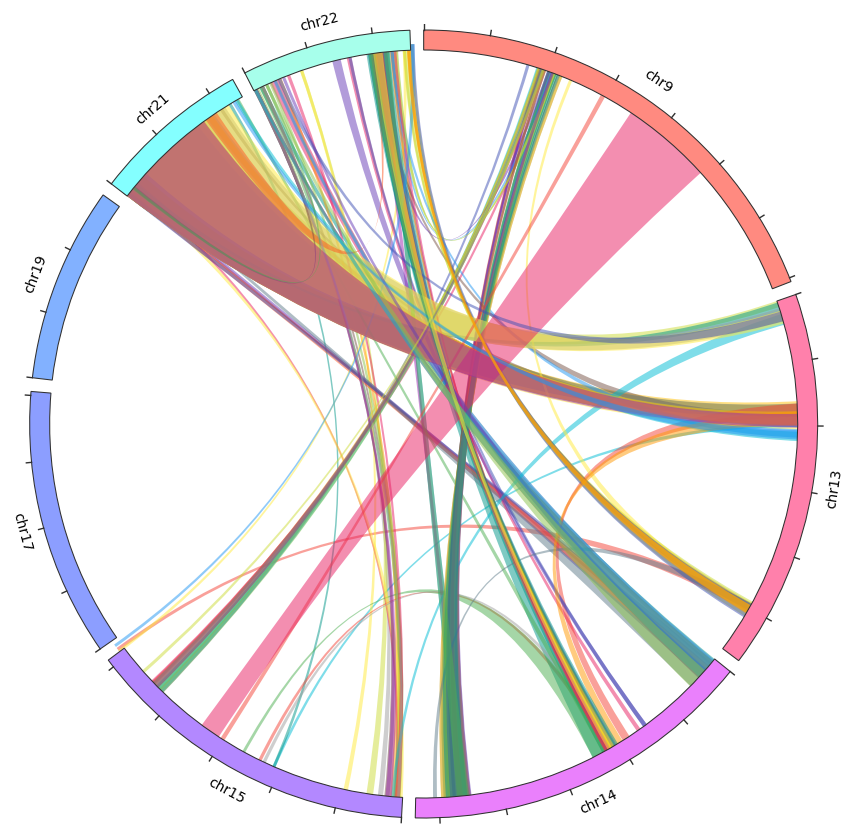

#### Supplementary Figure 8: Visualization of inter-chromosomal mis-joins in APR assemblies.

The circular plot represents different chromosomes labeled as ‘chr’ followed by their respective numbers. Each connecting line signifies an inter-chromosomal mis-join, with the width of the line corresponding to the length of the mis-join. The most prominent mis-join, observed between chr21 and chr13, is also found in the HPRC dataset.

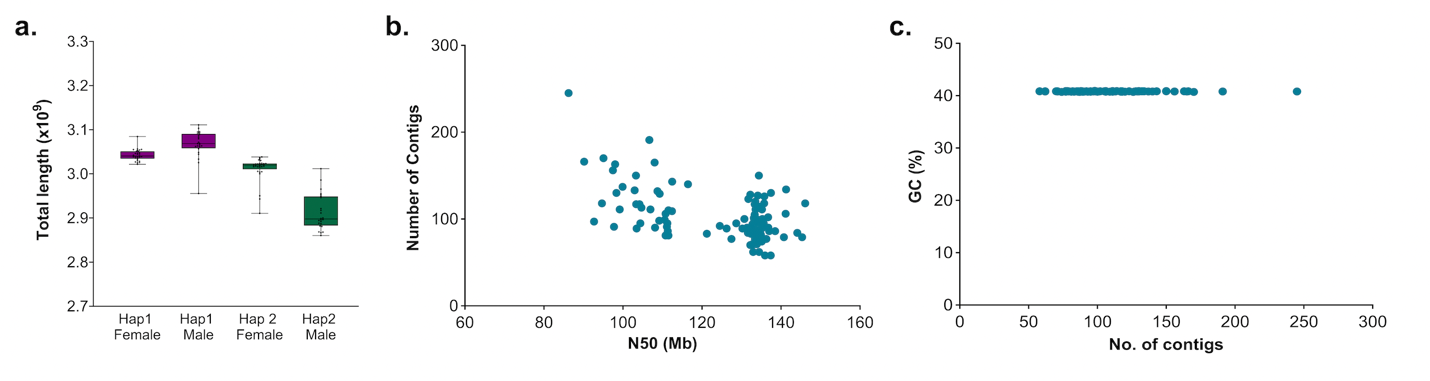

#### Supplementary Figure 9: APR assembly characteristics.

**a.** Box plot comparing total genome length for samples across two haplotypes (Hap1 and Hap2) separated by sex (female and male). The y-axis represents the total length in gigabases (x10^9^). **b.** Scatter plot depicting the relationship between the N50 length metric (x-axis, in megabases) and the total number of contigs (y-axis). Each blue dot represents an individual sample. **c.** Scatter plot illustrating the GC content percentage (y-axis) against the number of contigs (x-axis) for each sample.

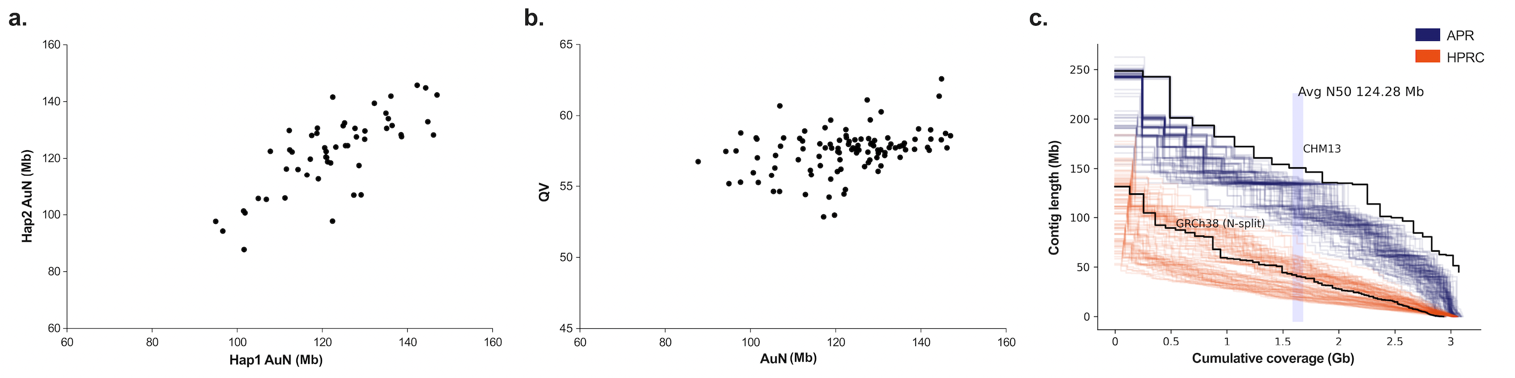

#### Supplementary Figure 10: Assembly contiguity and quality metrics.

**a.** Scatter plot presenting Assembly Unit Numbers (AUN) in Mb of Hap2 on y-axis against Hap1 on x-axis. **b.** Scatter plot showing relationship between QV on y-axis and AUN on x-axis. **c.** Line graph comparing contig length against cumulative assembly coverage. APR dataset is indicated in blue, and HPRC dataset in orange. Contiguity reference lines for the CHM13 and GRCh38 genomes are provided. The highlight at the average N50 value (124.28 Mb) offers a benchmark for contiguity within APR assemblies.

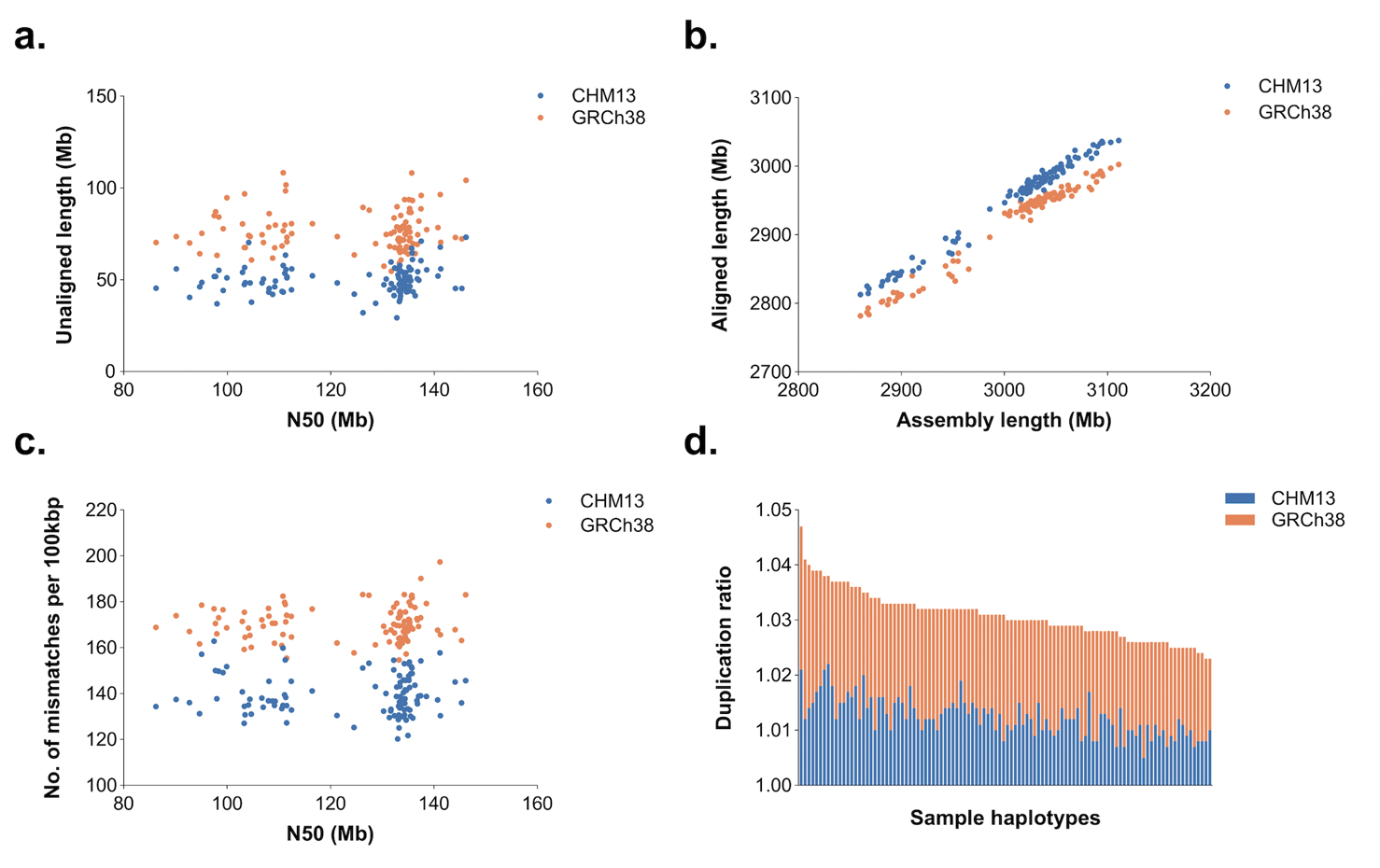

#### Supplementary Figure 11: Aligned and unaligned contigs.

**a.** The y- and x-axis refer to unaligned contig length in mega bases (Mb) and their contig N50 length, respectively, for two references. The data points depicted in orange represent GRCh38 and blue represent CHM13 genome reference. **b.** The aligned length (y-axis) and the assembly length (x-axis) for each APR sample. **c.** The mismatch in every 100,000 bases (y-axis) and the contig N50 length (x-axis). **d.** Bar graph displaying the distribution of duplication ratios across assemblies.

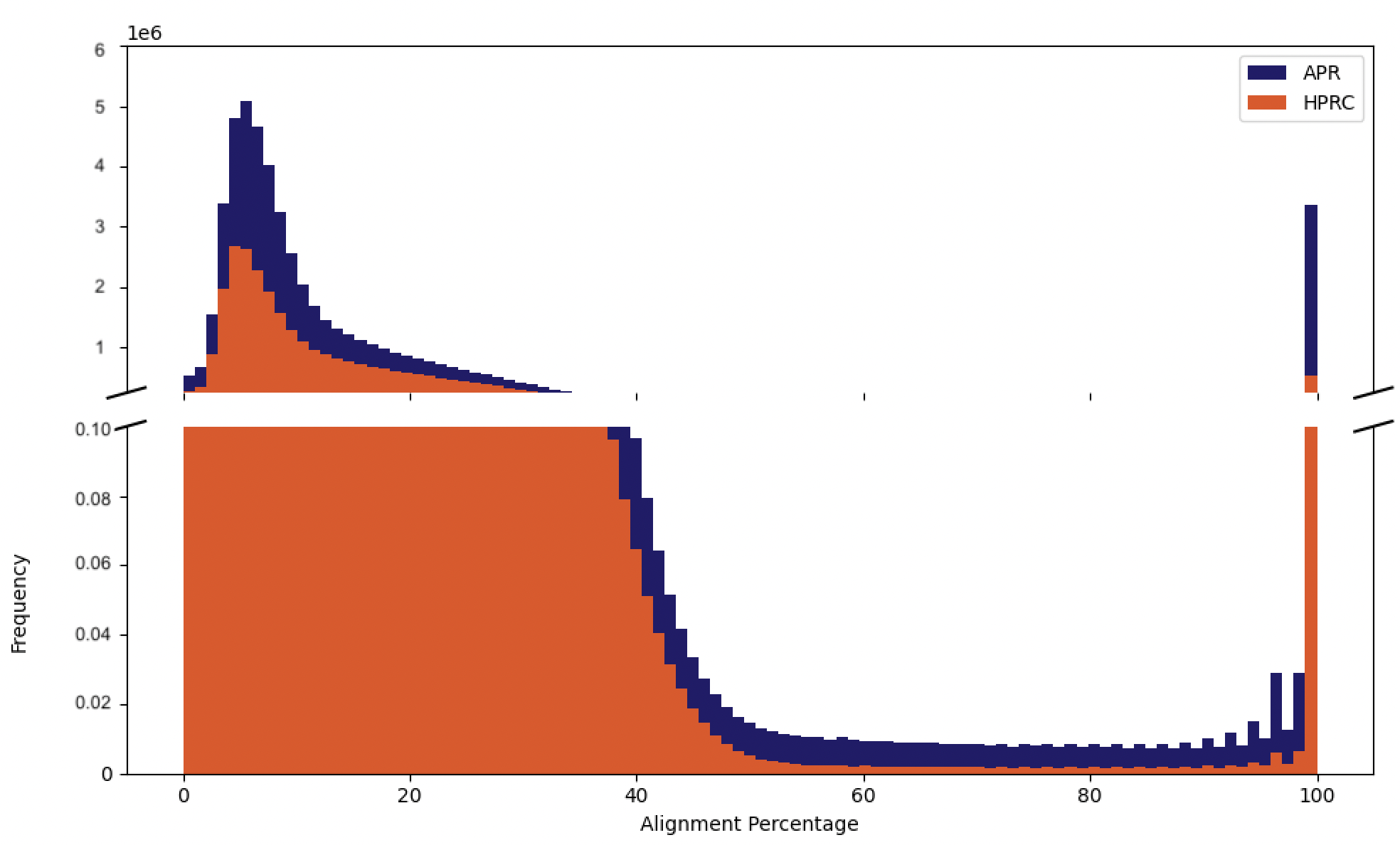

#### Supplementary Fig. 12: Distribution of alignment percentages in centromeric regions computed using UniAligner tool.

The histogram displays the frequency distribution of alignment percentages for 100bp windows across all centromeric regions, comparing the APR and HPRC datasets. The alignment percentage was computed using the UniAligner tool, illustrating the comparative performance of APR (blue) versus HPRC (orange).

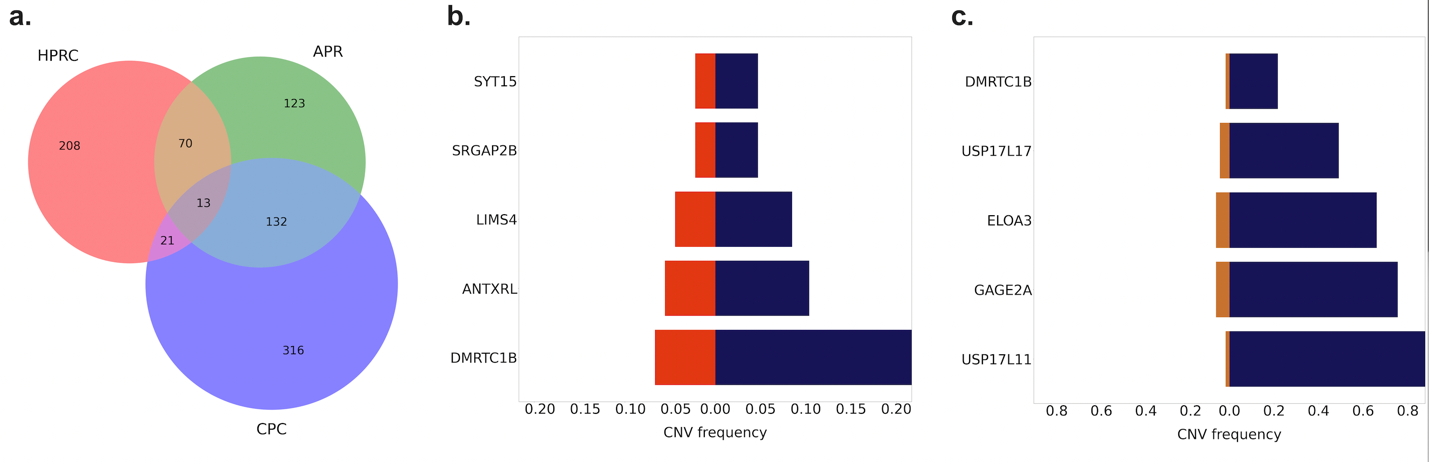

#### Supplementary Fig. 13: Analysis of recurrent gene duplication patterns.

**a.** Venn diagram representing the overlap of unique recurrent duplicated genes among APR, HPRC, and CPC assemblies. **b.** Comparative frequency of duplicated genes between APR and HPRC assemblies. The bar graph depicts five overlapped duplicated genes with a higher frequency (≥5%) in Arab assemblies (blue) compared to HPRC (orange). **c.** Comparative frequency of duplicated genes between APR and CPC assemblies. The bar chart illustrates five overlapped duplicated genes with a significantly higher frequency (≥5%) in Arab assemblies (blue) in contrast to CPC (orange).

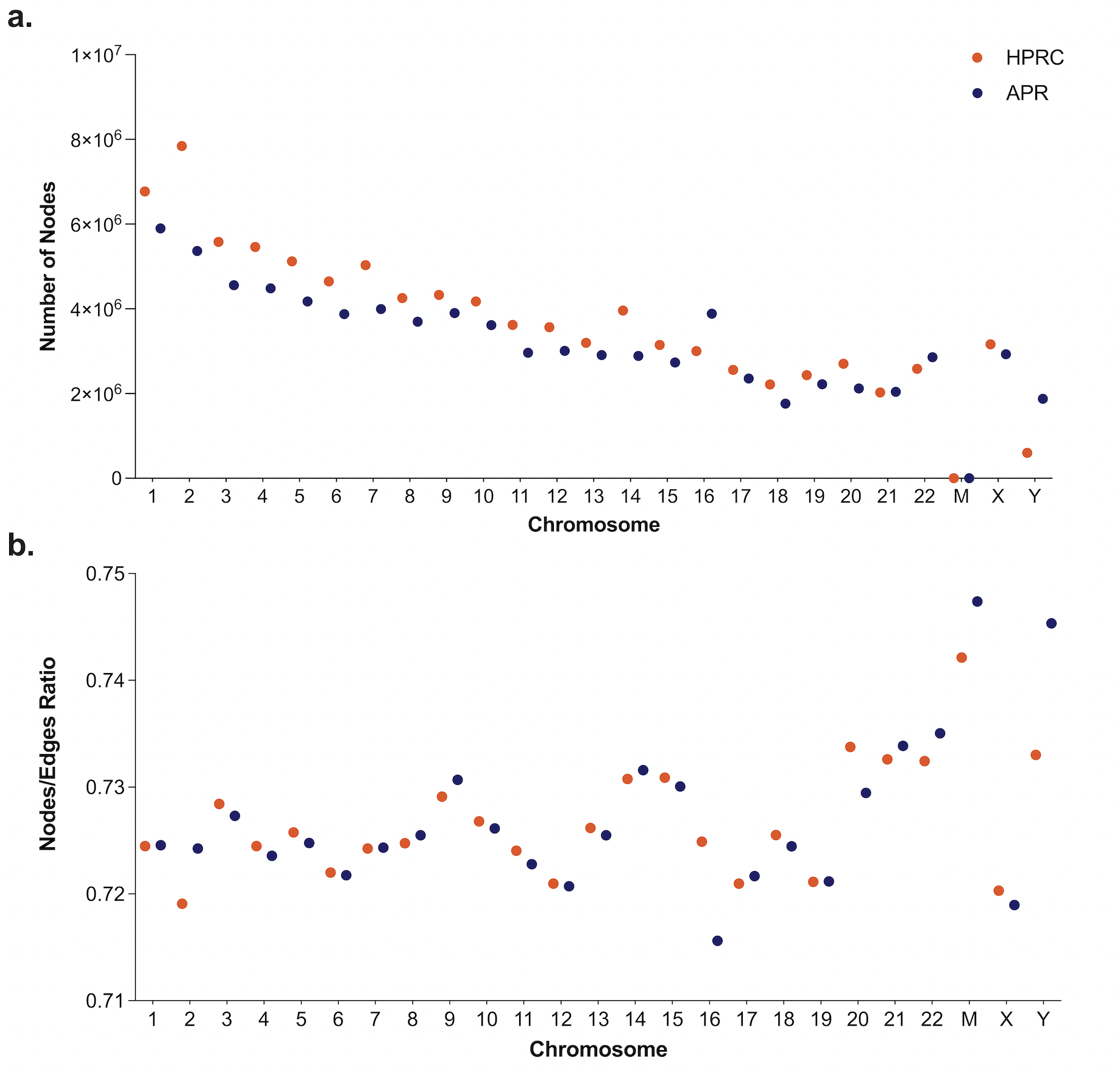

#### Supplementary Figure 14: Pangenome graph characteristics.

**a.** Each dot represents the number of nodes per chromosome. The blue and orange dot represents number of nodes in APR and HPRC pangenome graph, respectively. **b.** The Nodes/edges ratio comparison of HPRC and APR pangenome for each chromosome.

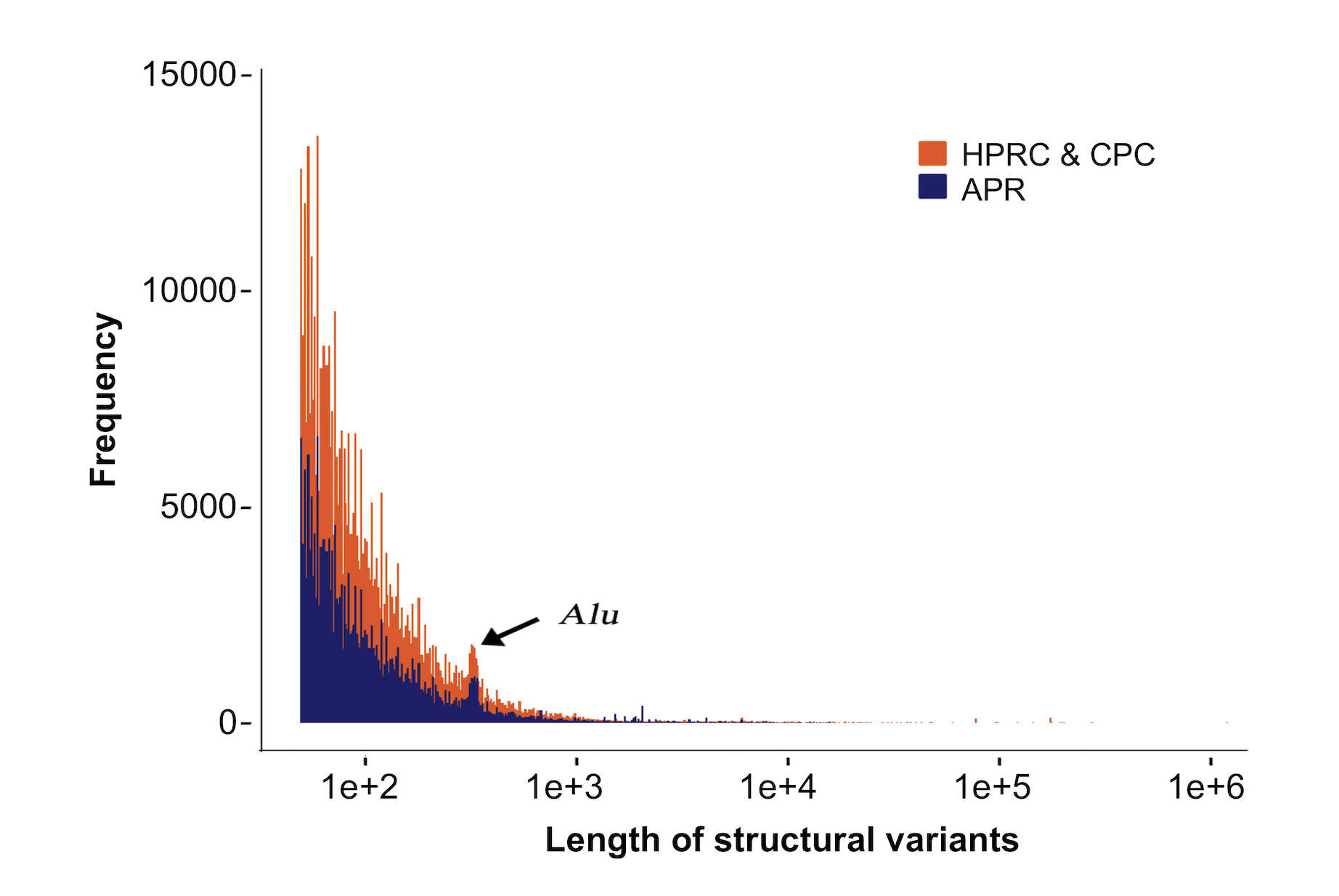

#### Supplementary Figure 15: Structural variants from pangenome graph.

SV length distribution for APR (blue) and HPRC+CPC (orange). The peak at 300 bp for Alu repeats are highlighted.

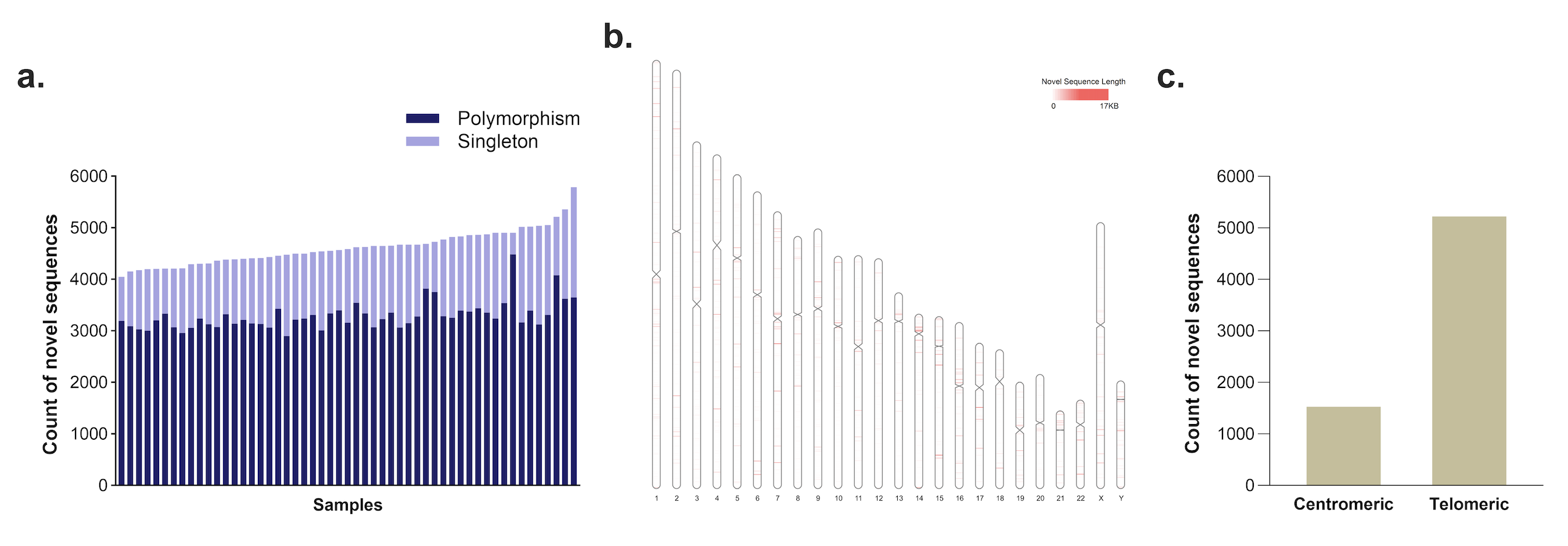

#### Supplementary Figure 16: Novel sequences identified in APR assemblies in comparison to HPRC and CPC, CHM13, GRCh38, 1000G and DGV.

**a.** Bar graph indicating the count of newly discovered sequences for each sample, differentiated between polymorphism (dark blue) and singleton (light blue). **b.** Loci of APR specific novel sequences. Visualization of Arab-specific novel sequences across chromosomes. The line (red) width is proportional to the length of the novel sequence (kb). **c.** Bar graph indicating count of novel sequences across centromeric and telomeric regions.

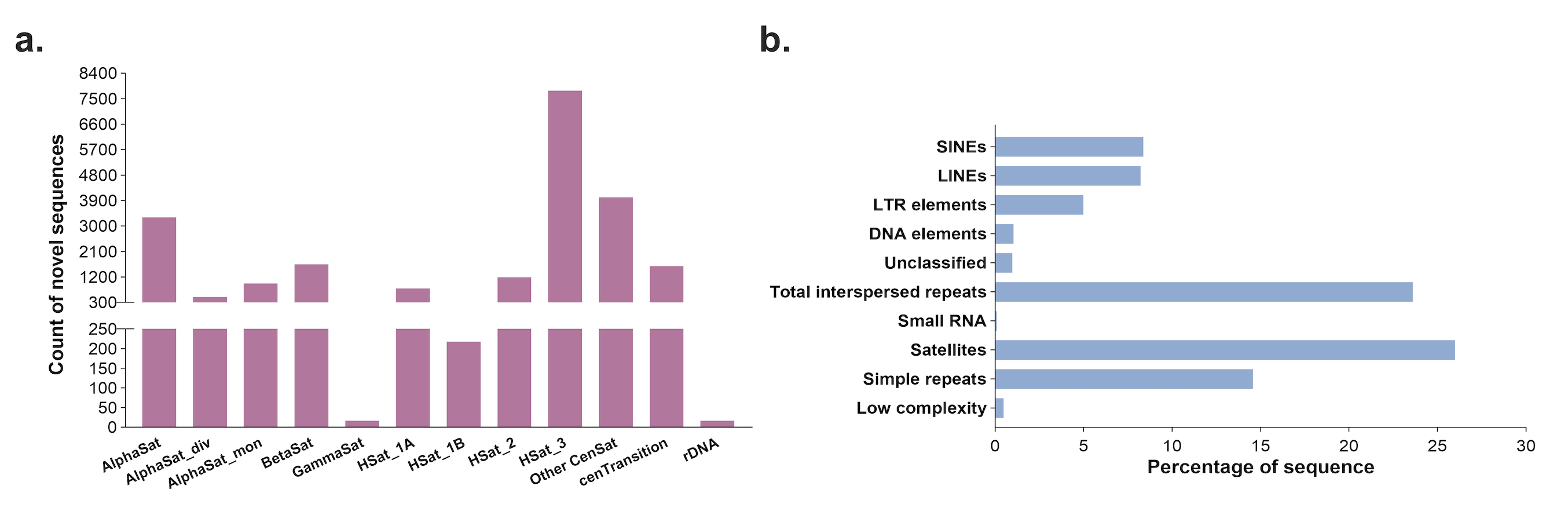

#### Supplementary Figure 17: APR Novel euchromatic Sequences.

**a.** Bar graph indicating counts of novel sequences across microsatellite regions. **b.** Satellite repeats in novel sequences identified using Repeatmasker.

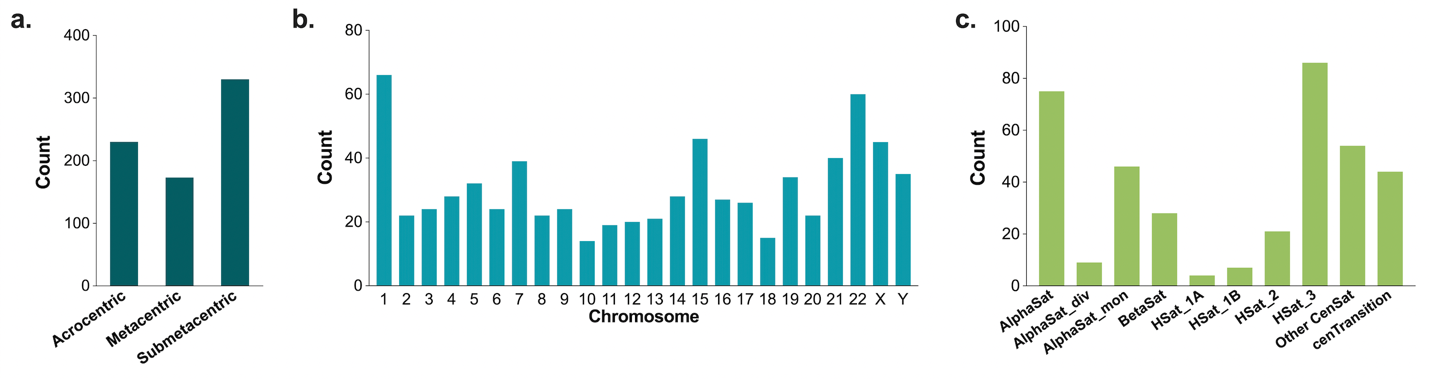

#### Supplementary Figure 18: Chromosome wise distribution of complex structural variations (CSVs) sites.

**a.** Bar graph indicating the count (y-axis) of CSV sites across chromosome types: acrocentric, metacentric, and submetacentric. **b.** Bar graph showing the count of CSV sites dispersed across all individual chromosomes. **c.** Bar graph indicating counts of CSV sites across microsatellite regions.

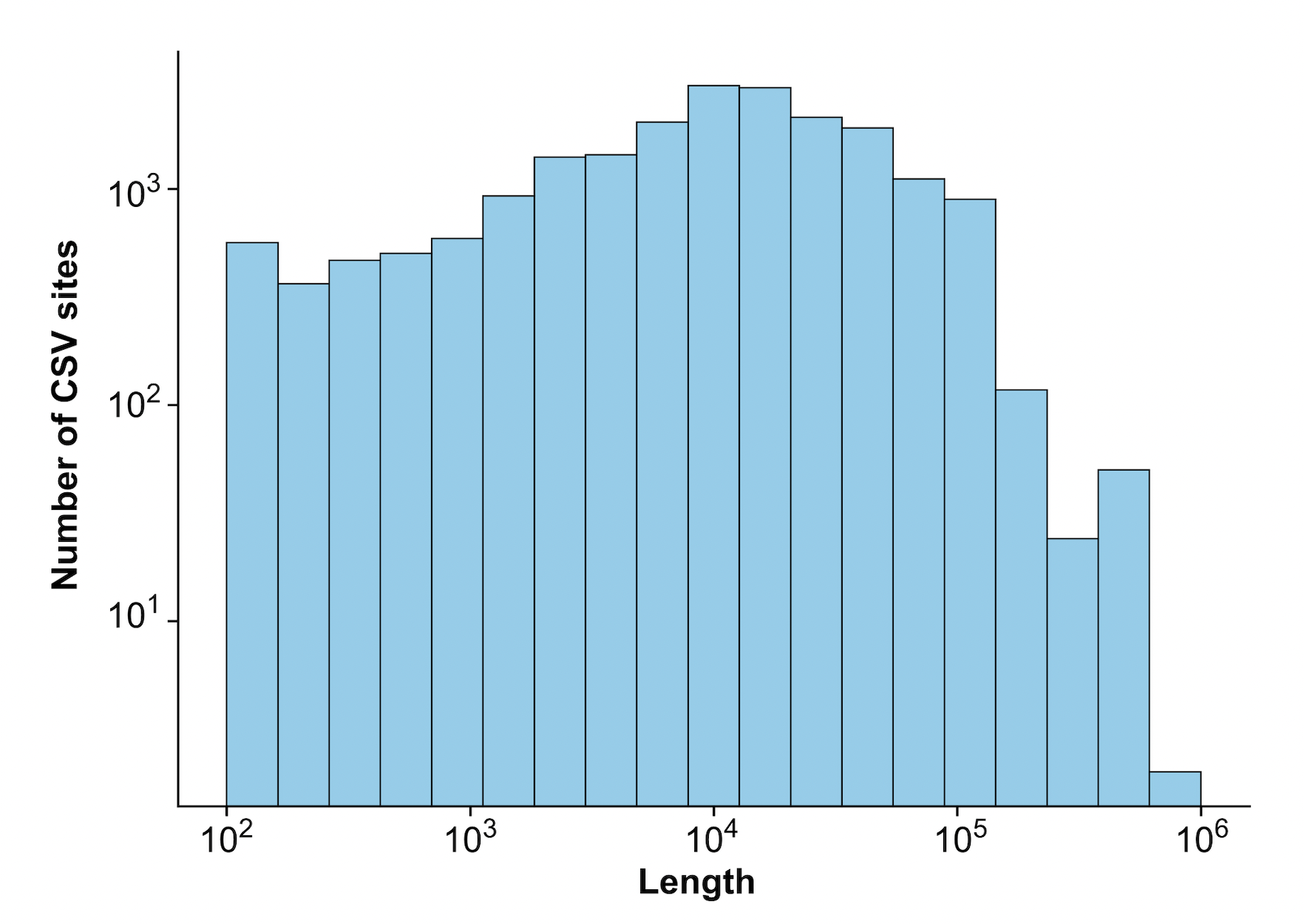

#### Supplementary Figure 19: Length distribution of CSV sites.

Histogram showcasing number of CSV sites (log scale y-axis) falling within each length interval (log scale x-axis).

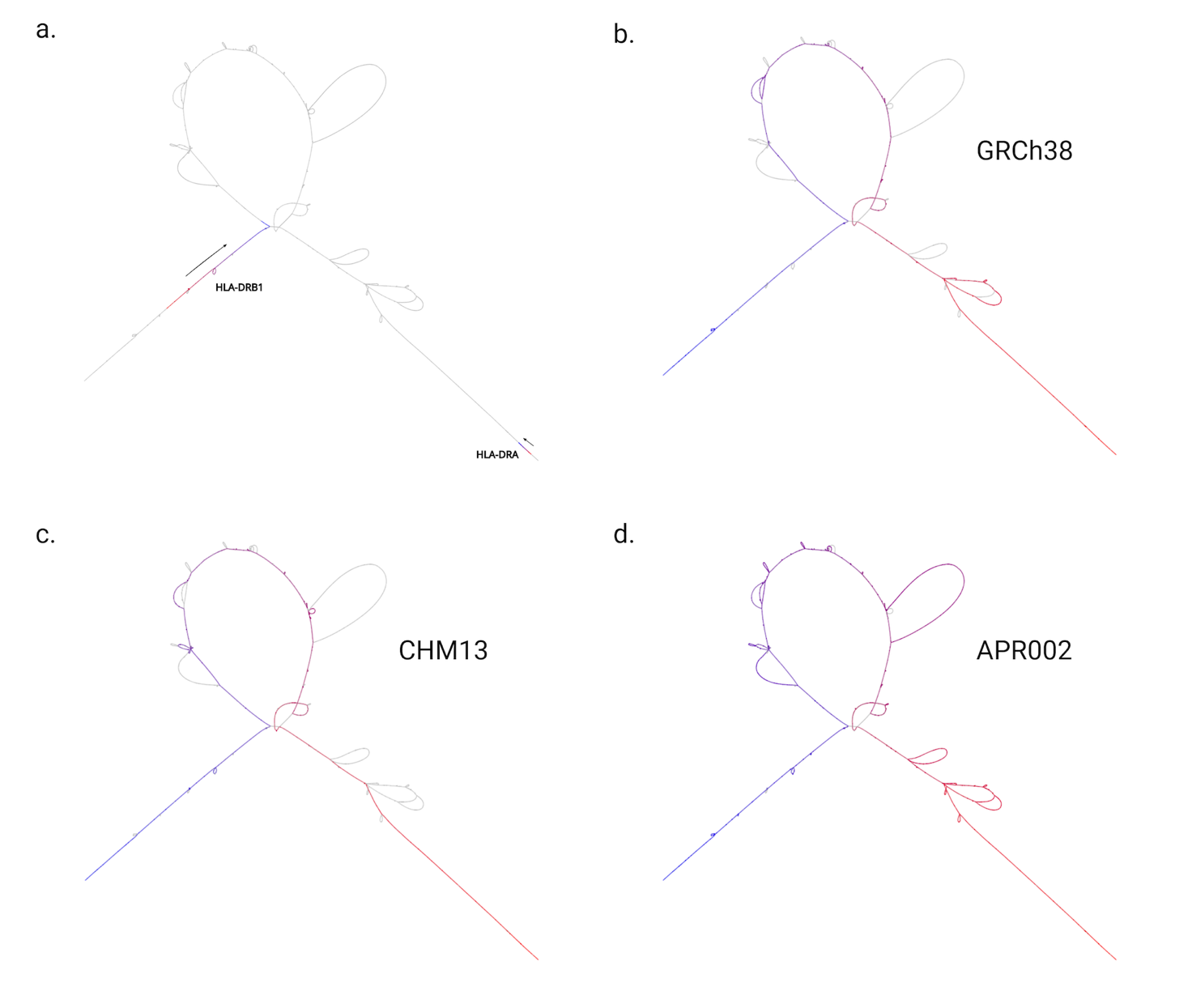

#### Supplementary Figure 20: Visualizing complex pangenome loci.

HLA region encompassing *HLA-DRB1* and *HLA-DRA* gene showing varying paths taken by APR sample which is absent in references (GRCh38 and CHM13). Variation among haplotype walks that did not involve genes was visualized using color coded lines, from red to blue to indicate directions.

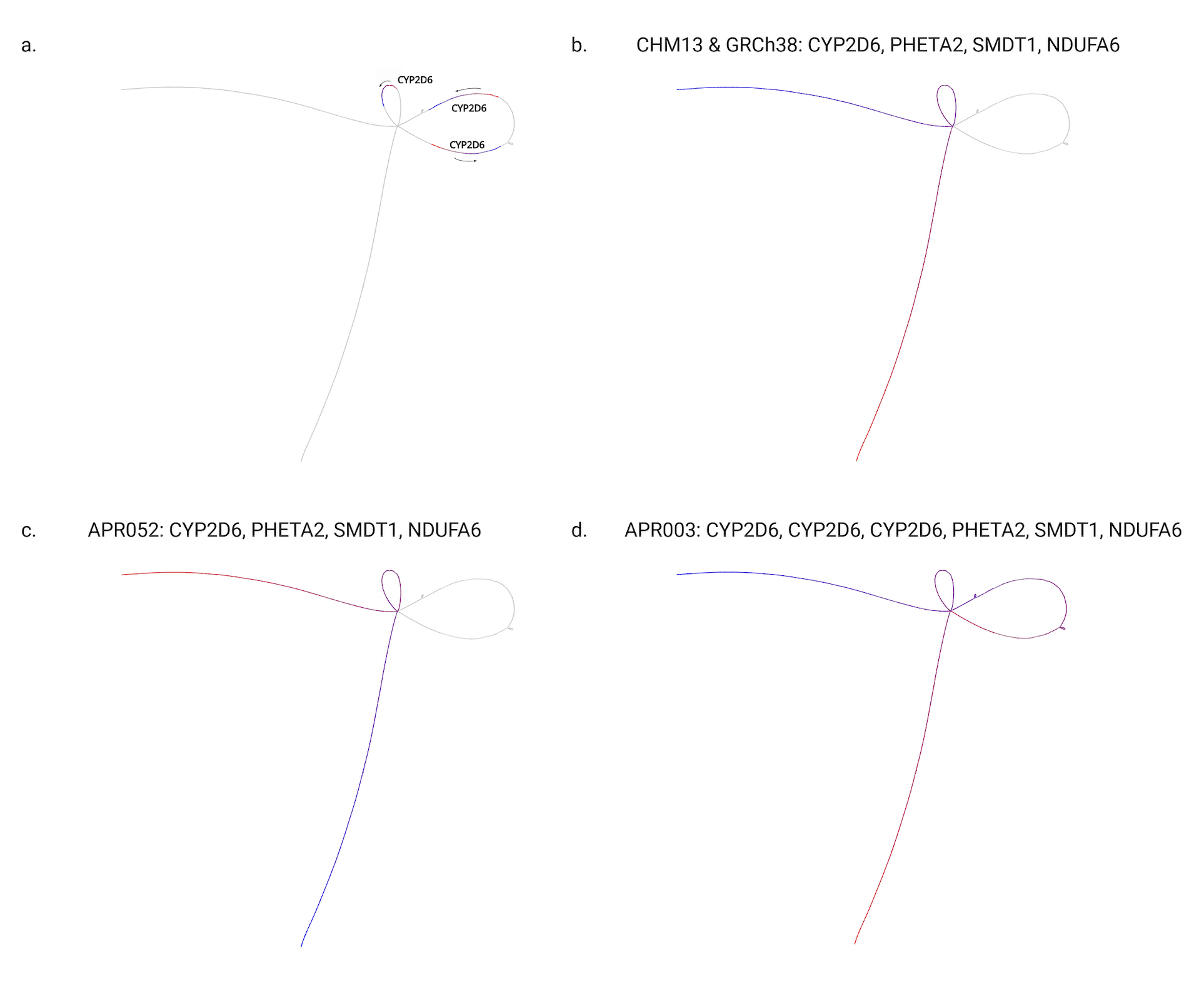

#### Supplementary Figure 21: Visualizing complex pangenome loci.

CYP2D6 region encompassing SVs showing absence of haplotypes in references, GRCh38 and CHM13. Inversions and insertions within CYP2D6 in APR samples are displayed. Variation among haplotype walks that did not involve genes was visualized using color coded lines, from red to blue to indicate directions.

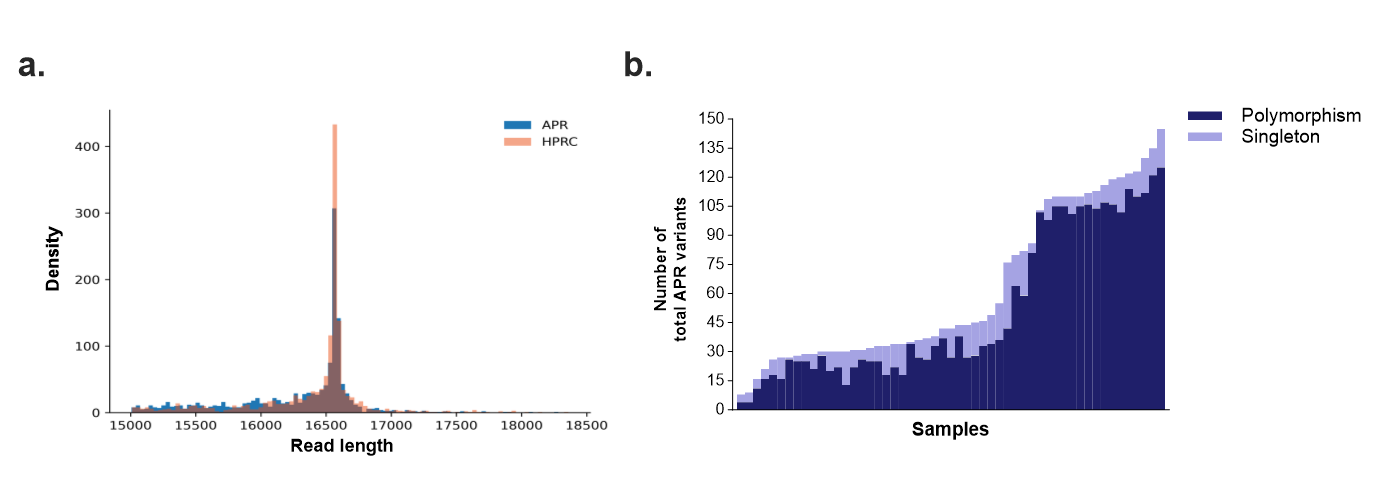

#### Supplementary Figure 22: Mitochondrial read and variant distribution.

**a.** Histogram representing density (y-axis) of read length (x-axis) distribution for mitochondrial reads in APR (blue) and HPRC (orange), where represents density. **b.** Bar chart representing the total number of small variants observed across different APR samples, differentiated between polymorphism (dark blue) and singleton (light blue).

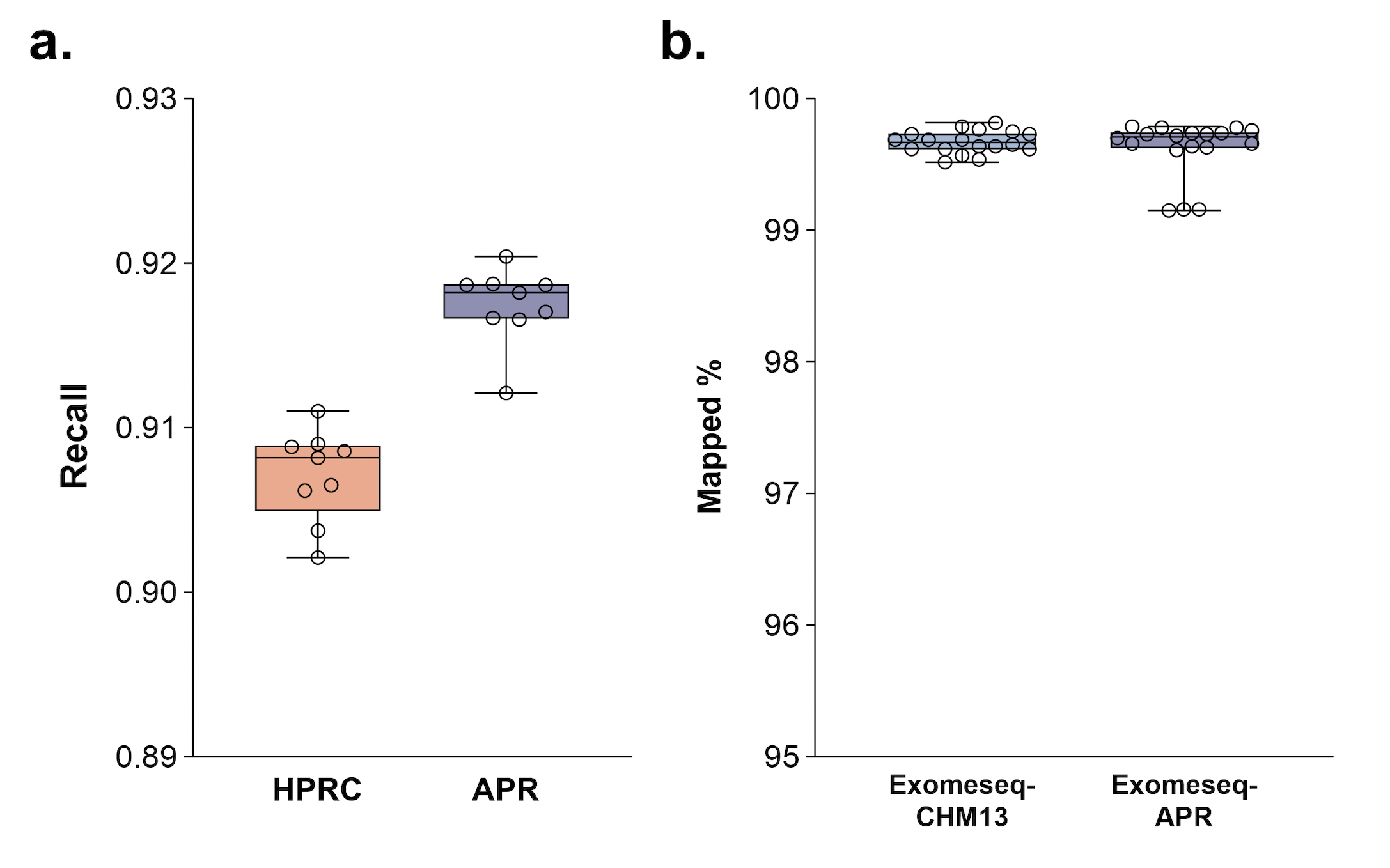

#### Supplementary Figure 23: Short read mapping performance gain.

**a.** Genotyping recall for SNPs. Box plot depicting the recall rates for genotyping polymorphic variants across all genomic region based on CHM13 variant calls. **b.** Mapping rate of whole exomeseq data from ASD trio samples to CHM13 and APR.

### References

1.	Sherry, S.T., Ward, M.H., Kholodov, M., Baker, J., Phan, L., Smigielski, E.M., and Sirotkin, K. (2001). dbSNP: the NCBI database of genetic variation. Nucleic Acids Res *29*, 308–311. https://doi.org/10.1093/nar/29.1.308.

2.	Karczewski, K.J., Francioli, L.C., Tiao, G., Cummings, B.B., Alföldi, J., Wang, Q., Collins, R.L., Laricchia, K.M., Ganna, A., Birnbaum, D.P., et al. (2020). The mutational constraint spectrum quantified from variation in 141,456 humans. Nature *581*, 434–443. https://doi.org/10.1038/s41586-020-2308-7.

3.	1000 Genomes Project Consortium, Auton, A., Brooks, L.D., Durbin, R.M., Garrison, E.P., Kang, H.M., Korbel, J.O., Marchini, J.L., McCarthy, S., McVean, G.A., et al. (2015). A global reference for human genetic variation. Nature *526*, 68–74. https://doi.org/10.1038/nature15393.

4.	Scott, E.M., Halees, A., Itan, Y., Spencer, E.G., He, Y., Azab, M.A., Gabriel, S.B., Belkadi, A., Boisson, B., Abel, L., et al. (2016). Characterization of Greater Middle Eastern genetic variation for enhanced disease gene discovery. Nat Genet *48*, 1071–1076. https://doi.org/10.1038/ng.3592.

5.	MacDonald, J.R., Ziman, R., Yuen, R.K.C., Feuk, L., and Scherer, S.W. (2014). The Database of Genomic Variants: a curated collection of structural variation in the human genome. Nucleic Acids Res *42*, D986-92. https://doi.org/10.1093/nar/gkt958.

6.	Cheng, H., Concepcion, G.T., Feng, X., Zhang, H., and Li, H. (2021). Haplotype-resolved de novo assembly using phased assembly graphs with hifiasm. Nat Methods *18*, 170–175. https://doi.org/10.1038/s41592-020-01056-5.

7.	Rautiainen, M., Nurk, S., Walenz, B.P., Logsdon, G.A., Porubsky, D., Rhie, A., Eichler, E.E., Phillippy, A.M., and Koren, S. (2023). Telomere-to-telomere assembly of diploid chromosomes with Verkko. Nat Biotechnol *41*, 1474–1482. https://doi.org/10.1038/s41587-023-01662-6.
